# The rise, decline and fall of clades

**DOI:** 10.1101/2025.03.20.644316

**Authors:** Ignacio Quintero, Jérémy Andréoletti, Daniele Silvestro, Hélène Morlon

## Abstract

How and why species diversity varies over geological time scales remains disputed. We analyzed the evolutionary history of 27 radiations with phylogenetic information incorporating extinct and extant species under a new ‘fossilized birth-death diffusion’ model that provides a detailed characterization of past diversification and resulting diversity dynamics. Clade diversity trajectories follow rise and decline dynamics, with fast accumulation following recurrent speciation while slowdowns and losses are modulated primarily by changes in extinction. Diversity dynamics do not appear to be governed by clade-level processes expected from adaptive radiations or diversity equilibria. Rather, these patterns emerge from dynamics at the species-level, where lineages tend to become increasingly vulnerable to extinction and less likely to speciate with time. Those species that counteract this trend create and maintain biodiversity through deep time.

**One Sentence Summary:** 

## Main Text

Despite its central importance to evolutionary theory, the question of how and why species diversity varies over geological time scales remains highly debated (*1, 2*). Key aspects of this debate include the importance of equilibrium versus non-equilibrium dynamics (*3, 4*), the preponderance of adaptive radiations (*5*), the temporal dynamics of ‘wax’ and ‘wane’ diversity phases for clades that rise and fall (*6, 7*), and the role of speciation versus extinction in driving these dynamics (*8, 9*). Extensive work in the field has led to substantial advancement in our understanding of past speciation and extinction dynamics using either phylogenetic data, the fossil record, or a combination of the two (*e.g*., (*9–17*)). Yet recovering diversity declines and extinction rates remains difficult, especially when using phylogenies of extant-only species (*18*), and for clades with scarce fossil record (*19, 20*).

Phylogenetic analyses of extant species typically suggest that diversity is currently expanding or at equilibrium, with very few exceptions (*9, 18, 21*). This has partly crystallized the debate around the existence and significance of ‘ecological limits’, *i.e*., the potential role of limited ecological opportunities in generating diversity-dependent dynamics and determining an equilibrium number of species within a clade (*3, 22*). This debate has also played a prominent role among paleobiologists (*23, 24*), together with controversy on how and why clades thrive and go extinct (*2, 6–8*). For instance, the temporal predictability, if any, of the rise and decline phases of diversity across clades remains contentious (*6–8, 25–27*). Under constant rates of species gains and losses, the duration of the rise and decline phases should be, on average, comparable, resulting in largely symmetric diversity trajectories (*8, 25, 27, 28*). However, a prevailing view holds that much of evolutionary history is driven by the colonization of preexisting ‘adaptive zones’ after the appearance of evolutionary innovations, extinction of prior occupants, or invasion of new geographic areas (*29–31*). Here, diversity should accumulate early and then slowly erode as speciation events no longer exceed extinctions, resulting in a ‘bottom-heavy’ trajectory (*6, 29, 32*). Alternatively, if clades are rapidly wiped out, then the decline phase could be shorter, resulting in a ‘top-heavy’ trajectory (*25*). Regardless of how diversity fluctuates, these variations are determined by the balance between speciation and extinction events, and one of the most enduring questions is the relative importance of these two processes (*8, 33, 34*). Specifically, periods of diversity expansion may correspond to times characterized by a surge in speciation or few extinctions, while periods of diversity decline may align with times marked by few speciations or many extinctions. It is unclear whether speciation or extinction predominantly drives diversity trends, and to what extent there are consistent patterns across clades.

We evaluated these questions by examining diversity and diversification dynamics of 27 taxonomic groups with species-level phylogenetic trees including fossils, under a new ‘fossilized birth-death diffusion’ (‘FBDD’) model (fig. S1). These evolutionary radiations represent a total of 5416 species, of which 3944 are extinct, spanning the walnut tree family, palms, sea scorpions, crocodiles, turtles, lung- and ray finned-fishes, non-avian dinosaurs, pterodactyls, penguins, therocephalians and several Cenozoic mammal clades (Materials and Methods, Table S1). Under the fossilized birth-death process, lineages speciate with rate *λ*, go extinct with rate *µ*, and leave fossils that are recovered with rate *ψ* (*14, 35*); this model is identifiable and allows to estimate diversification dynamics using information from phylogenetic and fossil data jointly (*36*). We expanded this model in two main directions (fig. S1; Materials and Methods). First, we introduced a fine-grained characterization of the diversification process by considering that lineage-specific speciation and extinction rates are inherited at speciation and evolve according to a Geometric Brownian motion, following the birth-death-diffusion model (*16*). Our assumption of heritability of speciation and extinction rates is supported by theoretical and empirical evidence (*16, 37*) and implicit in all other phylogenetic and paleontological models of diversification where rates are shared across taxonomic groups of species. We include two drift terms, *α*_*λ*_ and *α*_*µ*_, which capture overall trends in speciation and extinction through time, respectively, and two diffusion terms, *σ*_*λ*_ and *σ*_*µ*_, reflecting their variability. Explicitly incorporating the fossilization process allows our model to estimate lineage-specific speciation and extinction dynamics jointly using total-evidence, rather than to estimate extinction rates independently using the fossil record at the clade-level, as done in (*16*). Second, we developed and validated Bayesian data augmentation (DA) (*38*) techniques for fossilized birth-death models (figs. S2-S4). We accounted for temporal differences in fossilization rates by allowing piece-wise constant variation (*39*). As empirical trees typically include a single representative fossil occurrence per species, misrepresenting the expectations under a fossilized birth-death process (*40*), we allowed the addition of other known fossil occurrences of species in the tree (see Materials and Methods). Taking as input an empirical tree and, optionally, additional fossil occurrences for species in this tree, our data augmentation approach provides posterior samples of *complete trees*, which contain plausible unsampled extant and extinct taxa under the model and can thus directly be used to reconstruct diversity trajectories. It also provides estimates of the piece-wise constant fossilization rates and, in the case of the fossilized birth-death-diffusion model, of the trends *α*_*λ*_ and *α*_*µ*_, the diffusion term for speciation *σ*_*λ*_ and for extinction *σ*_*µ*_ (although this parameter should be interpreted with caution; fig. S4), as well as instantaneous speciation and extinction rates specific to all sampled and unsampled lineages.

Applied to the 27 taxonomic groups, our FBDD model recovered complex diversity trajectories with successive periods of increases and decreases (Fig. 1A & fig. S6) that would have been difficult to find using extant-only species (*18*). Focusing on the initial, highest and final diversity of these clades, we found 8 ‘on the rise’, with highest diversity at the present, including the new world monkeys (platyrrhini), penguins (spheniscidae) and turtles (testudinata); 14 ‘in decline’, with lower present than past diversity, including sphenodontids, ruminants and proboscideans, and 5 that went extinct, including sea scorpions (eutyperida), pterosaurs and brontotheriids (Materials and Methods; Fig. 1 & fig. S6). We find largely consistent reconstruction for groups for which past diversity has been previously estimated, such as cetaceans (*9, 41*) and proboscideans (*42*).

**Fig. 1:**
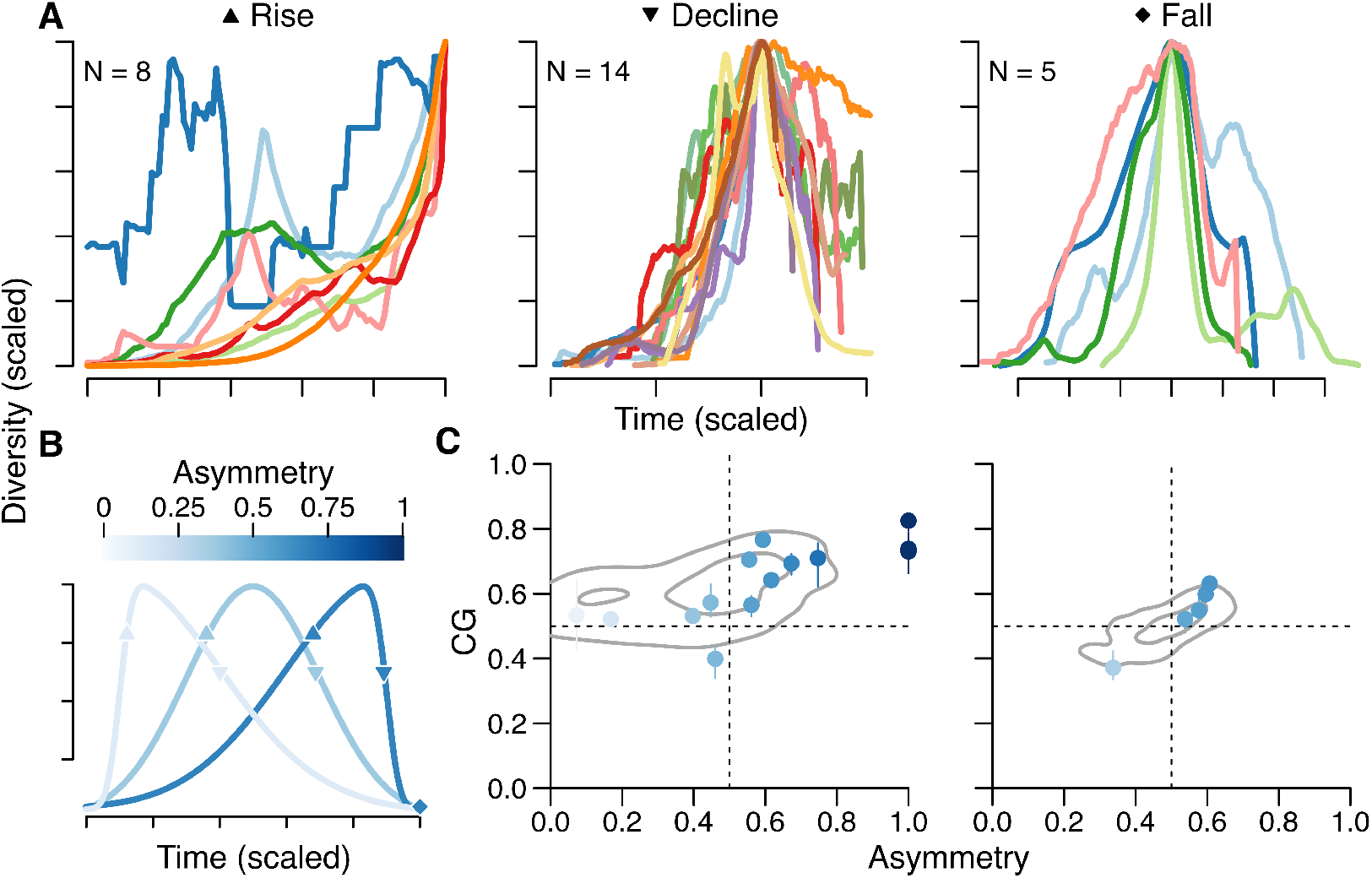
Rise, decline and fall of diversity show asymmetries for 27 taxonomic groups. **A)** Posterior average diversity trajectories estimated from our Bayesian data augmentation algorithm under the fossilized birth-death diffusion (FBDD) model using *complete trees* for each of the 27 taxonomic groups studied. Time and diversity are scaled, and diversity trajectories are overlaid with respect to their point of maximum diversity. Groups on the rise (*left*) have a higher diversity at the present, those on decline (*middle*) have a a lower present-day diversity, and those that fall (*right*) have gone extinct. **B)** Illustration of different asymmetries in diversity curves: from *<* 0.5 being ‘bottom-heavy’ (whiter colors), through ≈ 0.5 being ‘symmetrical’, to *>* 0.5 being ‘top-heavy’ (bluer colors). **C)** Respective asymmetries using the Center of Gravity (CG) and our parametric Asymmetry (see Main Text and Materials and Methods) for those clades in decline (*left*) and that fall (*right*). Grey contour lines denote, respectively, the 50% and 95% density of asymmetries and CG values for diversity trajectories resulting from simulations of clades on decline or extinct under a constant birth-death rate model with a species turnover of 1 per My. The three asymmetry values of 1 result from groups whose recent decline was not inferred by the parametric asymmetry model.

Diversity trajectories did not resemble the logistic curve expected under a single equilibrium diversity, or a succession of logistic curves with multiple equilibria (*7, 21, 25*) (Fig. 1A). The absence of equilibrium dynamics was confirmed by testing whether estimates of average speciation and extinction rates across lineages through time were associated with diversity: at the clade level, diversity had idiosyncratic effects on speciation of mostly negligible magnitude, with 11 radiations showing a negative and 8 a positive influence, and it marginally decreased extinction in 6 and increased extinction in 15 out of the 27 groups (fig. S8, Materials and Methods). In contrast to previous work (*4, 8, 43*), this resulted in an absence of support for an overall effect of diversity in either speciation (mean effect of diversity = −0.00012, understood as a 0.012% average percentage decrease in speciation rates by the addition of one species, 95% Highest Posterior Density (HPD) Intervals = [−0.14%, 0.094%]) or extinction rates (mean effect of diversity = 0.00052, understood as a 0.052% average percentage increase in extinction rates by the addition of one species, 95% Credible Intervals (CI) =[−0.23%, 0.41%], fig. S8, Materials and Methods).

For groups that are either extinct or in decline, we measured wax and wane symmetry using two correlated metrics that range between 0 and 1, have values around 0.5 when diversity trajectories are largely symmetrical, below 0.5 when they are bottom-heavy, and above 0.5 when they are top-heavy (Fig. 1B; Materials and Methods). Contrary to expectations under adaptive radiation theory (asymmetry *<* 0.5; (*6, 29*)) or rapid declining phases (asymmetry *>* 0.5; (*25*)), we find asymmetries that are mostly indistinguishable from those expected under a null model where speciation and extinction rates are constant and balance each other out (Fig. 1C & fig. S7; Materials and Methods).

Diversity expands when there is a positive net diversification (that is, when speciation exceeds extinction), and it contracts when there is a negative net diversification, but such changes can stem from fluctuations in either speciation, extinction, or both rates. We determine the relative prominence of speciation and extinction rates in driving changes in net diversification rates, and thereby in diversity (Materials and Methods). Their estimated importance, *I*_*s*_ for speciation and *I*_*e*_ for extinction, measures their relative role during changes in diversification, such that if, for instance, there is a linear increase in diversification rates of 0.2 stemming from an increase of speciation rates of 0.05 and a decrease in extinction rates of 0.15, then *I*_*s*_ = 0.25 and *I*_*e*_ = 0.75 (Fig. 2A-C; Materials and Methods). Because their importance might change during different diversity stages (Fig. 2), we measure *I*_*s*_ and *I*_*e*_ when i) there is an accelerating gain in diversity (*i.e*., when the net diversification rate is positive and increasing), ii) there is a decelerating gain in diversity (*i.e*., when the net diversification rate is positive and decreasing), iii) there is a decelerating loss of diversity (*i.e*., when the net diversification rate is negative but is increasing), and iv) there is an accelerating loss of diversity (*i.e*., when the net diversification rate is negative and decreasing). Combined across taxa, stage *i* sums up to about 778 Mys of evolutionary history (*ca*. 26 % of total), stage *ii* to about 1249 Mys (*ca*. 42 % of total), stage *iii* to about 351 Mys (*ca*. 12 % of total), and stage *iv* to about 582 Mys (*ca*. 20 % of total).

**Fig. 2:**
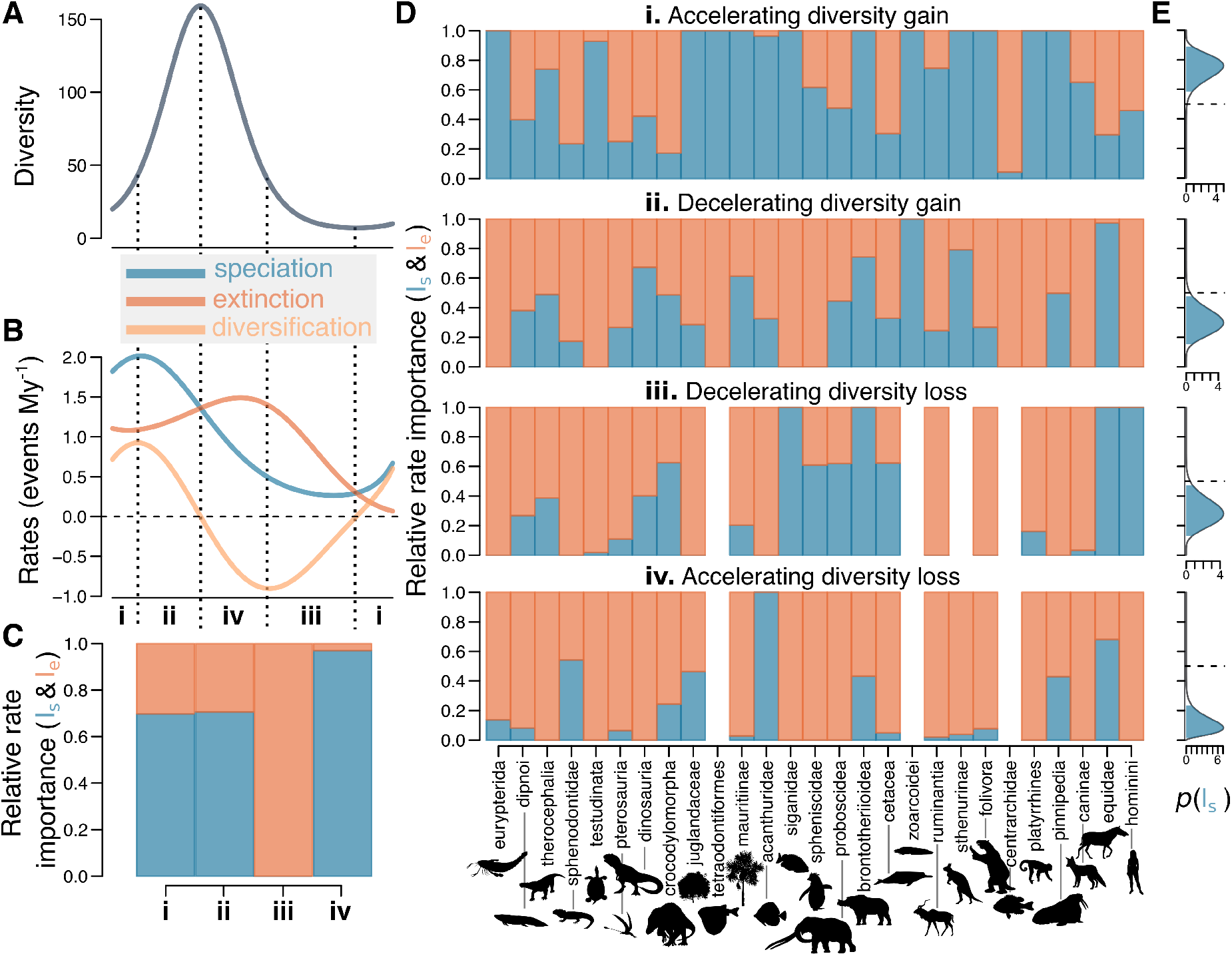
Relative importance of speciation versus extinction in driving diversification dynamics. **A)** An hypothetical diversity trajectory, determined completely by the underlying **B)** average rates of speciation (blue), extinction (red) and diversification (orange) rates, the latter being speciation minus extinction. **C)** The relative importance of speciation (‘*I*_*s*_’, in blue) versus extinction (‘*I*_*e*_’, in red) in driving the changes in diversification rates across four different diversity stages, **i-iv**, described in D (see Materials and Methods) and demarcated in the illustrative example of A & B. The higher the importance, the larger role speciation (or extinction) plays in driving the change in overall diversification. **D)** Empirical results of the relative rate importance across the 27 evolutionary radiations considered for each of the diversity stages (**i-iv**). Taxonomic groups are ordered by order of first appearance. **E)** The combined posterior probability of speciation being more responsible than extinction in driving diversification (and thus diversity) changes for each diversity stage across clades. The 95% Highest Posterior Density (HPD) interval is shown in blue.

We find that the importance of speciation and extinction in driving fluctuations in diversification rates varies across clades (Fig. 2D), but that overall, speciation is the dominating factor during periods of accelerated gain in diversity (average *I*_*s*_ 0.742, 95% HPD [0.58, 0.89], Fig. 2E). Variations in diversification are otherwise primarily related to changes in the intensity of extinctions: it is the case during periods of decelerating gain in diversity (average *I*_*e*_ 0.69, 95% HPD [0.52, 0.85]), accelerating diversity loss (average *I*_*e*_ 0.71, 95% HPD [0.53, 0.87]), and decelerating diversity loss (average *I*_*e*_ 0.88, 95% HPD [0.76, 0.986]; Fig. 2E). Hence, rapid accumulation of diversity results from an increase in speciation rates that together account for about *ca*. 0.26 % of groups’ evolutionary history, but its slowdown and depletion is modulated primarily by fluctuations in extinction dynamics. The buffering of clades’ decline (diversity stage **iii**) is primarily due to a decline in extinction rates, and can explain the frequency of clades persisting for a long time with low diversity (*44*).

A particular issue when using a phylogenetic approach is that the ‘depth’ along the tree of life at which to define focal clades is somewhat arbitrary, and that different phylogenetic scales can result in different evolutionary dynamics. For instance, including a non-diverse stem group to an otherwise diverse crown that radiated early can change the asymmetry in the diversity trajectory from *<* 0.5 to *>* 0.5. So far, the 27 taxonomic groups here studied, delineated by researchers on a number of synapomorphies *a priori*, have been treated as units, but their differences in number of tips, evolutionary age and morphological cohesiveness, among others, might make them unsuitable as comparable evolutionary units.

Because we are interested in how species richness rise and fall and their underlying diversification, irrespective of their morphological similarity and taxonomic level, we delineated mutually-exclusive monophyletic groups comprising between 20 to 40 tip species, ‘subclades’, across the posterior distribution of complete trees from the 27 taxonomic groups (Materials and Methods). The designation ‘subclade’ is relative and we use it here to emphasize their nested structure within the studied radiations. Although the number of species is arbitrary, the criterion itself leads to a greater equivalence and represents more comparable evolutionary units (*45*).

An analysis of diversity and diversification patterns in these subclades (Materials and Methods) reveal patterns that are mostly consistent with those at the higher level. Diversity is not necessarily accumulated at the end or beginning of a radiation (Fig. 3A). Rather, it undergoes idiosyncratic trajectories, even within the same clade (fig. S36-S37). Extinction remains the main driver during diversity loss (stages **iii** and **iv**; Fig. 3B) and speciation remains the main driver of enhanced diversity gain (diversity stage **i**). However, in contrast to the results at the higher level, it now plays a stronger role than extinction in decelerating diversity gain (diversity stage **ii**). This difference suggests that the overall trajectory of a clade in a “slow-down” phase results from the aggregation of concurrent radiations, some of which may be expanding, primarily driven by speciation, and others declining, driven primarily by higher extinction, with the latter overtaking the signal in the overall diversification trajectory of the clade.

**Fig. 3:**
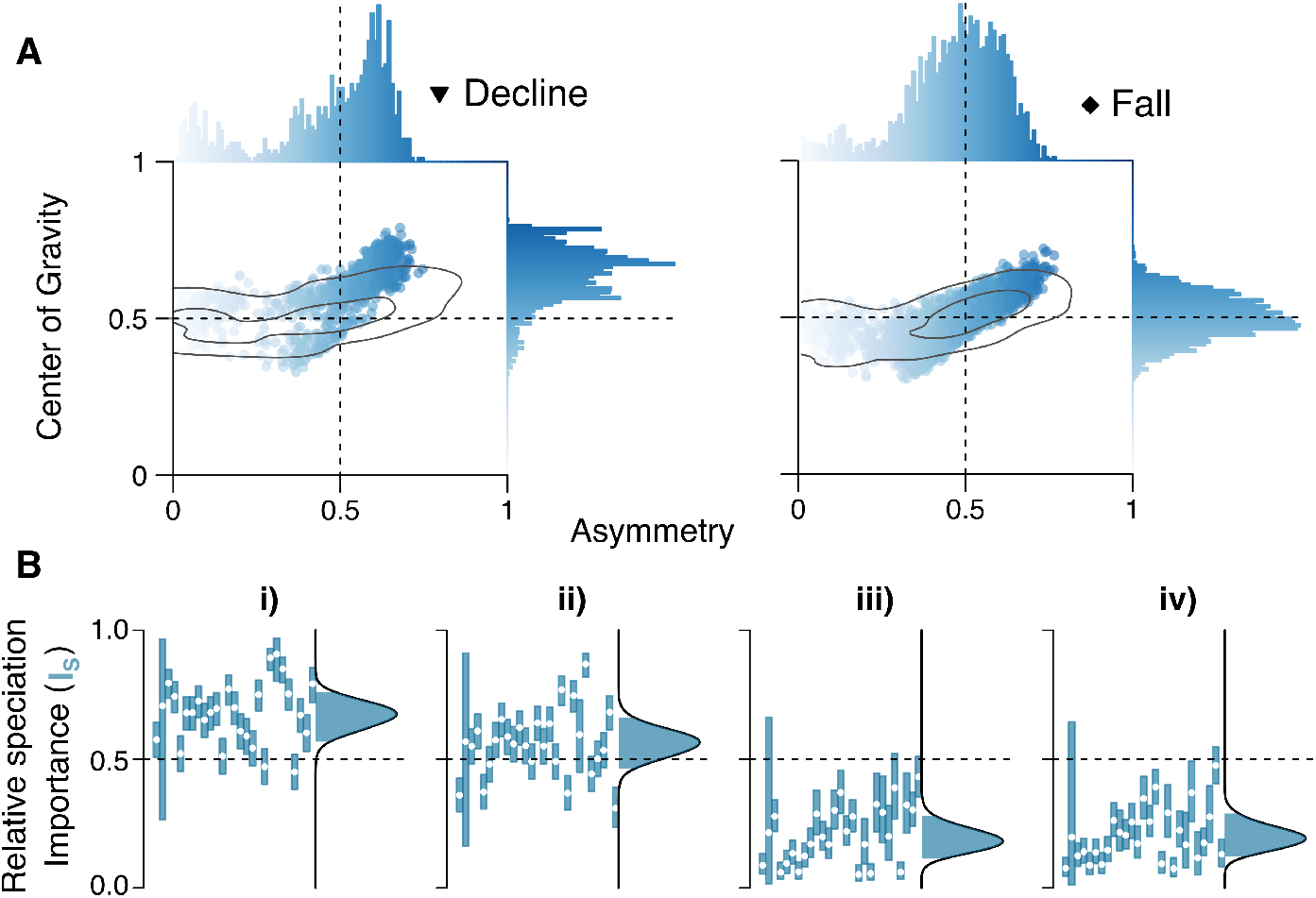
Diversity trajectories and their underlying diversification dynamics across subclades. **A)** Center of Gravity (CG) and Asymmetry measures for all the subclades of 20 − 40 tips considered in decline (*left*) and that fall (*right*) across the 27 taxonomic groups. Points show the multivariate distribution between both metrics and their respective marginal distributions are shown on the edges. Color gradient follows Fig. 1B. Grey contour lines denote, respectively, the 50% and 95% density of asymmetries and CG values for diversity trajectories resulting from simulations of clades on decline or extinct under a constant birth-death rate model with a species turnover of 1 per My. **B)** The relative importance of speciation, *I*_*s*_, in driving diversification and diversity dynamics during each of the four diversity stages, **i-iv** (see Fig. 2E), as given by the hierarchical model detailed in the Main Text and Material and Methods. For each diversity stage, the vertical bars to the left represent the mean (white dot) and 95% Credible Interval (CI) for the posterior probability of the importance of speciation across the subclades for each of the 27 groups; the distribution to the right is the combined posterior probability distribution for the importance of speciation (black thick line shows the median parameters of this distribution, with its 95% Highest Posterior Density (HPD) interval in blue; gray lines show the posterior variance of this distribution).

Underlying these aggregated patterns, there is substantial lineage-level variation in diversification rates within evolutionary radiations (figs. S9-S35). We find a larger heterogeneity in speciation rates (posterior *σ*_*λ*_ median and range across taxa: 0.13, [0.08, 0.18]; fig. S38) than previous analyses using extant-only trees (*16, 46*). The level of heterogeneity in extinction rates appears comparable to that of speciation rates (posterior *σ*_*µ*_ median and range across taxa: 0.16, [0.13, 0.36]; fig. S38). We find a widespread tendency for lineage-specific speciation rates to decline by 3.8% every My (cross-clade mean *α*_*λ*_ = −0.038, 95% CI = [−0.054, −0.024]; Fig. 4 & fig. S38), consistent with previous results (*16, 46*). Lineage-specific extinction rates concomitantly tend to increase by 8.5% every My (cross-clade mean *α*_*µ*_ = 0.085, 95% CI = [0.056, 0.11]; Fig. 4 & fig. S38), contrary to expectations under the ‘Red Queen’ model, which predicts that extinction probabilities are independent or negatively correlated to age (*47*).

**Fig. 4:**
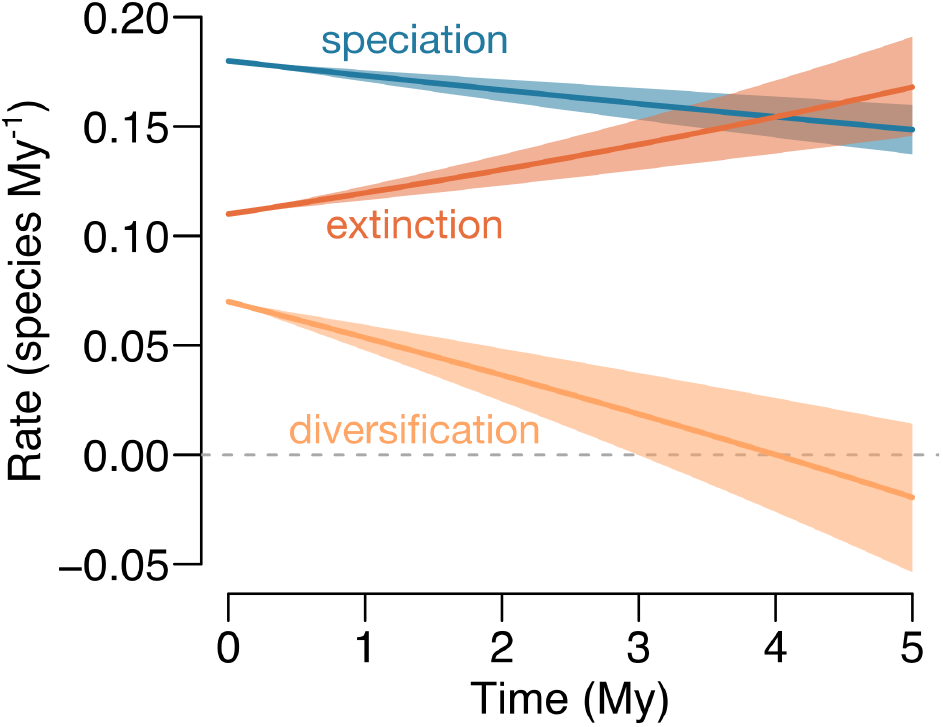
The insidious loss of species macroevolutionary fitness. Expected change in speciation (blue), extinction (red) and net diversification (speciation - extinction, in yellow) in time for a lineage starting with a speciation rate of 0.18 and an extinction rate of 0.11 species per My^−1^ as given by the cross-taxon posterior estimates for *α*_*λ*_ and *α*_*µ*_. Shaded areas represent the 95% Credible Intervals.

Most species that have ever lived on Earth are now extinct, and extinction therefore unsurprisingly plays a pivotal role in main macroevolutionary theories. Combining data from both extant and extinct species in a model that accounts for lineage-specific speciation and extinction rates allows us to test a series of long-standing questions about extinction that have otherwise been notoriously hard to examine. Across 27 evolutionary radiations spanning a variety of taxa, our results reveal symmetric diversity trajectories, which are mostly driven by changes in extinction rather than speciation dynamics, except during periods of accelerated diversity gain. Moreover, variations in extinction rates are not clade-wide, mirroring speciation dynamics (*16*).

As lineages age, their propensity to speciate declines, and they are more likely to go extinct (Fig 4). This can be explained if most evolutionary change occurs at speciation (*48–50*). Since species struggle to keep up with their changing biotic and abiotic environment (*51,52*), a paucity of speciation progressively leads to an evolutionary disadvantage with respect to an unstable environment and to those undergoing speciation (and changing) (*53*). We call this potential mechanism for both a decrease in speciation rates and an increase in extinction risk, the ‘Zarathustran hypothesis’, in reference to F. Nietzsche’s idea that the maintenance of life does not merely follow from conservation, but rather from constant renovation, movement, and expansion: “And life itself confided the secret to me: behold, it said, I am that which must always overcome itself.” (*54*). The few lineages that withstand this loss in diversification rates generate a strong imbalance, wherein only one of the descendants retains high rates of speciation and low rates of extinction along consecutive nodes (figs. S9-S35).

Like all historical disciplines, macroevolution suffers from hindsight bias (*55*): in retrospect, it seems that the evolutionary dynamics of clades were determined to be so. Rather, our results highlight the importance of non-equilibrium and imbalanced diversification dynamics in generating diversity, in disagreement with radiations along a limited set of ‘adaptive zones’ (*29*). We venture that stagnation brought about by a lack of speciation causes most lineages to lag behind a continuously reshaping environment. The few successful individual lineages that depart from this general tendency disproportionally engender biodiversity, yet sometimes fail to compensate a general insidious loss of macroevolutionary fitness, leading to the recurrent decline and fall of clades.

## Acknowledgments

We thank Nicolas Lartillot and the Morlon lab in general for early discussion of these ideas. D.S. received funding from ETH Zurich, from the Swiss National Science Foundation (PCEFP3 187012), and the Swedish Foundation for Strategic Environmental Research MISTRA within the framework of the research programme BIOPATH (F 2022/1448).

## Notes

### Competing Interest Statement

The authors have declared no competing interest.

## References

1. D. M. Raup, Science 231, 1528 (1986).

2. M. J. Benton, Science 323, 728 (2009).

3. L. J. Harmon, S. Harrison, The American Naturalist 185, 584 (2015).

4. D. L. Rabosky, A. H. Hurlbert, The American Naturalist 185, 572 (2015).

5. S. Gavrilets, J. B. Losos, Science 323, 732 (2009).

6. S. J. Gould, N. L. Gilinsky, R. Z. German, Science 236, 1437 (1987).

7. I. Žliobaitė, M. Fortelius, N. C. Stenseth, Nature 552, 92 (2017).

8. T. B. Quental, C. R. Marshall, Science 341, 290 (2013).

9. H. Morlon, T. L. Parsons, J. B. Plotkin, Proceedings of the National Academy of Sciences 108, 16327 (2011).

10. M. E. Alfaro, et al., Proceedings of the National Academy of Sciences 106, 13410 (2009).

11. T. Stadler, Proceedings of the National Academy of Sciences 108, 6187 (2011).

12. R. S. Etienne, B. Haegeman, The American Naturalist 180, E75 (2012).

13. D. L. Rabosky, PloS one 9, e89543 (2014).

14. T. A. Heath, J. P. Huelsenbeck, T. Stadler, Proceedings of the National Academy of Sciences 111, E2957 (2014).

15. A. Gavryushkina, et al., Systematic biology 66, 57 (2017).

16. I. Quintero, N. Lartillot, H. Morlon, Science 384, 1007 (2024).

17. T. Hauffe, J. L. Cantalapiedra, D. Silvestro, Science Advances 10, eadl2643 (2024).

18. G. Burin, L. R. Alencar, J. Chang, M. E. Alfaro, T. B. Quental, Systematic Biology 68, 47 (2019).

19. D. Silvestro, R. C. Warnock, A. Gavryushkina, T. Stadler, Nature Communications 9, 5237 (2018).

20. R. C. Warnock, T. A. Heath, T. Stadler, Paleobiology 46, 137 (2020).

21. O. Billaud, D. Moen, T. L. Parsons, H. Morlon, Systematic Biology 69, 363 (2020).

22. D. L. Rabosky, Ecology letters 12, 735 (2009).

23. J. J. Sepkoski, Paleobiology 19, 43 (1993).

24. J. Alroy, Proceedings of the National Academy of Sciences 105, 11536 (2008).

25. M. Foote, Paleobiology 33, 517 (2007).

26. H. Morlon, M. D. Potts, J. B. Plotkin, PLoS biology 8, e1000493 (2010).

27. N. Hohmann, E. Jarochowska, PeerJ 7, e8011 (2019).

28. S. Nee, Annu. Rev. Ecol. Evol. Syst. 37, 1 (2006).

29. G. G. Simpson, Tempo and mode in evolution (Columbia University Press, 1953).

30. D. Schluter, The ecology of adaptive radiations (Oxford: Oxford University Press, 2000).

31. C. Calderón del Cid, et al., Journal of Evolutionary Biology 37, 290 (2024).

32. M. Hughes, S. Gerber, M. A. Wills, Proceedings of the National Academy of Sciences 110, 13875 (2013).

33. D. M. Raup, J. J. Sepkoski, Science 215, 1501 (1982).

34. R. K. Bambach, A. H. Knoll, S. C. Wang, Paleobiology 30, 522 (2004).

35. T. Stadler, Journal of theoretical biology 267, 396 (2010).

36. K. Truman, T. G. Vaughan, A. Gavryushkin, A. S. Gavryushkina, Systematic Biology p. syae058 (2024).

37. D. Jablonski, Science 238, 360 (1987).

38. M. A. Tanner, W. H. Wong, Journal of the American Statistical Association 82, 528 (1987).

39. S. M. Holland, Philosophical Transactions of the Royal Society B: Biological Sciences 371, 20150130 (2016).

40. W. Pett, T. A. Heath, Phylogenetics in the Genomic Era, C. Scornavacca, F. Delsuc, N. Galtier, eds. (No commercial publisher — Authors open access book, 2020), pp. 5.1:1–5.1:18.

41. J. Andréoletti, et al., Systematic Biology 71, 1440 (2022).

42. R. B. Cooper, J. T. Flannery-Sutherland, D. Silvestro, Nature Communications 15, 4199 (2024).

43. M. Foote, The American Naturalist 201, 680 (2023).

44. B. D. Barnes, J. A. Sclafani, A. Zaffos, Proceedings of the National Academy of Sciences 118, e2019208118 (2021).

45. S. J. Gould, D. M. Raup, J. J. Sepkoski, T. J. Schopf, D. S. Simberloff, Paleobiology 3, 23 (1977).

46. O. Maliet, F. Hartig, H. Morlon, Nature ecology & evolution 3, 1086 (2019).

47. L. Van Valen, The American Naturalist 111, 809 (1977).

48. N. Eldredge, S. J. Gould, Models in paleobiology, T. J. M. Schopf, ed. (Freeman Cooper & Co, 1972), vol. 1972, pp. 82–115.

49. G. Hunt, Proceedings of the National Academy of Sciences 104, 18404 (2007).

50. O. Sanisidro, M. C. Mihlbachler, J. L. Cantalapiedra, Science 380, 616 (2023).

51. L. Van Valen, Evolutionary Theory 1, 1 (1973).

52. A. Spiridonov, S. Lovejoy, Nature 607, 307 (2022).

53. P. N. Pearson, Historical Biology 10, 119 (1995).

54. F. Nietzsche, Thus Spoke Zarathustra: A Book for All and None. Translated by Walter Kaufmann. Modern Library Edition (Random House, New York, 1995).

55. B. Fischhoff, Journal of Experimental Psychology: Human perception and performance 1, 288 (1975).

