## Supplementary Information for "The rise, decline and fall of clades"

### Materials and Methods

#### Fossilized Birth-Death Diffusion model

We assume that, at some time $t$, each lineage $l$ has an instantaneous rate of producing new species of $\lambda_l(t)$ (*i.e.*, speciation rate), an instantaneous rate of going extinct of $\mu_l(t)$ (*i.e.*, extinction rate), and an instantaneous fossilization rate of $\psi_l(t)$ (*1*). This general fossilized birth-death (FBD) process generates a bifurcating phylogenetic tree, $\Psi$, starting at some time $T_\Psi$ in the past and continues to time $T_0$ (not necessarily the present), with three separate events: the origination of a new lineage, the extinction of an existing lineage, and the fossilization of a lineage's individual that was recovered, sampled and incorporated in the tree. Furthermore, each clade $c$ has a specific probability for extant lineages to be represented in the reconstructed tree, specified by $\rho_c \in [0, 1]$.

Here, we introduce the Fossilized Birth-Death Diffusion model (FBDD), which expands on the Birth-Death Diffusion model (BDD; (*2*)) to incorporate fossils either as tips, when no further descendant has been sampled (*i.e.*, 'tip fossil') or within a lineage with sampled descendants (*i.e.,* 'sampled ancestor') in the reconstructed tree (fig. S1). We assume that $\lambda_l(t)$ and $\mu_l(t)$ each evolve anagenetically according to Geometric Brownian motion (GBM) processes, such that

$$d\ln(\lambda_l(t)) = \alpha_\lambda dt + \sigma_\lambda dW(t) \quad (1)$$

$$d\ln(\mu_l(t)) = \alpha_\mu dt + \sigma_\mu dW(t), \quad (2)$$

where $\alpha_\lambda$ and $\alpha_\mu$ represent the overall trend in rates of speciation and extinction, respectively (*i.e.*, the drifts), $\sigma_\lambda$ and $\sigma_\mu$ represent the accumulation of their respective rate variance (*i.e.*,, the diffusion coefficients) and $W(t)$ is the Wiener process (*i.e.*, the standard Brownian motion). The fossilization rate does not vary across contemporary lineages but we allow for it to by temporarily heterogeneous following a piecewise constant function across $K$ predefined periods. Specifically, we estimate different fossilization rates delimited by $|k| - 1$ times, $t^\psi$, such that $t^\psi_k \in [T_\Psi, T_0]$ where rates become $\psi_l(t) = \psi_k$, for $t^\psi_k < t < t^\psi_{k+1}$. This accommodates differential rates of fossil preservation across stratigraphic ranges (*3, 4*).

At time $T_\Psi$, the FBDD process starts with values $\lambda_l(T_\Psi) = \lambda_0$ and $\mu_l(T_\Psi) = \mu_0$ and $\psi_1$. If a lineage $a$ undergoes speciation at time $t_s$, yielding two daughters, $d_1$ and $d_2$, rates of speciation and extinction (and fossilization) are inherited identically, such that $\lambda_a(t_s) = \lambda_{d_1}(t_s) = \lambda_{d_2}(t_s)$ and $\mu_a(t_s) = \mu_{d_1}(t_s) = \mu_{d_2}(t_s)$. We note that, to our knowledge, all other diversification models assume stronger inheritance (except the nested BDD model (*2*)), either within different diversification regimes (*e.g.*, (*5, 6*)) or within time bins (*e.g.*, (*4, 7*)).

**Bayesian data augmentation (DA)** Because there is no analytical solution available for the likelihood under the FBDD model, we resort to Bayesian data augmentation (DA) techniques, building upon (*2*). This algorithm 'augments' the data such that the evaluation of the likelihood given the augmented data becomes tractable, leading to an exact estimation of the parameters' posterior distribution using Markov Chain Monte Carlo (MCMC) under the Metropolis-Hastings (MH) algorithm (*8, 9*). For the FBDD this entails the augmentation of unobserved lineages, including those that went extinct or were not sampled at the present as well as the lineage continuation (and descendants) from tip fossils, and lineage-specific speciation and extinction rates at discrete time intervals (the GBM diffusion paths at discrete time intervals, fig. S1). Performing inference using DA on the FBDD model yields a posterior distribution of *complete trees*, that is, trees that include both extant and extinct species and their associated speciation and extinction rates estimates. Let $\Psi_o$ represent the reconstructed (observed) tree, $\Psi_a$ the unobserved speciation and extinction events as well as the whole GBM discrete paths, and $\Psi$ ($= \Psi_o \cup \Psi_a$) the complete tree. We are able to estimate the desired posterior density $p(\lambda_0, \mu_0, \alpha_\lambda, \alpha_\mu, \sigma_\lambda, \sigma_\mu, \psi|\Psi_o)$ (*i.e.*, the probability of the set of model parameters, given the empirical tree $\Psi_o$), by treating $\Psi_a$ as a random variable to integrate over using MCMC. Specifically, $p(\lambda_0, \mu_0, \alpha_\lambda, \alpha_\mu, \sigma_\lambda, \sigma_\mu, \psi|\Psi_o) = \int_{\Psi_A} p(\lambda_0, \mu_0, \alpha, \sigma_\lambda, \sigma_\mu, \psi|\Psi_o, \Psi_a)p(\Psi_a|\Psi_o)d\Psi_a$, where $\Psi_A$ represent all the possible augmented histories given the empirical tree under the FBDD model.

We first developed DA algorithms for the constant fossilized birth-death model (CFBD) and an expanded version with piecewise constant fossilization rates (episodic fossilized birth-death model 'eFBD'), where speciation and extinction rates are constant and shared across lineages (*i.e.*, $\lambda_l(t) = \lambda$ and $\mu_l(t) = \mu$) and validated our approach (Supplementary Text). Then we developed a DA algorithm to make inference under the FBDD that expands on the one previously developed for the BDD model (*2*), so here we highlight the differences and touch briefly on previous derivations.

**Probability of a complete tree** To approximate the likelihood of a complete tree (*i.e.*, $\Psi$) under the FBDD model, we sample unobserved discrete GBM paths across complete trees. We set a sufficiently small time step $\delta$, such that $\delta = t_{i+1} - t_i$, where $t_i < t_{i+1}$, and divide each branch of the tree into $m$ small time steps such that $m = \lfloor t_b/\delta \rfloor + 1$, where $t_b$ represents the edge length of branch $b$. For a branch $b$ starting at $t_s$ and ending at $t_f$ we sample the data augmented diffusion process at times $t = \{t_1 = t_s, t_2 = t_1 + \delta, \ldots, t_{m+1} = t_f\}$, obtaining the stochastic evolution of speciation $\Lambda_b = \{\lambda_b(t_1), \ldots, \lambda_b(t_{m+1})\}$ and of extinction $\mathrm{M}_b = \{\mu_b(t_1), \ldots, \mu_b(t_{m+1})\}$ in a discrete time grid. For clarity and conciseness we denote $\lambda_l(t_i)$ as $\lambda_i$ (and thus $\lambda_l(t_b) = \lambda_b$) and $\mu_l(t_i)$ as $\mu_i$ from now on.

The likelihood that no event happens during time $\delta_i = t_{i+1} - t_i$ and that the rates starting at $\lambda_i$ and

$\mu_i$ end up at $\lambda_{i+1}$ and $\mu_{i+1}$ can be approximated as:

$$p(\text{no event in } [t_i, t_{i+1}], \lambda_{i+1}, \mu_{i+1} | \lambda_i, \mu_i, \psi_k, \alpha_\lambda, \alpha_\mu, \sigma_\lambda, \sigma_\mu) =$$
$$\exp\left\{-(\bar{\lambda}_{i,i+1} + \bar{\mu}_{i,i+1} + \psi_k)\delta\right\} \times \frac{1}{\sigma_\lambda\sqrt{2\pi\delta}} \exp\left\{-\frac{(\Delta\lambda_i - \alpha_\lambda\delta)^2}{2\delta\sigma_\lambda^2}\right\} \times \frac{1}{\sigma_\mu\sqrt{2\pi\delta}} \exp\left\{-\frac{(\Delta\mu_i - \alpha_\mu\delta)^2}{2\delta\sigma_\mu^2}\right\},$$

where $\bar{\lambda}_{i,i+1}$ and $\bar{\mu}_{i,i+1}$ represent the geometric mean for speciation and extinction at times $t_i$ and $t_{i+1}$ and $\Delta\lambda_i$ and $\Delta\mu_i$ are differences in logarithmic scale for speciation and extinction. The combined likelihood for the complete tree with speciation and extinction rates along the tree, $\Lambda$ and M respectively, under the FBDD process becomes

$$p(\Psi | \Lambda, \mathrm{M}, \lambda_0, \mu_0, \psi, \alpha_\lambda, \alpha_\mu, \sigma_\lambda, \sigma_\mu) =$$
$$\prod_{k \in K} f_k \psi_k \prod_{b \in \Psi_\mathrm{I}} \lambda_b \prod_{b \in \Psi_{\mathrm{T}_\mu}} \mu_b \prod_{b \in \Psi} \prod_{i=1}^{m} p(\text{no event in } [t_i, t_{i+1}], \lambda_{i+1}, \mu_{i+1} | \lambda_i, \mu_i, \alpha_\lambda, \alpha_\mu, \psi_k, \sigma_\lambda, \sigma_\mu), \tag{3}$$

where $f_k$ are the number of sampled fossils in period $k$, $\Psi_\mathrm{I}$ are the set of internal branches and $\Psi_{\mathrm{T}_\mu}$ are the set of terminal extinct branches. The conditioning on survival of the process is specified according to either the survival of both crown lineages, the survival of only one of them, or no survival, using a pseudo-marginal approach following (*2*). If not all extant species have been sampled (any $\rho_c < 1$), we follow (*2, 10*) to allow for heterogeneous incomplete sampling. Briefly, we specify each tip in the tree with their specific sampling fraction and propagate this fraction towards the roots of the tree multiplicatively. We then accept proposals for non-sampled species according to a binomial distribution following their branch-specific sampling fraction.

Finally, we use $K$ gamma priors for $\psi_k$, log-normal conjugate priors for the starting root rates $\lambda_0$ and $\mu_0$, normal conjugate priors for $\alpha_\lambda$ and $\alpha_\mu$, of mean $\nu$ and variance $\tau^2$, an inverse gamma for $\sigma_\lambda^2$, of shape $\kappa$ and scale $\zeta$, an inverse gamma for $\sigma_\mu^2$, of shape $\eta$ and scale $\xi$ and a uniform prior over $(0, 100)$ for $\lambda_0$ and $\mu_0$. The priors for $\lambda_0$, $\mu_0$, $\alpha_\lambda$, $\sigma_\lambda^2$ and $\sigma_\mu^2$ are conjugate with the augmented likelihood as shown in (*2*), and $\alpha_\mu$ follows the same derivation as and $\alpha_\lambda$ but for extinction rates. For $\psi_k$ we also use the conjugate Gamma prior, $\Gamma(\kappa, \varsigma)$ which results in the following full conditional distribution from which we can sample directly

$$p(\psi_k|\cdot) = \Gamma(\kappa + f_k, \varsigma + L_k),$$

where $L_k$ is the sum of branch lengths in the complete tree in period $k$.

**Data augmentation updates** We integrate over possible data augmented histories using diffusion updates and forward simulation. We use the same diffusion path updates described in (*2*) based on sampling internal nodes conditioned on their ancestor and descendants. For an internal node denoting a speciation event, this entails one ancestor and two daughters, and for a sampled ancestor node, this entails one ancestor and one daughter. Similarly, forward simulation proposals follows (*2*), but, evidently, simulating from the FBDD model and with the following peculiarities. First, simulated proposals are rejected when a fossilization event occurs (since we only observe those fossils that were effectively sampled). Second, we need new simulation proposals for a branch that ends in a sampled ancestor or in a tip fossil. For a sampled ancestor, we simulate the process with the same initial values as the ancestor during the branch length, and if there is more than one descendant at the time of the sampled ancestor, we pick one of them uniformly and carry on the simulation with the other descendants. For a tip fossil, we simulate in the same fashion during the branch leading to the fossil, but then carry on the simulation after the tip fossil. As in other forward simulations, the number of surviving species in the simulated trees must comply with the sampling fraction of the clade (*2*). Metropolis-Hasting ratios for these new forward simulation plots are found in the Supplementary Text.

The process parameters $\alpha_\lambda$, $\alpha_\mu$, $\sigma_\lambda$, $\sigma_\mu$ and $\psi_k$ are updated by Gibbs sampling from conjugate priors, as described in (*2*). However, note that if a clade survives to the present, we condition on survival, which involves an additional MH step using the pseudomarginal approach of (*2*).

**Validation and implementation** We validated our inference approach for CFBD by comparing it with the analytical solution and, together with FBDD, using extensive simulations detailed in the Supplementary Text, describing simulations, model statistical coverage and accuracy (figs. S2-S4). We implemented all the new diversification methods in the 'Tapestree' package for Julia (*11*), available at https://github.com/ignacioq/Tapestree.jl. The software documentation details the simulation, inference and data structures implemented in the package as well as its summary and plotting capabilities.

#### Clades studied

We assembled empirical phylogenetic trees with robust fossil and, if non-extinct, extant representation. We conducted a thorough search in the literature for clades whose evolutionary histories and time-calibration were inferred probabilistically using living and extinct taxa under the Fossilized Birth-Death ('FBD') model (*1*). We only included phylogenetic trees that made inference at the species-level and that included sampled extinct diversity. Table S1 describes the 27 final groups included in our study, where 26 are monophyletic and 1 paraphyletic. The latter, Dinosauria, was included and analyzed as a clade, while acknowledging that we exclude their alive descendants, Aves. In total, they comprise $5,416$ species, where $1,472$ are extant and $3,944$ extinct across 27 phylogenetic trees. On average the proportion of extinct species is $0.68$, with a $50\%$ interquantile range of $[0.41, 0.97]$ and a range of $[0.04, 1.0]$. Overall, their turnover rates under constant speciation and extinction (*i.e.*, $\mu/\lambda$) were found to be high $> 0.45$, even with a small proportion of fossils (Fig. S5).

These groups span a broad taxonomic spectrum, including plants, arthropods, lung- and ray finned-fish, reptiles, and mammals (Table S1). For each clade, we extracted and randomly sampled $100$ posterior trees from the study's supplementary material or, in most cases, by directly emailing the authors of the study. However, for Dipnoi and Folivora, we were only able to obtain the Maximum Clade Credibility (MCC) tree, and 6 different phylogenetic hypotheses for Crocodylomorpha. We removed the outgroups and established their respective sampling fractions. Given the massive amount of computing nodes required if we were to analyze all these trees ($> 3 \times 10^3$ separate nodes, see below), we subset only 10 posterior trees per clade to carry on subsequent analyses while accounting for phylogenetic uncertainty (*12*).

A peculiarity of observed (empirical) trees that include fossils, whose divergence times were inferred under the FBD model, is that the proportion of 'tip fossils' (*i.e.*, those without sampled descendants; Fig. S1) to 'sampled ancestors' (*i.e.*, those with at least one sampled descendants; Fig. S1) is very high. Under the constant FBD model, at least $> 0.5$ of all fossils are expected to be sampled ancestors (*13*), but we observe much less. This difference results primarily from the fact that empirical FBD trees are mostly built using only one fossil (usually the first or last appearance) per extinct taxa, and fossils that are highly likely to belong to extant species are not included. This exclusion, to avoid computational costs, biases the number and placement of fossils and thereby the fossilization rate estimates, which, in turn, can have major effects on the inferred diversification rates (*6, 13*). Consequently, we include fossil occurrences for extinct and extant species represented in the tree (*i.e.*, as sampled ancestors), but that were excluded when the tree was built.

For each clade, we searched the Paleobiology Database (PaleoDB) for all occurrences of fossil taxa that did not come from the same deposit. We used the 'paleobioDB' package (*14*) for R (*15*) to query PaleoDB and matched all occurrences of the fossil taxa in our tree. We first vetted the resulting database by removing occurrences with a reported interval larger than $5$ My for occurrences younger than $23.04$ Mya, larger than $10$ My for occurrences younger than $66.02$ Mya, and larger than $15$ My for all other occurrences. If multiple occurrences came from the same geographical location and time, we only counted them as one. Finally, we discarded one of the fossil occurrences per species (if there was more than one), which represents the one in the tree, and included the rest as sampled ancestors in the observed tree.

#### Diversity and diversification analyses

We performed inference on each of the posterior trees for each clade under the FBDD model. Each run combined information of the phylogenetic tree concomitant with their clade specific sampling fractions with the external fossil occurrences and was run for $5.5 \times 10^6$ iterations, where the first $5 \times 10^5$ were discarded as burnin, and sampling every $25 \times 10^3$. We used predefined $K$ periods of separate fossilization rates based on the major stratigraphic Epochs (specifically, towards the past in Mya: $2.58, 5.33, 23.03, 33.9, 56, 66.02, 100.5, \ldots,$ a full list is available as part of the 'Tapestree.jl' package). We set the following priors, which worked well under our simulation study: $p(\lambda_0) = p(\mu_0) =$ Log-N$(0.05, Inf)$ (*i.e.*, effectively flat prior), $p(\alpha_\sigma) =$ N$(0.0, 1.0)$, $p(\alpha_\mu) =$ N$(0.0, 1.0)$, $p(\alpha_\sigma) =$ N$(0.0, 1.0)$, $p(\sigma_\lambda) = \Gamma^{-1}(3.0, 0.01)$, $p(\sigma_\mu) = \Gamma^{-1}(3.0, 0.01)$ (the influence of priors on the diffusion coefficients was explored in (*2*)). Every sample saves the set of FBDD parameters along with the complete tree at a particular iteration, thus yielding a posterior distribution of 200 posterior parameter and complete (DA) trees sampled. For the FBDD models applied to empirical data alone, we ran a total of 248 separate analyzes (10 posterior trees for 24 clades, 6 trees for Crocodylomorpha and 1 tree for Dipnoi and Folivora). For each clade we combined the complete trees sampled under the FBDD for each of the 10 posterior trees (or less for the three clades with a lower number of posterior trees), yielding $200 \times 10 = 2000$ complete trees for most clades.

**Rise, decline and fall in diversity** The combined approach of using DA for inference of the FBDD model returns complete trees that incorporate both sampled (*i.e.*, observed in the empirical tree) and unsampled taxa, enabling a robust characterization of past diversity dynamics (*2, 16*). Importantly, this characterization takes into account incomplete species sampling and heterogeneous diversification rates across lineages and through time. For each clade we estimated the diversity through time trajectories across each of the complete trees and then uniformly sampled this curve at 100 regular time steps spanning the maximum duration of the clade (*i.e.*, the maximum tree height; fig. S6). We took the average diversity through time for each clade (Fig. 1A), and simply designated clades on the *rise* as those holding the highest diversity at the present, on *decline* as those holding less extant diversity than in the past, and those that *fall* as those that have no extant representatives.

We then measured the asymmetry between the overall increase and decrease in diversity, that is before and after the global peak in diversity. Since measuring asymmetry in clades on the rise is not possible, we do so only for clades in decline (while acknowledging that the present state of decline could be circumstantial) and that fell. We first use the 'Center of Gravity' metric (CG) (*17*), which estimates when the average diversity was accumulated along standard time, such that $\frac{\sum_i N(t_i)t_i}{\sum_i N(t_i)}$, where time $t_i$ is scaled to be between $[0, 1]$ and $N(t_i)$ is the number of species at time $t_i$. Overall, a CG $< 0.5$ means that average diversity is higher at the beginning ('bottom-heavy'), while $> 0.5$ indicates higher diversity towards the end ('top-heavy').

The CG metric will be biased for diversity trajectories in clades that have not gone extinct (*i.e.*, clades in decline, since not all diversity of the clade is accounted for), and cannot make any future prediction in these cases. For this reason, we also used a hierarchical parametric model to, first, enable flexible predictions of asymmetry for clades in decline, and, second, to properly integrate over all the probable diversity realizations given by the posterior complete trees for each clade. In brief, we specify a two-level Negative Binomial regression model. Specifically, for each complete tree $j$, we specify the following expectation of the number of species at time $t$, $\bar{N}_j(t)$,

$$\bar{N}_j(t) = N_0 \exp\left\{-\frac{r_j}{u_j}\left(1 - e^{u_j t}\right) + \frac{v_j}{g_j}\left(1 - e^{g_j t}\right)\right\} \tag{4}$$

where $N_0$ is the starting number of species (*i.e.*, 1 for stem and 2 for crown trees), and $r_j$, $u_j$, $v_j$ and $g_j$ are the parameters for complete tree $j$. Note that this function is the solution to the diversity expectation of a birth-death model with log-linear changes in birth and death rates (as in (*18*) but in forward time). We employ this function since it strikes a balance between allowing for sufficient complexity in describing asymmetrical wax and wane diversity dynamics and simplifying the many stochastic vicissitudes observed in the diversity curves across complete trees (fig. S6). Moreover, it enables an unbiased estimation of the diversity shape for clades in decline. To estimate the probabilities, we assume that diversity estimates are distributed around the Expectation following a Negative Binomial distribution, which adequately represents the stochastic variance of richness under birth-death models (*19*), such that $N_j(t) \sim \text{NegBinomial}(\bar{N}(t)_j, \gamma)$, where $\gamma$ controls the overdispersion relative to that from a Poisson distribution. We assume this overdispersion, $\gamma$, is shared across all of the complete trees. Finally, we consider each complete tree to be a stochastic estimate of the true underlying diversity trajectory of a clade, such that $r_j \sim \text{LogNormal}(r, s_r),\ u_j \sim \text{LogNormal}(u, s_u),\ v_j \sim \text{LogNormal}(v, s_v),\ g_j \sim \text{LogNormal}(g, s_g)$, where $s_r$, $s_r$, $s_r$, and $s_g$ represent the variance across each of the four parameters in the diversity trajectory.

We then used $r$, $u$, $v$, and $g$ to construct a diversity function for each clade using posterior predictive sampling from the diversity trajectory distributions. We implemented and performed inference of this model in a Bayesian framework using Stan (*20*) through the 'RStan' package (*21*) for R (*15*). We defined ambiguous priors for all parameters by not specifying them (*21*) and ran models for $2 \times 10^3$ iterations, discarding the first $10^3$ as burn-in. For a few clades, very small changes in parameters can dramatically change the shape of the function from Eq. 4, which led to bad behavior during inference. In this case, we alleviated the problem by specifiying sensible initial parameter values. This function (Eq. 4), however, is fixed only on the initial diversity $N_0$, but not on the final diversity, as in (*22*), simply because a regression of diversity through time based on these probabilities would be computationally prohibitive. We note that this biases the measure of asymmetry towards 'bottom-heavy' patterns, since for some regressions of diversity that end up going extinct quickly, the regression is not able to predict such steep diversity decrease. Following this procedure, we obtain diversity trajectories that "extend into the future", therefore avoiding the limitations of the CG metric for clades in decline.

**Testing for diversity-dependent diversification** We test whether speciation or extinction rates are associated with clade richness, as predicted under diversity-dependence diversification hypotheses, in particular, the idea that the level of standing richness of a clade affects its own diversification dynamics when reaching some carrying capacity (*23, 24*). We examined whether average speciation and extinction rates are each associated with diversity through time using a three-level hierarchical auto-regressive model of order 1 (higher orders were not found significant). This model takes into account the correlation in time of rates, that is, rates at a given time is strongly predicted by those just before, while examining the effect of diversity. Thus,

$$\bar{r}_{c,j}(t) \sim \text{LogNormal}(\beta_{0,c,j} + \beta_{1,j}\bar{r}_{c,j}(t-1) + \beta_{2,c,j}d_{c,j}(t-1), \gamma_{c,j}),$$

where $\bar{r}_{c,j}(t)$ represents the average rate across lineages, either speciation or extinction, $d_{c,j}(t)$ the richness estimated directly from the FBDD model at time $t$, and $\gamma_{c,j}$ the rate variance, for complete tree $j$ in taxon $c$. Note that we account for the fact that the rates, the response, are auto-correlated along time by including the $\beta_{1,j}\bar{r}_{c,j}(t-1)$ term. Then we have

$$\beta_{0,c,j} \sim \text{N}(\beta_{0,c}, \gamma_{0,c});\ \beta_{1,c,j} \sim \text{N}(\beta_{1,c}, \gamma_{1,c});\ \beta_{2,c,j} \sim \text{N}(\beta_{2,c}, \gamma_{2,c}),$$

such that $\beta_{2,c}$ represents the effect of diversity on each speciation and extinction rates across all samples for a given clade $c$. Because the amount of observed species information across clades varied dramatically, we weighted the $\beta_{2,c}$ associated to each clade by the number of observed species in that clade (extant and extinct tips and sampled ancestors in the observed phylogeny). Thus, the overall effect of diversity on either rate across all clades, $\beta_2$, is given by specifying

$$p(\beta_2|\cdot) \propto \prod_c (\beta_{2,c} \sim \text{N}(\beta_2, \gamma))^{w_c}$$

where $\gamma$ represents the variance across clades, and $w_c$ the weight for the number of observed species in the empirical tree. The global parameter $\beta_2$ represents the percentage change in rate with the addition of one species. Because if we were to use all of the posterior complete trees for each clade, the number of data points would be $> 0.5 \times 10^6$, and $> 3 \times 10^4$ parameters, we sampled $100$ trees out of the 200, thus reducing the input data (and computational cost) by more than half. We ran this model for speciation and extinction rates in 'RStan' (*21*) for R (*15*) using $5$ independent chains, each with $500$ warm-up and $500$ logging iterations and visually checked for convergence.

We note that, as with previous tests on diversity-dependence on individual clades (*e.g.*, (*23, 25, 26*)), our analyses assume that all species are able to interact with one another and that these interactions are limited to only those species within the clade.

**Asymmetry in diversity trajectories** We then measured asymmetry by integrating the diversity trajectory (including the part of the curve that extends to the future) left and right from the inferred point of maximum diversity from and up to $N(t) = N_0$, that is, from the initial diversity to the maximum (we note this integrated diversity $S_l$) and from the maximum to the predicted time when reaching $N_0$ again (we note this integrated diversity $S_r$). For most clades $N_0 = 2$ (for those phylogenies that start with a stem fossil $N_0 = 1$). To get a comparable metric to CG, and interpretable between $[0, 1]$, we measure asymmetry as $S_l/(S_l + S_r)$, that is, the proportion of integrated diversity before the peak of diversity over all the integrated diversity. In general, diversity trajectories that are bottom-heavy have values of asymmetry and CG $< 0.5$, those around $0.5$ are largely symmetrical, while values $> 0.5$ are top-heavy (Fig. 1B & fig. S6).

To ascertain the degree to which asymmetry values emerge simply from stochasticity in the diversification process we performed simulations under a constant birth-death process (*27*). For a radiation to go extinct, the number of extinction events must equal the number of speciation events plus one, which materialize on relatively equal speciation and extinction rates under a constant birth-death process. Thus, to obtain clades with diversity trajectories of decline and fall, we simulated trees with equal speciation and extinction rates of $1$ spp time$^{-1}$ and ran the process during $10$ time units. We first considered only simulations with at least $100$ extinct tips to obtain diversity trajectories with a reasonable number of species with respect to our taxonomic groups (*i.e.*, trajectories with about 20-40 species at their peak). We retained $200$ simulations of clades with decline diversity trajectories (*i.e.*, where diversity was higher before the termination) and $200$ with fall diversity trajectories (*i.e.*, where all species went extinct before termination). We then followed the same procedure as above to measure CG and asymmetry on each of the $400$ diversity curves. We removed asymmetry values for those few simulated clades in decline not detected by the parametric approach. Some samples of these simulations are shown in fig S7.

**Relative importance of speciation and extinction** Ultimately, the observed diversity patterns are the result of the underlying diversification process, where diversity increases with positive diversification and decreases when diversification is negative. However, distinct combinations of speciation and extinction dynamics can be responsible for changes in diversification, the latter being defined as speciation minus extinction. For instance, an increase in diversification can result from an increase in speciation, a decrease in extinction, or both. The FBDD model provides the means to evaluate their relative role.

We estimated average posterior speciation, $\bar{\lambda(t)}$, extinction, $\bar{\mu(t)}$, and diversification, $\bar{\delta(t)}$ through time for evolutionary radiation across all lineages and all complete tree samples (figs. S9-S35). We then estimated their respective change, that is, the first derivative with respect to time, $\bar{\lambda(t)}'$, $\bar{\mu(t)}'$ and $\bar{\delta(t)}'$, such that $x(t)' = (x(t + \delta t) - x(t))/\delta t$. With these quantities, we can then evaluate the relative roles of speciation and extinction in driving changes in diversification (Fig. 2A-C). Specifically, we estimate an 'Importance' metric for speciation, $I_s$ and extinction, $I_e$, such that $I_s = \frac{\int_T \mathrm{sgn}(\bar{\delta(t)})\bar{\lambda(t)}' dt}{\int_T \mathrm{sgn}(\bar{\delta(t)})\bar{\delta(t)}' dt}$ and $I_e = \frac{\int_T -\mathrm{sgn}(\bar{\delta(t)})\bar{\mu(t)}' dt}{\int_T \mathrm{sgn}(\bar{\delta(t)})\bar{\delta(t)}' dt}$, where the $\mathrm{sgn}(x)$ is the signum function, returning the sign of $x$. Notice that $I_s + I_e = 1$. Intuitively, the Importance measures the proportion of change in diversification explained by either speciation or extinction integrated over period $T$. For example, if diversification increased $0.5$ over a time unit of $1$, following an increase of speciation of $0.3$ and a decrease of extinction of $0.2$, then $I_s = 0.6$ and $I_e = 0.4$. Sometimes, the Importance of a rate can be less than $0$ if the change in diversification is in the opposite direction of the rate change (*e.g.*, if diversification decreases in spite of an increase in speciation, then $I_s < 0$) or more than $1$ if the change in diversification is in the same direction of the rate change, yet the the other rate acts in the opposite direction (*e.g.*, if diversification increases following an increase in speciation in spite of an increase in extinction, then $I_s > 1$). Since both an Importance of $1$ or larger than $1$ reflect that all of the change in diversification is being driven by a particular rate (and of $0$ or below $0$ reflect that the other rate is fully responsible), and for simplicity in further computations, we cap the Importance to be between $[0, 1]$. Notice that $I_s + I_e = 1$.

To understand the role of speciation and extinction at different diversity stages, we numerically calculate $I_s$ and $I_e$ at intervals following the following criteria (Fig. 2): *i)* accelerating diversity gain (*i.e.*, $\bar{\delta(t)} > 0$ & $\bar{\delta(t)}' > 0$); *ii)* decelerating diversity gain (*i.e.*, $\bar{\delta(t)} > 0$ & $\bar{\delta(t)}' < 0$); *iii)* deceleratin diversity loss (*i.e.*, $\bar{\delta(t)} < 0$ & $\bar{\delta(t)}' > 0$); and *iv)* accelerating diversity loss (*i.e.*, $\bar{\delta(t)} < 0$ & $\bar{\delta(t)}' < 0$). We perform these computations for each evolutionary radiation.

Finally, to test whether the role of speciation and extinction was more or less prevalent at each of these four stages of diversity across the taxonomic groups, we estimated the Importance of speciation, $I_s$ (note that $I_e$ is just $1 - I_s$), and used it as the probability from a Bernoulli distribution for each of the diversity stages (*i-iv*). However, we did not consider that all evolutionary radiations contributed equally to $I_s$ at each diversity stage; rather, we weighted $I_s$ by the number of observed species in the empirical trees, as a measure of information about evolutionary dynamics, multiplied by the length of time (in My) spent in each given diversity stage. That is, for a given diversity stage, more weight is given to groups with more species information and that spent more time in such stage, proportionally. The posterior distribution for these parameters can be analytically determined by assuming the following prior $p(p_h) = \text{Beta}(\varrho, \varsigma)$. Let $x_h = \frac{\sum_c 1}{\sum_c w_c w_{t,c,h}} \sum_j I_{s,c,h} w_c w_{t,c,h}$, where, for each taxonomic group $c$, $w_c$ is the standardized weight based on the number of species and $w_{t,c,h}$ is the standardized weight of the amount of time spent in diversity stage $h$. Letting $n_h$ the total number of clades at stage $h$, then the posterior distribution for $p_h$ is

$$p(p_h | x_h, n_h) = \text{Beta}(\varrho + x_h, \varsigma + n_h - x_h).$$

Note that if $I_{s,j,}$ is 0 or 1 and all weights are equal, $x_h$ would equal the sum of time in which $I_s = 1$, as in a unweighted Binomial distribution with a Beta prior. We specified $\varrho = 1$ and $\varsigma = 1$ as a prior (*i.e.*, the Uniform standard distribution, $\text{U}(0, 1)$) and characterized this distribution for each diversity stage (Fig. 2E).

**Subclade analyses** To create more comparable evolutionary units of analyses across taxa, which greatly differ in their age, time and diversity trajectories, we generated, for each complete tree, monophyletic and exclusive 'subclades' of size between 20 to 40 species. This was done by simply starting at the root of each tree, and then testing whether the left or right daughter consists of a subclade with more than 20 but less than 40 tips; if true, the subclade was saved, if not, we continued in recursive fashion in the subsequent descendant nodes. For clarity, we talk of 'taxa' when we refer to the 27 groups already analyzed as units, and of 'subclades' when we refer to these restricted to having between $20 - 40$ tips. We delineated such subclades on the posterior sample of 200 complete (DA) trees for each of the 27 taxa. Given the size differences between taxa, there were large differences in the total number of subclades per taxa (*e.g.*, $> 10^4$ in Dinosauria to just 4 in Dipnoi), and thus we randomly selected a maximum of 200 subclades per taxa to continue our analyzes. While we acknowledge that these subclades can share fairly complex patterns of non-independence (*i.e.*, belonging to the same complete tree sample or sharing some observed lineages across complete trees), our posterior analyses either account for some of them or are not particularly impacted, so we treat them simply as a posterior distribution of diversity estimates.

As at the taxon level, we first analyze the asymmetry in the diversity trajectories experienced by these subclades. We simplify Eq. 4 to analyze each subclade separately, such that the expected diversity follows:

$$\bar{N}(t) = N_0 \exp\left\{-\frac{r}{u}\left(1.0 - e^{ut}\right) + \frac{v}{g}\left(1.0 - e^{gt}\right)\right\}, \qquad (5)$$

with Negative Binomial deviations from the mean and performed Bayesian optimization using 'RStan' (*21*) in R (*15*), as done at the clade level. Given difficulties in fitting the curve from Eq. 5 across very different diversity trajectories (see Figs. S36-S37), for each subclade we ran 10 separate chains of 900 iterations (discarding the first 400 as burnin), each with different initial parameter settings and selected the one with the highest posterior probability. These analyses consisted initially of at least $50,000$ separate runs. We then plotted the diversity regression to check for inadequate fits where the regression parameters clearly did not follow the diversity trajectory and thus the optimization failed, and discarded pathological fits while performing new runs on other subclades if available. A small set of these subclade diversities and regression results is shown in Figs. S36-S37. We then assigned the diversity trajectories of each subclade as on the rise, on decline or fall (based on the diversity curves), and for the last two, estimated the center of gravity and the parametric asymmetry metric as done for the clade level (Fig. 3A).

We repeated 400 null simulations, 200 for decline and 200 for fall diversity trajectories, to assay the amount of stochasticity expected under no deterministic causes, as with the taxon-level. We used the same parameters but retained only those radiations that result in 20 to 40 total species (instead of 100 extinct species), since a lower number of stochastic speciation and extinction events should result in a wider range of asymmetry values.

Finally, to analyze the relative role of speciation and extinction on these diversity patterns, we estimated average speciation and extinction for each subclade and calculated their Importance, $I_s$ and $I_e$, for diversity stages **i-iv** (Fig. 2D), as done for taxa. We then estimated combined proportions of speciation Importance across all subclades in a hierarchical model that takes into account that subclades are nested across different clades, such that, for each subclade $j$, in clade $c$, at diversity stage $h$,

$$I_s^{h,c,j} \sim \text{Beta}(\theta_{h,c}, \beta_{h,c})$$

and

$$\theta_{h,c} \sim \Gamma(\varrho_{\theta,h}, \varsigma_{\theta,h}), \tag{6}$$

$$\beta_{h,c} \sim \Gamma(\varrho_{\beta,h}, \varsigma_{\beta,h}), \tag{7}$$

where $\text{Beta}_h(\frac{\varrho_{\theta,h}}{\varsigma_{\theta,h}}, \frac{\varrho_{\beta,h}}{\varsigma_{\beta,h}})$ describe the overall distribution for the Importance of speciation for diversity stage $h$ (Fig. 3B). We implemented and made inference under this model using 'RStan' (*21*) in R (*15*) for each diversity stage, with each run consisting of 2 chains each with 1000 warmup iterations followed by 5000 sampling iterations, sampling the posterior predictive distribution of $\text{Beta}_h$.

**Overall trends in speciation and extinction** To estimate cross-taxa trends in lineage-specific speciation and extinction rates, $\alpha_\lambda$ and $\alpha_\mu$, respectively, we used a simple Bayesian hierarchical model that incorporates the uncertainty around the estimates for every posterior distribution. Because drift is (approximately) normally distributed, for each tree $j$ of each clade $c$ we estimated the mean, $\bar{m_{c,j}}$, and variance, $\bar{s_{c,j}}$, of $\alpha_\lambda$ and, subsequently, for $\alpha_\mu$, and used them as normal priors $\alpha_{c,j} \sim N(\bar{m_{c,j}}, \bar{s_{c,j}})$. We then specified the following hierarchical model for $\alpha_\lambda$ and, separately, for $\alpha_\mu$

$$\alpha_{c,j} \sim N(\alpha_c, \sigma_c), \tag{8}$$

$$\alpha_c \sim N(\alpha_g, \sigma_g). \tag{9}$$

$\alpha_g$ would then represent the overall cross-taxa trend for lineage-specific speciation or extinction, while properly weighting for the amount of information in each tree, its uncertainty, and the tree of each clade. We implemented and made inference under this model using 'RStan' (*21*) in R (*15*) using 2 chains each with 1000 warmup iterations followed by 1000 sampling iterations.

### Supplementary Text

#### Constant rate Fossilized Birth-Death model

In the constant rate fossilized birth-death model (CFBD), at any time, all lineages share the same speciation, extinction, and fossilization rate $\lambda$, $\mu$, and $\psi$, respectively (*i.e.*, $\lambda_l(t) = \lambda$, $\mu_l(t) = \mu$ and $\psi_l(t) = \psi$). While there is an analytical solution of this likelihood, here we develop the DA approach, which yields the same exact posterior distribution of the parameters as its closed-form counterpart, but also yields posterior complete trees.

**Likelihood** The likelihood for a complete unordered crown tree under a CFBD process is simply

$$\ell(\Psi|\lambda, \mu, \psi) = \lambda^{s-1}\mu^{z}\psi^{f}\mathrm{e}^{-(\lambda+\mu+\psi)L},$$

where $s$, $z$ and $f$ are the number of speciation, extinction and fossilization events, respectively, and $L$ is the tree length (sum of all branches). For a stem tree, we do consider all $s$ speciation events.

**Bayesian data augmentation algorithm** To integrate over $\Psi$ during MCMC, we sample $\Psi_a$ directly from its conditional distribution given $\Psi_o$ and the model parameters (*i.e.*, Gibbs sampling) using forward simulation. We follow (*2, 10*) to impute probable hidden configurations of unobserved lineages because they went extinct or they were unsampled at the present. This procedure generates *complete trees* (*i.e.*, data augmented trees) which are concordant with the empirical (reconstructed) tree under the diversification model. (*2*) described the proposals for any general diversification model when the empirical tree consisted on extant-only lineages. Here we briefly describe the forward simulation procedure, highlighting the differences when making inference under a fossilized birth-death model.

We iterate over each branch in the empirical tree (a branch constitutes the length of time between speciation or fossilization events) and simulate fossilized birth-death processes forward in time throughout their branch lengths ($t_b$ for branch $b$), given the current iteration model parameters. For any branch, each initial simulation is a valid proposal if it survives throughout the branch length $t_b$ and if no other fossilization events occur. For internal branches, if there is more than one surviving lineage by $t_b$, we uniformly choose one at random and continue the simulation in the other ones. For branches followed by a speciation event the selected lineage represents the one that underwent speciation and is left as is, for branches followed by a fossilization event, the simulation is also pursued on the selected lineage. These simulations remain valid proposals if all unobserved lineages do not leave any fossilization events and either go extinct or are sampled with probability $1 - \rho_b$, where $\rho_b$ is the branch specific sampling fraction (*2, 10*). By default we assume that tip fossils did not leave unsampled descendants at the present.

Let $\psi_b$ denote the forward simulated tree in branch $b$, $n_b(t_b)$ denote the number of lineages alive at time $t_b$ (*i.e.*, at the end of the branch) and $n_b(0)$ denote the number of lineages alive at the present for $\psi_b$, then the acceptance ratio in the Metropolis-Hastings (MH) step for the proposed forward simulation $\psi'_b$ if $b$ is a branch of an extant lineage is

$$a = \min\left\{1, \frac{\ell(\Psi'|\lambda,\mu,\psi)\,\ell(\psi_b|\lambda,\mu,\psi)}{\ell(\Psi|\lambda,\mu,\psi)\,\ell(\psi'_b|\lambda,\mu,\psi)} \times \frac{n'_b(0)\rho_b(1-\rho_b)^{n'_b(0)-1}}{n_b(0)\rho_b(1-\rho_b)^{n_b(0)-1}}\right\},$$

and for the other branches $b$ is

$$a = \min\left\{1, \frac{\ell(\Psi'|\lambda,\mu,\psi)\,\ell(\psi_b|\lambda,\mu,\psi)}{\ell(\Psi|\lambda,\mu,\psi)\,\ell(\psi'_b|\lambda,\mu,\psi)} \times \frac{1/n_b(t_b)}{1/n'_b(t_b)} \times \frac{(1-\rho_b)^{n'_b(0)}}{(1-\rho_b)^{n_b(0)}}\right\},$$

where the $1/n_b(t_b)$ factor comes from the proposal density of randomly and uniformly choosing one of the extant lineages at time $t_b$ as the one sampled in the reconstructed tree. These acceptance ratios simplify then for all branches to

$$a = \min\left\{1, \frac{n'_b(t_b)}{n_b(t_b)}(1-\rho_b)^{n'_b(0)-n_b(0)}\right\}.$$

The MH acceptance probability for the sampling fraction is common to all further models, so, for simplicity, we do not show it in the rest of the manuscript.

The model parameters of the process, $\lambda$, $\mu$ and $\psi$, are updated using Gibbs sampling using the Gamma conjugate prior, $\Gamma(\kappa, \varsigma)$, for a Poisson process, from which we sample directly

$$p(\lambda|\cdot) = \Gamma(\kappa + s, \varsigma + L),$$

and similarly with $\mu$ and $\psi$ but using the number of extinction $z$ and fossilization $f$ events, respectively, in the complete tree of the current iteration. If there is conditioning on survival (see Materials and Methods), an additional MH step is needed using the pseudo-marginal approach described in (*2*).

**Validation for the CFBD** We validate our data augmentation implementation of the constant rate fossilized birth-death (CFBD) by comparing the rate posteriors with the analytical solution for the likelihood from (*28*) implemented in a Bayesian framework in the 'RevBayes' software (*29*) using Gamma priors $\Gamma(1.0, 1.0)$ across all three parameters (Fig. S2).

#### The episodic Fossilized Birth-Death model (eFBD)

We expand the CFBD to enable variation of fossilization rates at *a priori* defined periods, which can reflect differences across stratigraphic ranges (*3, 4*), which we call the episodic Fossilized Birth-Death model (eCFBD). While the eCFBD can easily accommodate different rates of speciation and extinction during different periods, we simply focus here on relaxing the constancy of fossilization rates as a build-up validation step towards the FBDD process (Materials and Methods). Thus, we allow the fossilization rate to vary temporarily following a piecewise constant function across $K$ predefined periods and estimate different fossilization rates delimited by $|k| - 1$ times, $t^{\psi}$, such that $t_k^{\psi} \in [T_{\Psi}, T_0]$ where rates become $\psi_l(t) = \psi_k$, for $t_k^{\psi} < t < t_{k+1}^{\psi}$. The likelihood for a complete tree under this model then becomes

$$\ell(\Psi|\lambda, \mu, \psi) = \lambda^{s-1}\mu^{z}\mathrm{e}^{-(\lambda+\mu)L} \prod_{k\in K} f_k \psi_k \mathrm{e}^{-\psi_k L_k},$$

where $f_k$ and $L_k$ are the number of fossilization events and tree length, respectively, during period $k$. The data augmented algorithm is then refined to accommodate this new source of variation by taking into account the change in rates when traversing periods.

**Simulation study for the eFBD** We explored the statistical behavior and validity of our inference algorithm under the eFBD using simulations. We simulated trees for a duration of 6 time units with three periods of equal duration ($K = 3$), from $[6, 4]$, $[4, 2]$ and $[2, 0]$ under 4 simulation scenarios with different temporal variation of fossilization rates: *i)* rates remain constant (*i.e.*, $\psi_1 = \psi_2 = \psi_3 = 0.2$), *ii)* rates increase (*i.e.*, $\psi_1 = 0.05; \psi_2 = 0.2; \psi_3 = 0.5$), *iii)* rates decrease and then increase (*i.e.*, $\psi_1 = 0.5; \psi_2 = 0.1; \psi_3 = 0.5$), and *iv)* rates decrease (*i.e.*, $\psi_1 = 0.5; \psi_2 = 0.2; \psi_3 = 0.05$). For each scenario we simulated 50 trees of size between 200 and 500 tips (either alive or fossil), and set $\lambda = 1.0$ and $\mu = 0.5$. We then made inference with Gamma priors $\Gamma(1.0, 1.0)$ across all parameters using our DA algorithm by running $5.2 \times 10^5$ iterations, discarding the first $2 \times 10^4$ as burnin and sampling every $5 \times 10^2$ iterations. The results are shown in Fig. S3. Overall, the inference under the eFBD model shows good statistical behavior, with good accuracy (Mean Absolute Error, MAE, being below 0.15 across parameters) and over 90% coverage for $\lambda$ and $\psi$, and over 86% coverage for $\mu$ across all simulations. As expected the variance for $\psi_k$ increases as one moves further back in time (*i.e.*, variance($\psi_1$) > variance($\psi_2$) > variance($\psi_3$); Fig. S3).

#### Metropolis-Hasting (MH) ratios for the FBDD model

Here we describe those additional MH ratios for the FBDD on top of those already described for the BDD model in (*2*). Let $\Psi, \Lambda, \mathrm{M}$ represent the topology, the latent speciation GBM and the latent extinction GBM, respectively, then the simplified acceptance ratio for a valid forward simulation proposal for a branch $pr$ that ends in a sampled ancestor with subsequent branch $d$ is

$$a = \min\left\{1, \frac{p(\lambda(t_{pr})|\lambda(t_d), \alpha_\lambda \sigma_\lambda)\, p(\mu(t_{pr})|\mu(t_d), \alpha_\mu \sigma_\mu)}{p(\lambda(t_{pr})'|\lambda(t_d), \alpha_\lambda \sigma_\lambda)\, p(\mu(t_{pr})'|\mu(t_d), \alpha_\mu, \sigma_\mu)} \times \frac{\ell(\Lambda', \mathrm{M}'|\Psi, \alpha_\lambda, \alpha_\mu, \sigma_\lambda, \sigma\mu, \psi, \mathcal{M}_{\text{FBDD}})}{\ell(\Lambda, \mathrm{M}|\Psi, \alpha_\lambda, \alpha_\mu, \sigma_\lambda, \sigma\mu, \psi, \mathcal{M}_{\text{FBDD}})}\right\},$$

where

$$p(\lambda(t_{pr})|\lambda(t_d), \alpha_\lambda, \sigma_\lambda) = \lambda(t_{pr}) \sim \mathrm{N}\left(\lambda(t_d) - \alpha_\lambda t_d, t_d \sigma_\lambda^2\right),$$

and similarly for $p(\mu(t_{pr})|\mu(t_d), \alpha_\mu, \sigma_\mu)$. For a tip fossil, the forward simulation is accepted or rejected based on it being a valid proposal as described in the Materials and Methods.

#### Simulation study for the FBDD

We conducted a simulation study to explore the behavior of the FBDD under our DA inference algorithm. Following (*2*), we take advantage of the following approximation to simulate under the FBDD: let $V$ be a random variable for the time of an event and let $r_i$ and $r_{i+1}$, be the event rate at time $t_i$ and $t_{i+1}$, respectively, where $t_{i+1} - t_i = \delta \geq 0$, then we have

$$p(t_i \leq V < t_{i+1} \mid t_i \leq V) \approx \bar{r}_{i,i+1}\delta.$$

To simultaneously explore the role of increasing fossil sampling across parameter space, we considered 5 scenarios with incremental fossilization rate. Specifically, we set $\psi$ to be $\{0.05, 0.1, 0.5, 1.0, 2.0\}$ and, for each fossil rate scenario, simulated 100 FBDD trees of 150 extant species by sampling from a range of parameter space (Fig. S3), yielding a total of 500 simulations. For each simulation, we set $\lambda_0 = 0.3$ and $\mu_0 = 0.25$ and sample $\alpha$ from the uniform distribution $\text{U}(-0.5, 0.5)$ and $\sigma_\lambda$ and $\sigma_\mu$ from the uniform distribution $\text{U}(0.1, 0.6)$. We start each simulation with two lineages and make sure that at least one survived to the present. We then sample a tree with a determined number of extant species following (*30*) to avoid biases.

For each simulation we then conducted inference specifying weakly informative priors for most parameters. Specifically, we used log-normal prior with infinite variance for $\lambda_0$ and $\mu_0$ (*i.e.*, effectively flat), an Inverse Gamma $\Gamma^{-1}(3, 0.1)$ for $\sigma_\lambda$ and $\sigma_\mu$, and a Gaussian $\text{N}(0, 1)$ for $\alpha_\lambda$ and $\alpha_\mu$. We note that an informative prior on $\sigma_\mu$ helps avoids pathological behavior when, for low values of $\psi$ and by chance, the resulting empirical tree results with few tip fossils. Note that the number of tip fossils are the very important for extinction rates, since each tip fossils forces at least one extinction event. In empirical trees, the number of tips fossils are usually over-represented because researchers usually include only one fossil occurrence per extinct species, while sampled ancestors are under-represented. While many tip fossils (and fossils in general) yield better extinction estimates, an under-representation of sampled ancestors in empirical datasets can be problematic and is explained in the following section and in the

Materials and Methods. We run each simulation for $3.5 \times 10^6$ iterations, discarding the first $5 \times 10^5$ as burnin and sampling every $3 \times 10^3$ iterations.

**Statistical coverage and accuracy** Overall, we find good statistical behavior from our inference algorithm of the FBDD model (Fig. S4). First, as expected from our simulation from eCFB, we can reliably retrieve the fossilization rate $\psi$. Second, and as expected from results from extant-only trees (*2*), $\alpha_\lambda$, $\alpha_\mu$, $\sigma_\lambda$ and $\Lambda$ (the whole speciation diffusion) show good coverage and accuracy across all fossilization scenarios (Fig. S4). Nonetheless, we find that M, the extinction diffusion, performs much better as the fossilization rate increases. Specifically, at $\psi = 0.05$ and $\psi = 0.1$ there is an underestimation of extinction rates, and while median coverage is around $95\%$, many simulations have a much lower coverage. However, with fossilization rates $\psi > 0.5$, the statistical behavior of M increases sharply, with almost no bias, small relative error and good coverage (Fig. S4). Finally, while the accuracy of $\sigma_\mu$ is very low, coverage remains moderate (over $50\%$), implying that while extinction is retrievable, its rate of diffusion is not, and the posterior distribution integrates over much of its parameter space (Fig. S4).

### References


1. T. A. Heath, J. P. Huelsenbeck, T. Stadler, *Proceedings of the National Academy of Sciences* **111**, E2957 (2014).

2. I. Quintero, N. Lartillot, H. Morlon, *Science* **384**, 1007 (2024).

3. S. M. Kidwell, S. M. Holland, *Annual Review of Ecology and Systematics* **33**, 561 (2002).

4. D. Silvestro, N. Salamin, J. Schnitzler, *Methods in Ecology and Evolution* **5**, 1126 (2014).

5. W. P. Maddison, P. E. Midford, S. P. Otto, *Systematic biology* **56**, 701 (2007).

6. J. S. Mitchell, R. S. Etienne, D. L. Rabosky, *Systematic Biology* **68**, 1 (2019).

7. T. B. Quental, C. R. Marshall, *Science* **341**, 290 (2013).

8. N. Metropolis, A. W. Rosenbluth, M. N. Rosenbluth, A. H. Teller, E. Teller, *The journal of chemical physics* **21**, 1087 (1953).

9. W. K. Hastings (1970).

10. O. Maliet, H. Morlon, *Systematic biology* **71**, 353 (2022).

11. J. Bezanson, A. Edelman, S. Karpinski, V. B. Shah, *SIAM Review* **59**, 65 (2017).

12. J. P. Huelsenbeck, B. Rannala, J. P. Masly, *Science* **288**, 2349 (2000).

13. W. Pett, T. A. Heath, *Phylogenetics in the Genomic Era*, C. Scornavacca, F. Delsuc, N. Galtier, eds. (No commercial publisher | Authors open access book, 2020), pp. 5.1:1–5.1:18.

14. S. Varela, J. González Hernández, L. Fabris Sgarbi, *paleobioDB: Download and Process Data from the Paleobiology Database* (2020). R package version 0.7.0.

15. R Core Team, *R: A Language and Environment for Statistical Computing*, R Foundation for Statistical Computing, Vienna, Austria (2023).

16. R. C. Warnock, T. A. Heath, T. Stadler, *Paleobiology* **46**, 137 (2020).

17. S. J. Gould, N. L. Gilinsky, R. Z. German, *Science* **236**, 1437 (1987).

18. H. Morlon, M. D. Potts, J. B. Plotkin, *PLoS biology* **8**, e1000493 (2010).

19. I. Quintero, M. J. Landis, W. Jetz, H. Morlon, *Proceedings of the National Academy of Sciences* **120**, e2220672120 (2023).

20. B. Carpenter, *et al.*, *Journal of statistical software* **76** (2017).

21. Stan Development Team, RStan: the R interface to Stan (2023). R package version 2.26.22.

22. O. Billaud, D. Moen, T. L. Parsons, H. Morlon, *Systematic Biology* **69**, 363 (2020).

23. D. L. Rabosky, *Annual Review of Ecology, Evolution, and Systematics* **44**, 481 (2013).

24. L. J. Harmon, S. Harrison, *The American Naturalist* **185**, 584 (2015).

25. D. L. Rabosky, *Ecology letters* **12**, 735 (2009).

26. R. S. Etienne, *et al.*, *Proceedings of the Royal Society B: Biological Sciences* **279**, 1300 (2012).

27. S. J. Gould, D. M. Raup, J. J. Sepkoski, T. J. Schopf, D. S. Simberloff, *Paleobiology* **3**, 23 (1977).

28. T. Stadler, *Journal of theoretical biology* **267**, 396 (2010).

29. S. Höhna, *et al.*, *Systematic biology* **65**, 726 (2016).

30. T. Stadler, *Systematic biology* **60**, 676 (2011).

31. A. Ruebenstahl, N. Mongiardino Koch, J. C. Lamsdell, D. E. Briggs, *Proceedings of the Royal Society B* **291**, 20241184 (2024).

32. C. D. Brownstein, R. C. Harrington, T. J. Near, *Journal of Biogeography* **50**, 1191 (2023).

33. H. R. Grunert, N. Brocklehurst, J. Fröbisch, *Scientific Reports* **9**, 5063 (2019).

34. T. R. Simões, M. W. Caldwell, S. E. Pierce, *BMC biology* **18**, 1 (2020).

35. B. M. Farina, P. L. Godoy, R. B. Benson, M. C. Langer, G. S. Ferreira, *Ecology and Evolution* **13**, e10201 (2023).

36. R. B. Benson, R. A. Frigot, A. Goswami, B. Andres, R. J. Butler, *Nature communications* **5**, 3567 (2014).

37. G. Lloyd, D. Bapst, M. Friedman, K. Davis, *Biology letters* **12**, 20160609 (2016).

38. E. W. Wilberg, A. H. Turner, C. A. Brochu, *Scientific reports* **9**, 514 (2019).

39. Q. Zhang, R. H. Ree, N. Salamin, Y. Xing, D. Silvestro, *Systematic Biology* **71**, 242 (2022).

40. E. M. Troyer, *et al.*, *Proceedings of the National Academy of Sciences* **119**, e2122486119 (2022).

41. C. D. Bacon, *et al.*, *Biology Letters* **18**, 20220214 (2022).

42. A. C. Siqueira, D. R. Bellwood, P. F. Cowman, *Journal of Biogeography* **46**, 1611 (2019).

43. A. Gavryushkina, *et al.*, *Systematic biology* **66**, 57 (2017).

44. J. L. Cantalapiedra, *et al.*, *Nature Ecology & Evolution* **5**, 1266 (2021).

45. O. Sanisidro, M. C. Mihlbachler, J. L. Cantalapiedra, *Science* **380**, 616 (2023).

46. G. T. Lloyd, G. J. Slater, *Systematic Biology* **70**, 922 (2021).

47. S. Hotaling, M. L. Borowiec, L. S. Lins, T. Desvignes, J. L. Kelley, *Molecular phylogenetics and evolution* **162**, 107211 (2021).

48. J. L. Cantalapiedra, *et al.*, *Proceedings of the Royal Society B* **286**, 20182896 (2019).

49. M. Cascini, K. J. Mitchell, A. Cooper, M. J. Phillips, *Systematic Biology* **68**, 520 (2019).

50. L. Varela, P. S. Tambusso, H. G. McDonald, R. A. Fariña, *Systematic Biology* **68**, 204 (2019).

51. T. J. Near, D. Kim, *Molecular Phylogenetics and Evolution* **161**, 107156 (2021).

52. D. Silvestro, *et al.*, *Systematic Biology* **68**, 78 (2018).

53. T. Park, *et al.*, *Evolution* p. qpae061 (2024).

54. H. A. Ogilvie, *et al.*, *Systematic Biology* **71**, 208 (2022).

55. J. L. Cantalapiedra, J. L. Prado, M. Hernández Fernández, M. T. Alberdi, *Science* **355**, 627 (2017).

56. H. P. Püschel, O. C. Bertrand, J. E. O'reilly, R. Bobe, T. A. Püschel, *Nature ecology & evolution* **5**, 808 (2021).

### Supplementary Tables

**Table S1: Taxonomic groups used in this study with their respective number of extinct and extant species represented, their present-day sampling fraction ($\rho$) and the reference from where the phylogenetic trees were obtained.**

| taxon | n extinct | n extant | $\rho$ | tree source |
|---|---|---|---|---|
| Eurypterida | 153 | 0 | 1 | (*31*) |
| Dipnoi | 16 | 5 | 0.87 | (*32*) |
| Therocephalia | 51 | 0 | 1 | (*33*) |
| Sphenodontidae | 34 | 1 | 1 | (*34*) |
| Testudinata | 448 | 327 | 0.49 | (*35*) |
| Pterosauria | 109 | 0 | 1 | (*36*) |
| Dinosauria (non-avian) | 624 | 0 | 1 | (*37*) |
| Crocodylomorpha | 127 | 15 | 0.55 | (*38*) |
| Juglandaceae | 130 | 47 | 0.77 | (*39*) |
| Tetraodontiformes | 52 | 187 | 0.53 | (*40*) |
| Mauritiinae | 9 | 9 | 0.9 | (*41*) |
| Acanthuridae | 25 | 72 | 0.9 | (*42*) |
| Siganidae | 10 | 24 | 0.8 | (*42*) |
| Spheniscidae | 36 | 19 | 0.95 | (*43*) |
| Proboscidea | 182 | 3 | 1 | (*44*) |
| Brontotheriidae | 57 | 0 | 1 | (*45*) |
| Cetacea | 405 | 89 | 0.97 | (*46*) |
| Zoarcoidei | 9 | 222 | 0.51 | (*47*) |
| Ruminantia | 1064 | 213 | 1 | (*48*) |
| Sthenurinae | 6 | 28 | 1 | (*49*) |
| Folivora | 62 | 2 | 0.28 | (*50*) |
| Centrarchidae | 19 | 40 | 0.98 | (*51*) |
| Platyrrhini | 33 | 88 | 0.44 | (*52*) |
| Pinnipedia | 89 | 36 | 1 | (*53*) |
| Caninae | 42 | 36 | 1 | (*54*) |
| Equidae | 130 | 8 | 1 | (*55*) |
| Hominini | 22 | 1 | 1 | (*56*) |

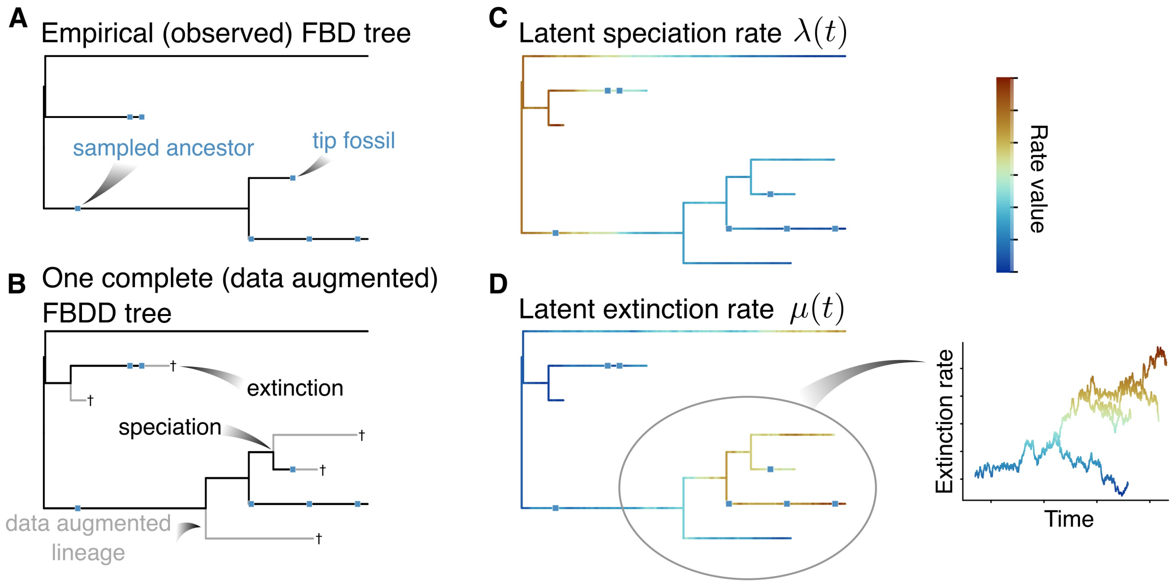


**Supplementary Figure 1: The fossilized birth-death diffusion (FBDD) model with data augmentation (DA) A** An empirical (reconstructed) fossilized birth-death (FBD) tree, with fossils depicted as blue squares. Those fossils that do not have descendants in the empirical tree are labelled 'tip fossils' and the rest are labelled as 'sampled ancestors'. **B** One sample from the posterior of a complete (data augmented) tree, in which unsampled (unobserved because of extinction or undersampling at the present) lineages are probabilistically 'imputed' using Bayesian DA. These 'augmented' lineages are shown in gray while the original empirical tree is left in black (notice that these black lineages are perfectly compatible with the reconstructed tree in A). **C** The latent speciation and **D** the latent extinction rates that follow a Geometric Brownian motion (GBM) from the posterior DA tree as specified by the FBDD (warmer colors represent higher rates). For extinction we show a subsection of the rates in the tree where the latter are now shown as the y axis.

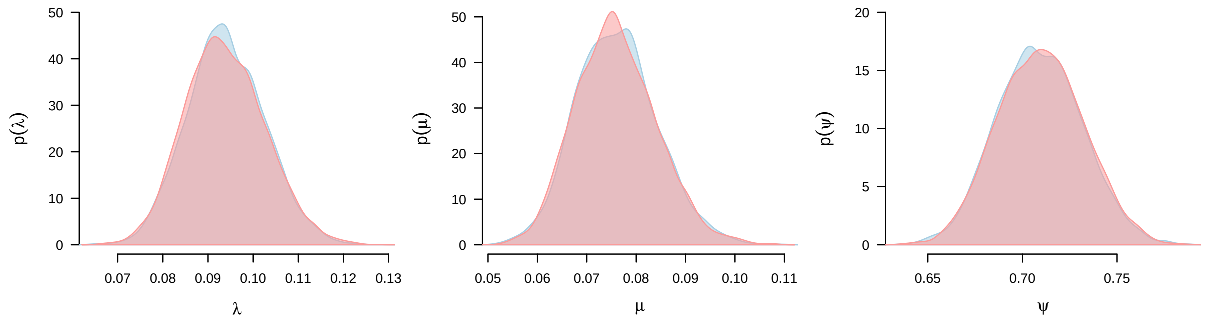


**Supplementary Figure 2: Validation of Bayesian data augmentation inference of the fossilized birth-death (CFBD)** For the same Posterior distributions for: *left* speciation rate ($\lambda$), *middle* extinction rate ($\mu$) and *right* fossilization rate ($\psi$) from the analytical solution of the likelihood using RevBayes (*29*) in blue and from the Bayesian data augmentation algorithm developed in this study in red.

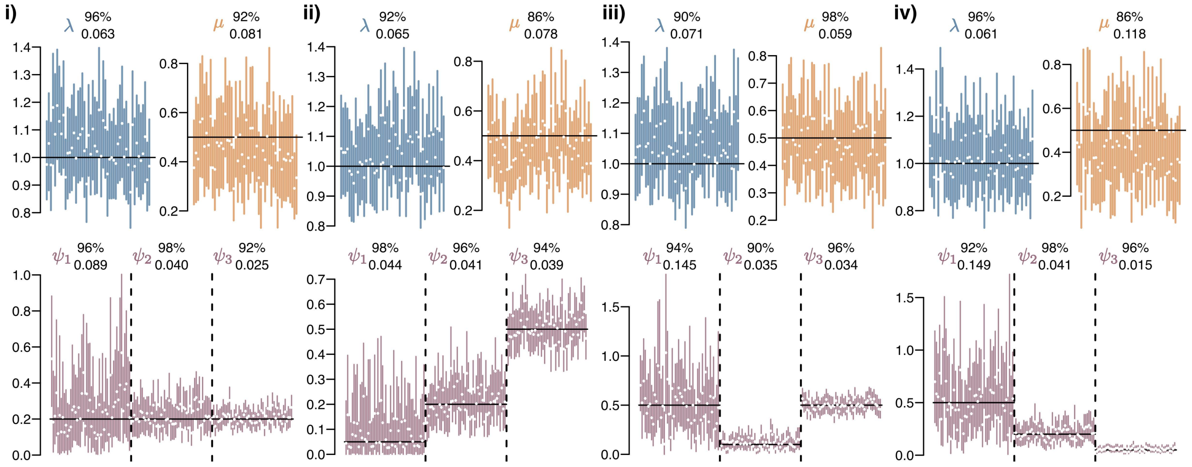


**Supplementary Figure 3: Validation of episodic fossilized birth-death (eFBD)** Results for each of the 4 scenarios (*i*-*iv*, see Supplementary Text) in the simulation study for inference under the eFBD. For each scenario, the top row shows speciation and extinction results, while the bottom row shows the results for fossilization rates across three periods. For each parameter, the black horizontal bar shows the true value, and for each of the 50 replicates vertical colored bars show the 95% Credible Interval (CI) and the white point the mean. The top number percentage represents the coverage (proportion of simulations in which the true value was within the CI) and the bottom number the Mean Absolute Error (MAE).

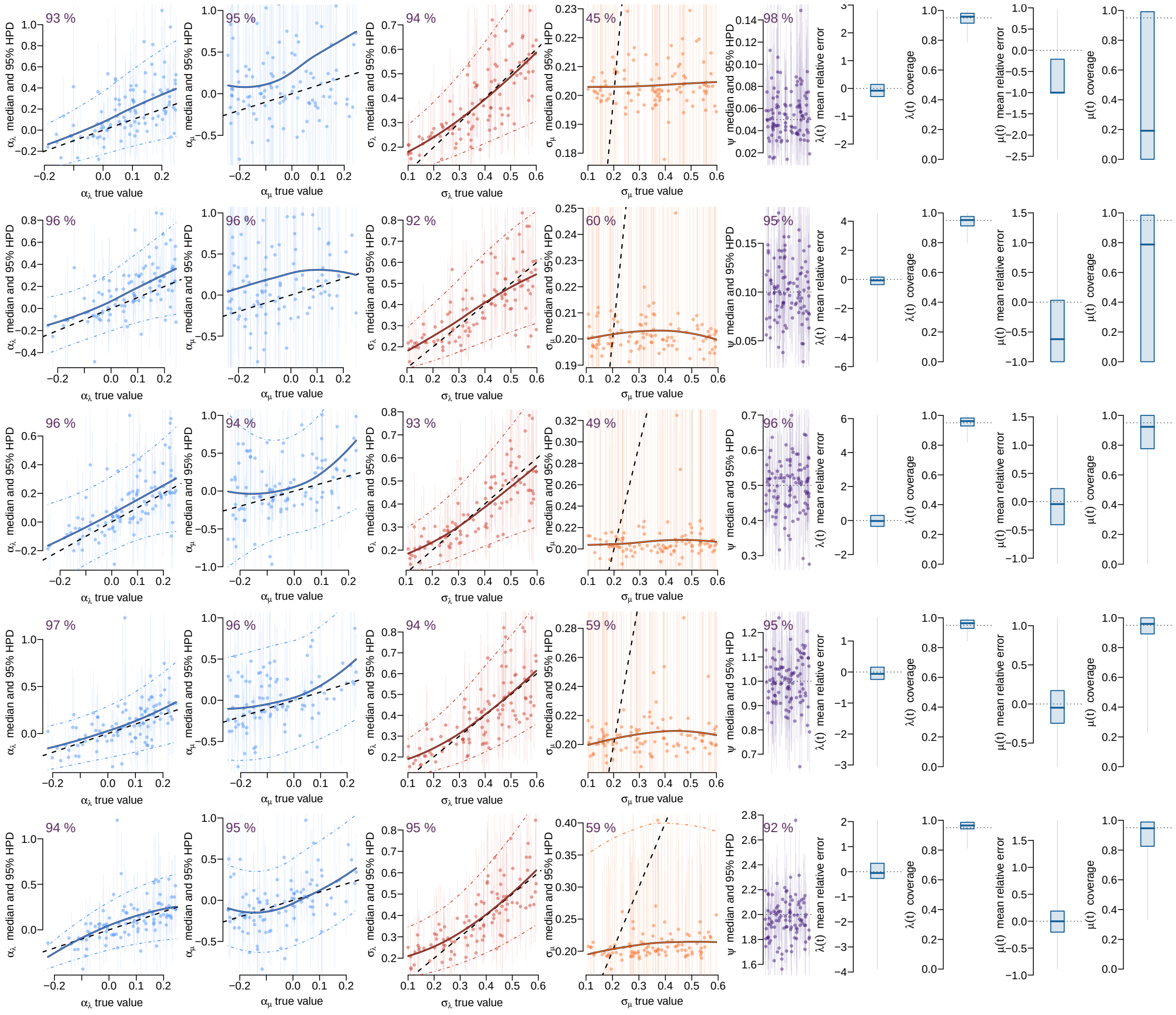

**Supplementary Figure 4: Validation of fossilized birth-death diffusion (FBDD)** Each row represent the results for 5 scenarios that evaluate the impact of increasing rate of fossilization, from top to bottom: $\psi = 0.05$, $\psi = 0.1$, $\psi = 0.5$, $\psi = 1.0$ & $\psi = 2.0$. For each scenario, we sampled from a range of parameter values across $\alpha$, $\sigma_\lambda$ and $\sigma_\mu$ and conducted 100 simulations with subsequent inference under the FBDD. For $\alpha$, $\sigma_\lambda$ and $\sigma_\mu$, the simulated (true) values are shown against their posterior median and their 95% Credible Interval (CI). Colored diagonal solid lines show the loess fit across medians, and the dashed circumscribing ones the 95% Credible Interval (CI). Black dashed lines show the 1:1 line. For $\psi$, the horizontal punctuated line shows the simulated (true) value, and the medians and 95% CI are shown as points and horizontal bars, respectively. Top left percentages represent the statistical coverage. We sampled the distributions of speciation and extinction along the posterior trees and show the mean relative error for speciation $\lambda(t)$ and extinction $\mu(t)$ as well as their statistical coverage.

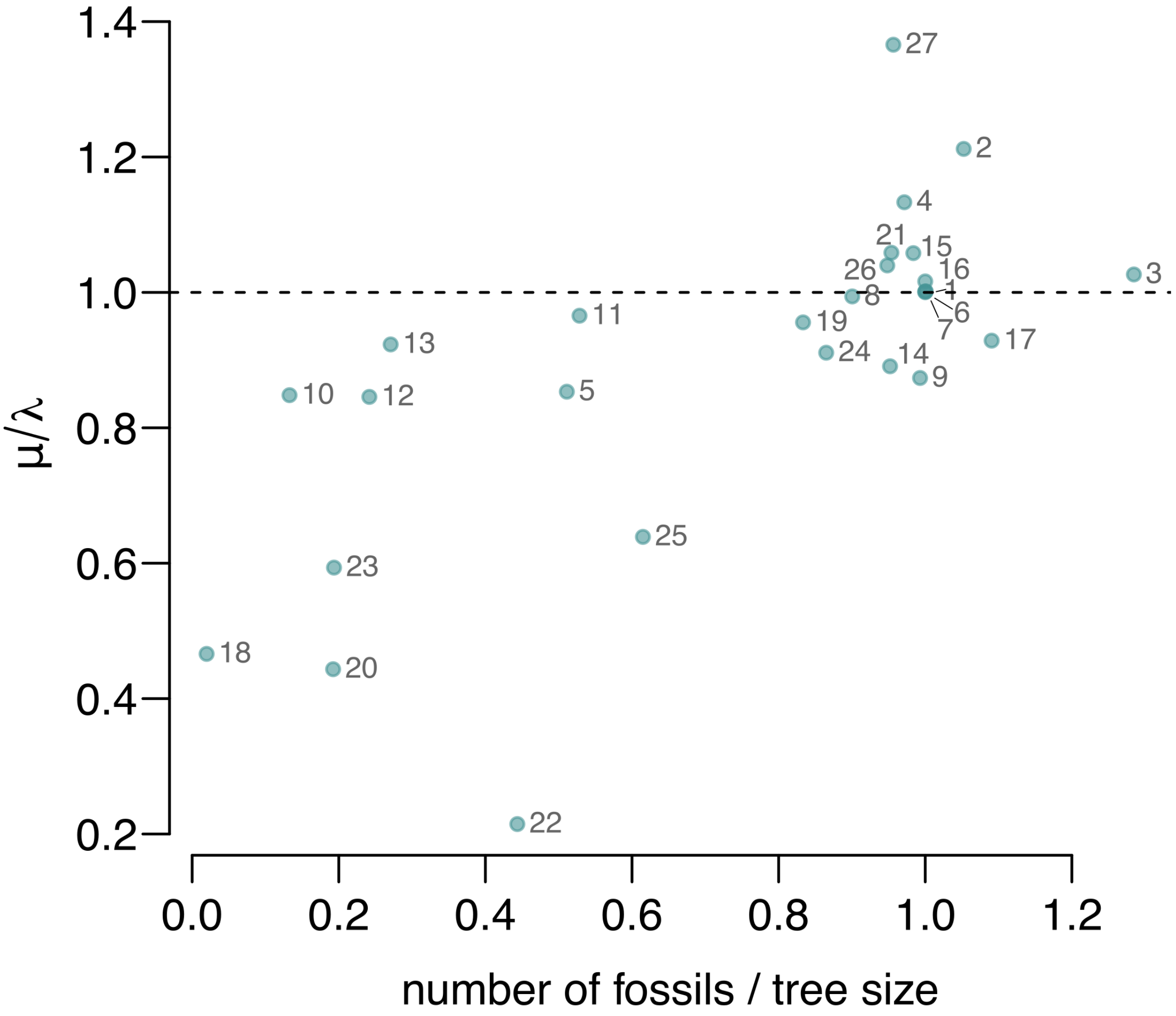


**Supplementary Figure 5: Impact of number of fossils per tree size on turnover rates** Turnover rates (*i.e.*, $\mu/\lambda$) inferred under the eFBD against the number of fossils per tree size (number of tips, both extant or extinct) for the 27 taxa studied here. With even a low percentage of fossils per tree size we already recover high rates of turnover. Enumeration of taxa, in order of appearance: 1) eurypterida, 2) dipnoi, 3) therocephalia, 4) sphenodontidae, 5) testudinata, 6) pterosauria, 7) dinosauria, 8) crocodylomorpha, 9) juglandaceae, 10) tetraodontiformes, 11) mauritiinae, 12) acanthuridae, 13) siganidae, 14) spheniscidae, 15) proboscidea, 16) brontotheriioidea, 17) cetacea, 18) zoarcoidei, 19) ruminantia, 20) sthenurinae, 21) folivora, 22) centrarchidae, 23) platyrrhines, 24) pinnipedia, 25) caninae, 26) equidae, 27) hominini.

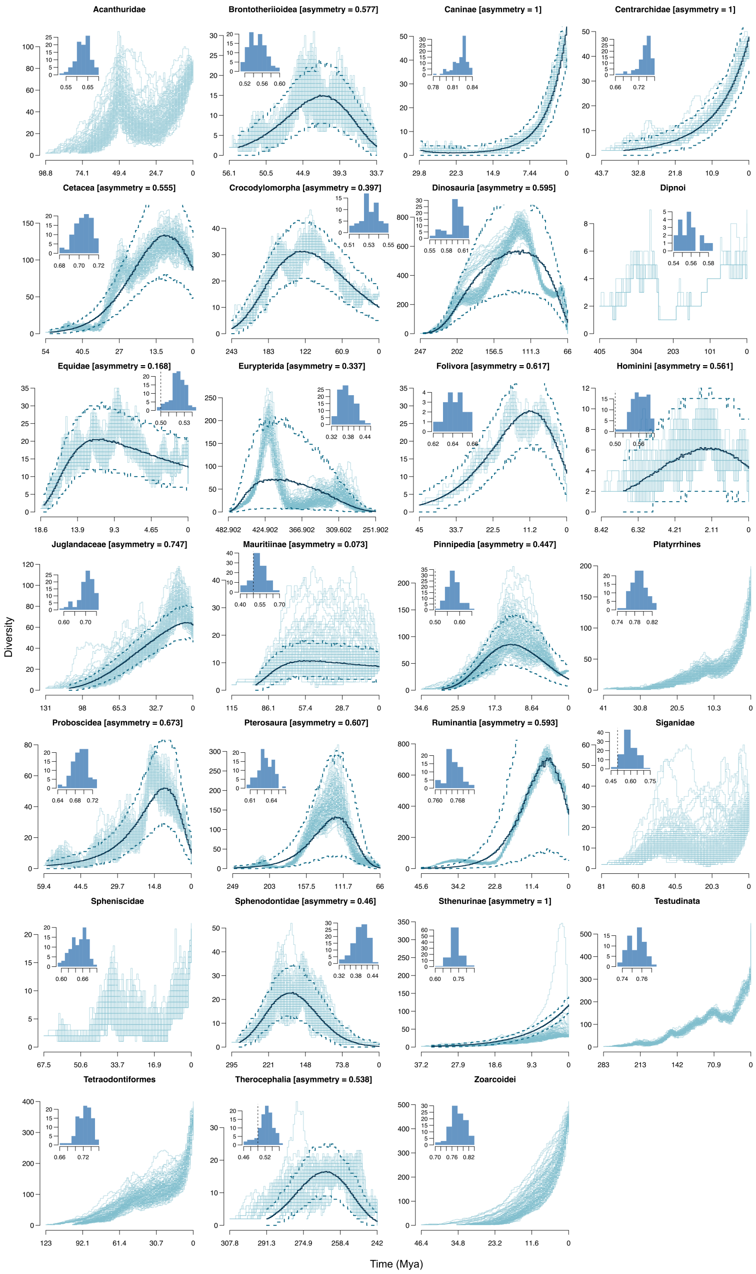


**Supplementary Figure 6: Diversity trajectories for the** 27 **evolutionary radiations studied** For each clade, 100 diversity trajectories are shown from a posterior sample of 100 complete trees. The asymmetry measured with the parametric approach (Materials and Methods) is given next to the clade name for clades in decline or fall, and the inset panel shows the distribution of Center of Gravity (CG; Materials and Methods) across the 100 trees. Caninae, Centrarchidae, and Sthenurinae are in decline, with larger diversity before the present, but because this past peak was very close to the present, the parametric model was not able to pick up a subsequent downward trend, rather just assuming is part of the expected variance. These three clades correspond then to the three points with asymmetry of 1 in main text Fig 1.

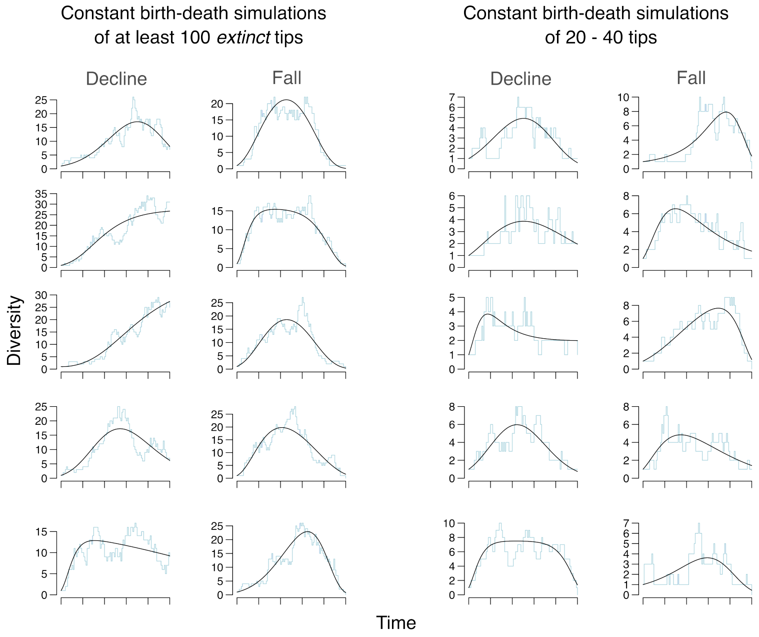


**Supplementary Figure 7: samples of diversity trajectories from constant birth-death simulations.** Examples of the 800 total simulated diversity trajectories using a constant-birth death model with turnover ($\mu/\lambda$) of 1, using $\lambda = 1$ and $\mu = 1$ over 10 time units. The first 400 simulations (left) were conditioned on at least having 100 extinct tips, with 200 trajectories of clades in decline and 200 trajectories for clades that went extinct (fall). The second set fo 400 simulations (right) were conditioned on clades that had from 20 to 40 tips, with 200 trajectories for clades in decline and 200 for clades that went extinct. 5 examples from each of the 200 simulations of the 4 categories above are shown.

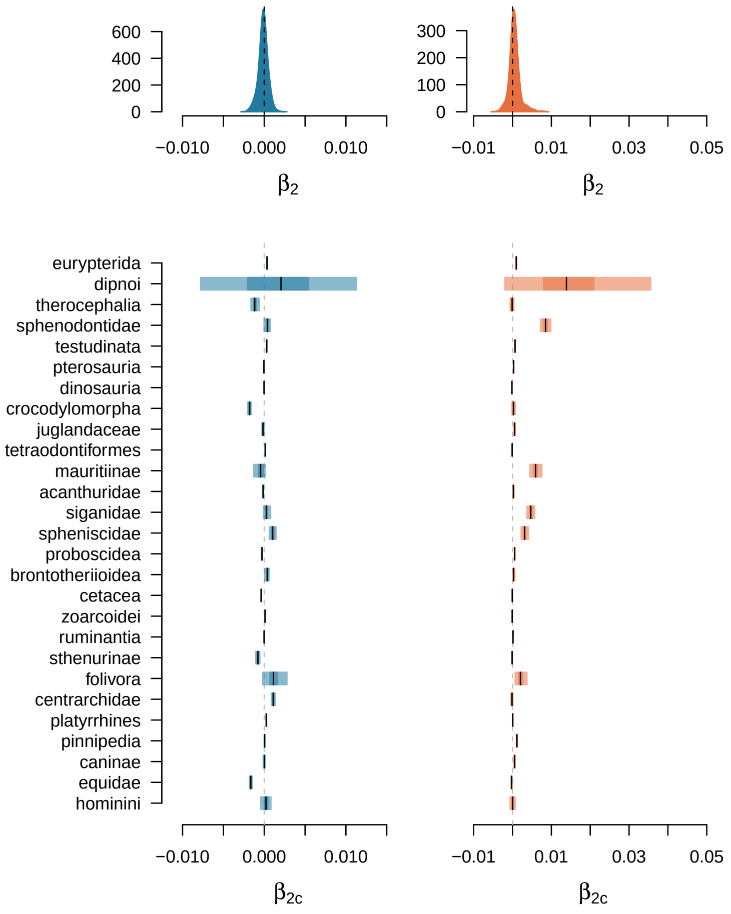


**Supplementary Figure 8: No effect of diversity on average speciation and extinction rates** Results from the three level hierarchical auto-regressive model testing for an effect of diversity on average speciation or extinction rates (Materials and Methods). The top row shows the global effects ($\beta_2$) *left* for speciation in blue and *right* for extinction in red. Note that the 95% Credible Interval (CI) contains 0. The bottom row shows the clade specific effects of diversity on *left* speciation and *right* extinction. Clades are shown in order of first appearance.

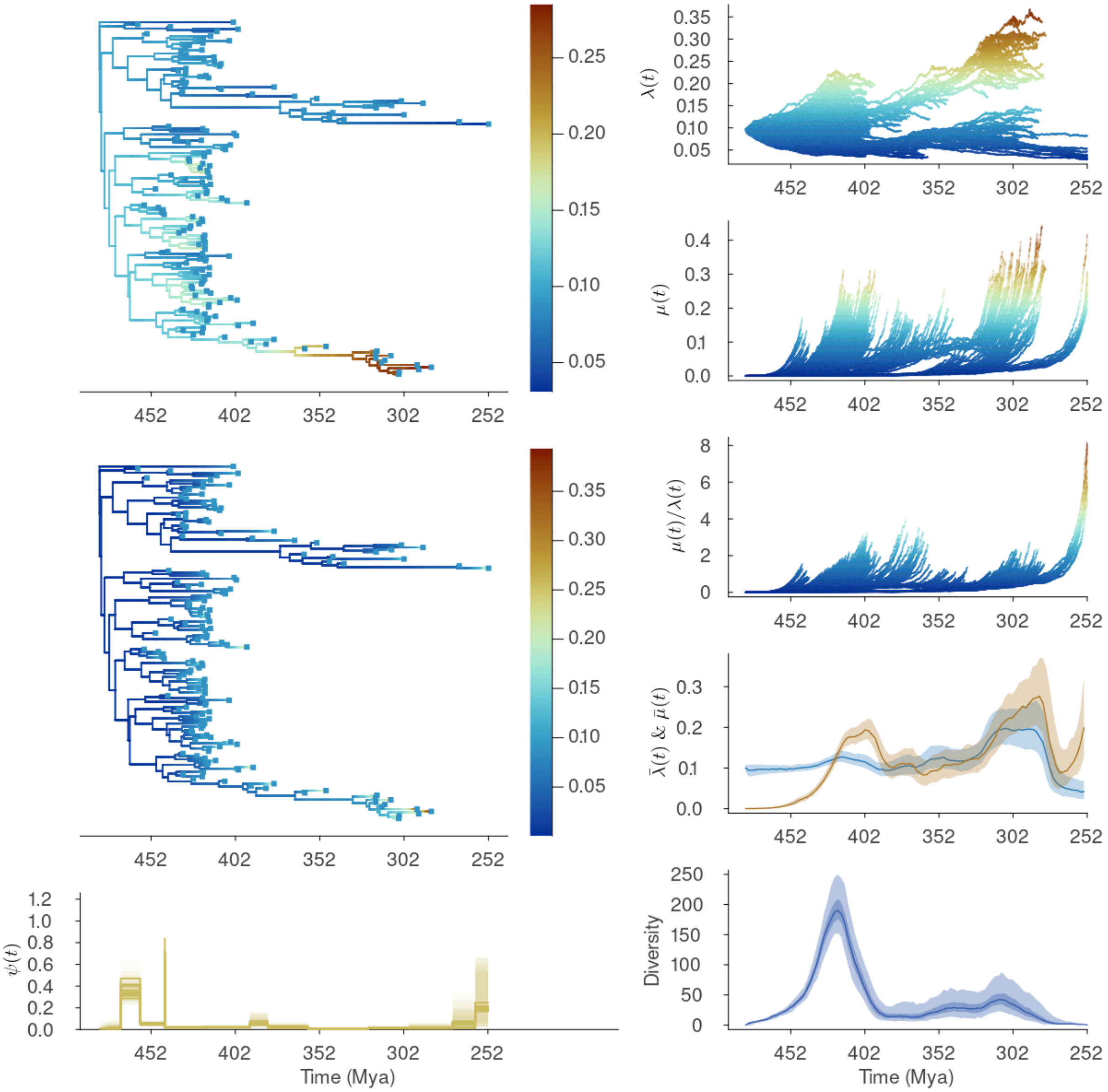


**Supplementary Figure 9: FBDD results for Eurypterida.** *Left* column shows: *top*, the posterior average rates for speciation and *middle* for extinction rates on the reconstructed tree of one of the empirical tree samples (Note that marginal posterior distributions for rates can only be estimated for the observed (reconstructed) part of the tree (*i.e.*, $\Psi_o$)). *bottom* shows the posterior fossilization rates estimates through time for each of the empirical tree samples. *Right* column from *top* to *bottom* shows: marginal posterior median rates of speciation and of extinction across the empirical tree samples (again, these two can only be estimated on the observed part of the tree), posterior cross-lineage average speciation ($\bar{\lambda}(t)$ in blue) and extinction ($\bar{\lambda}(t)$ in red) rates, and posterior cross-lineage variance in speciation ($V(\lambda(t))$ in blue) and extinction ($V(\lambda)(t)$ in red) rates across the complete trees (average and 95% Credible Interval), and, finally, the median, first quartile and 95% CI of estimated diversity trajectories.

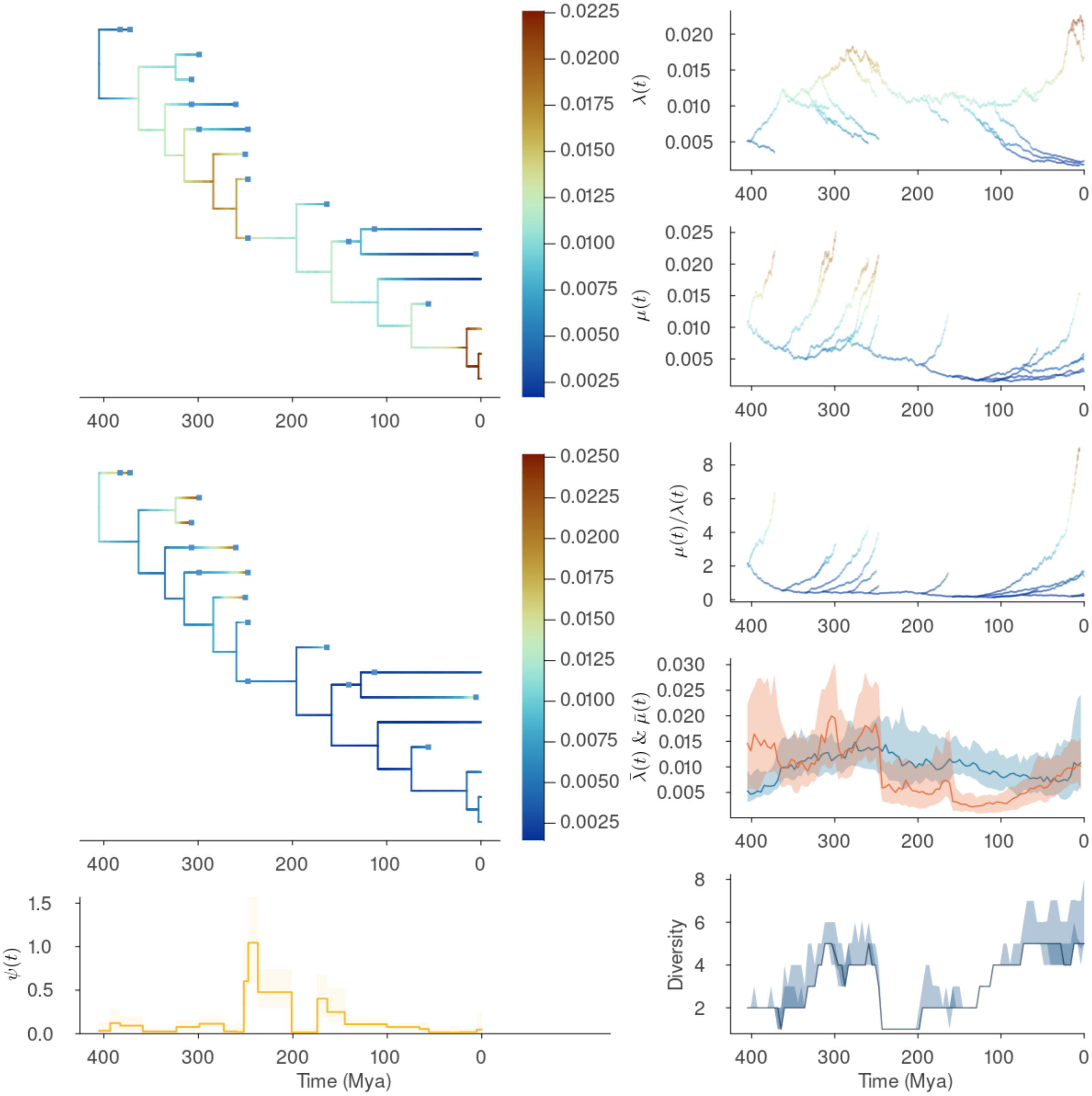


**Supplementary Figure 10: FBDD results for Dipnoi.** Details as in Fig. S9.

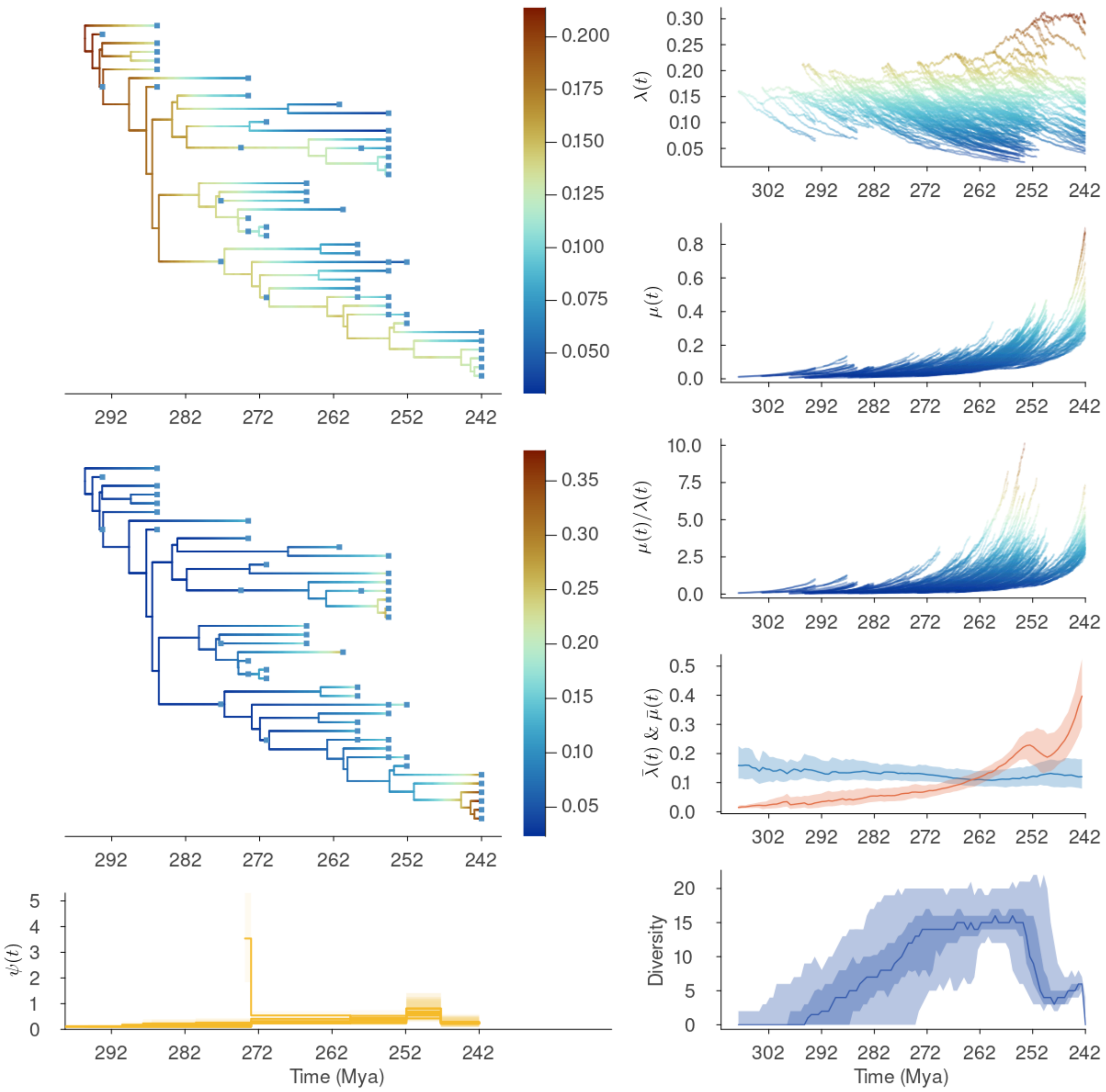


**Supplementary Figure 11: FBDD results for Therocephalia.** Details as in Fig. S9.

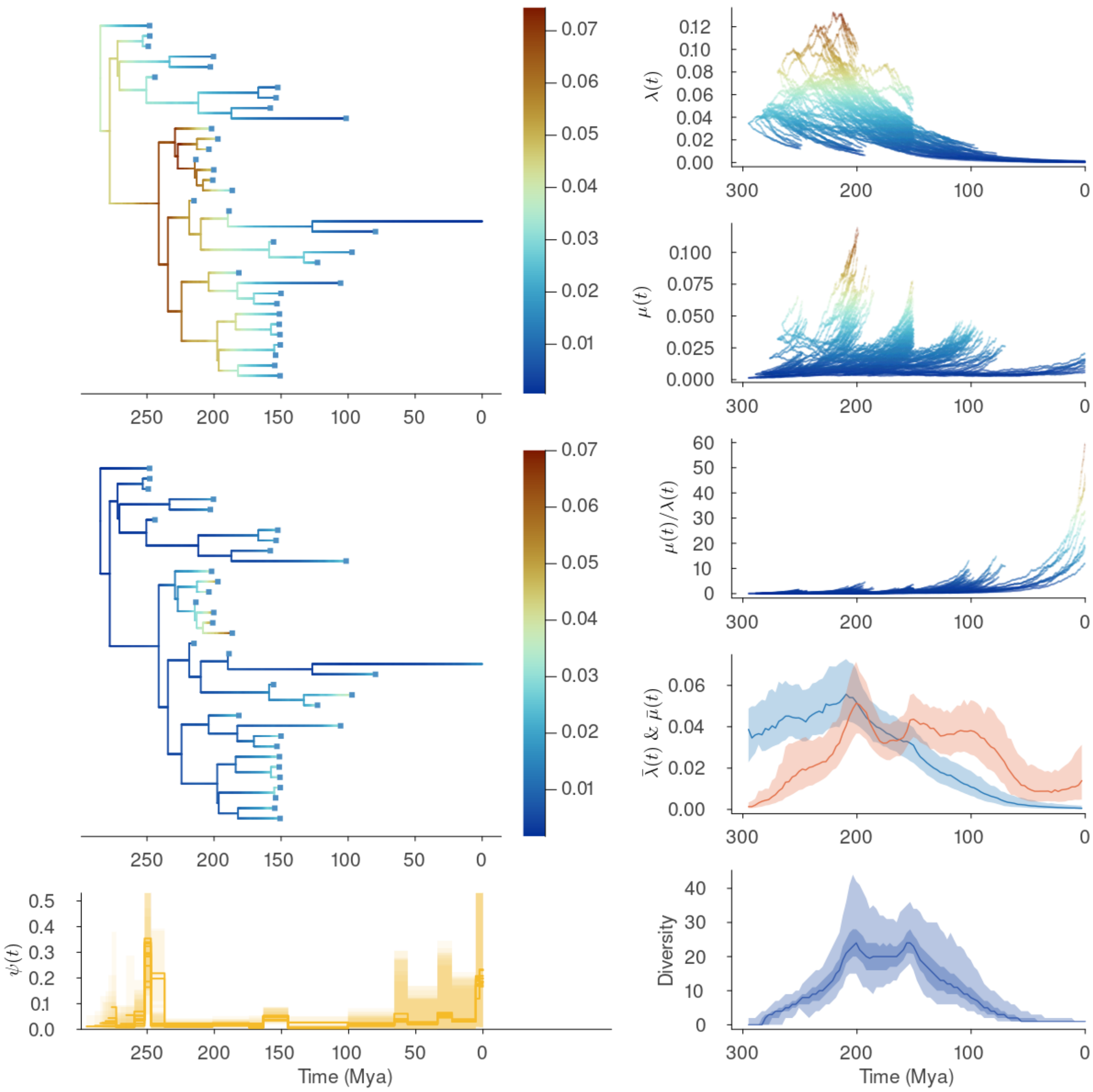


**Supplementary Figure 12: FBDD results for Sphenodontidae.** Details as in Fig. S9.

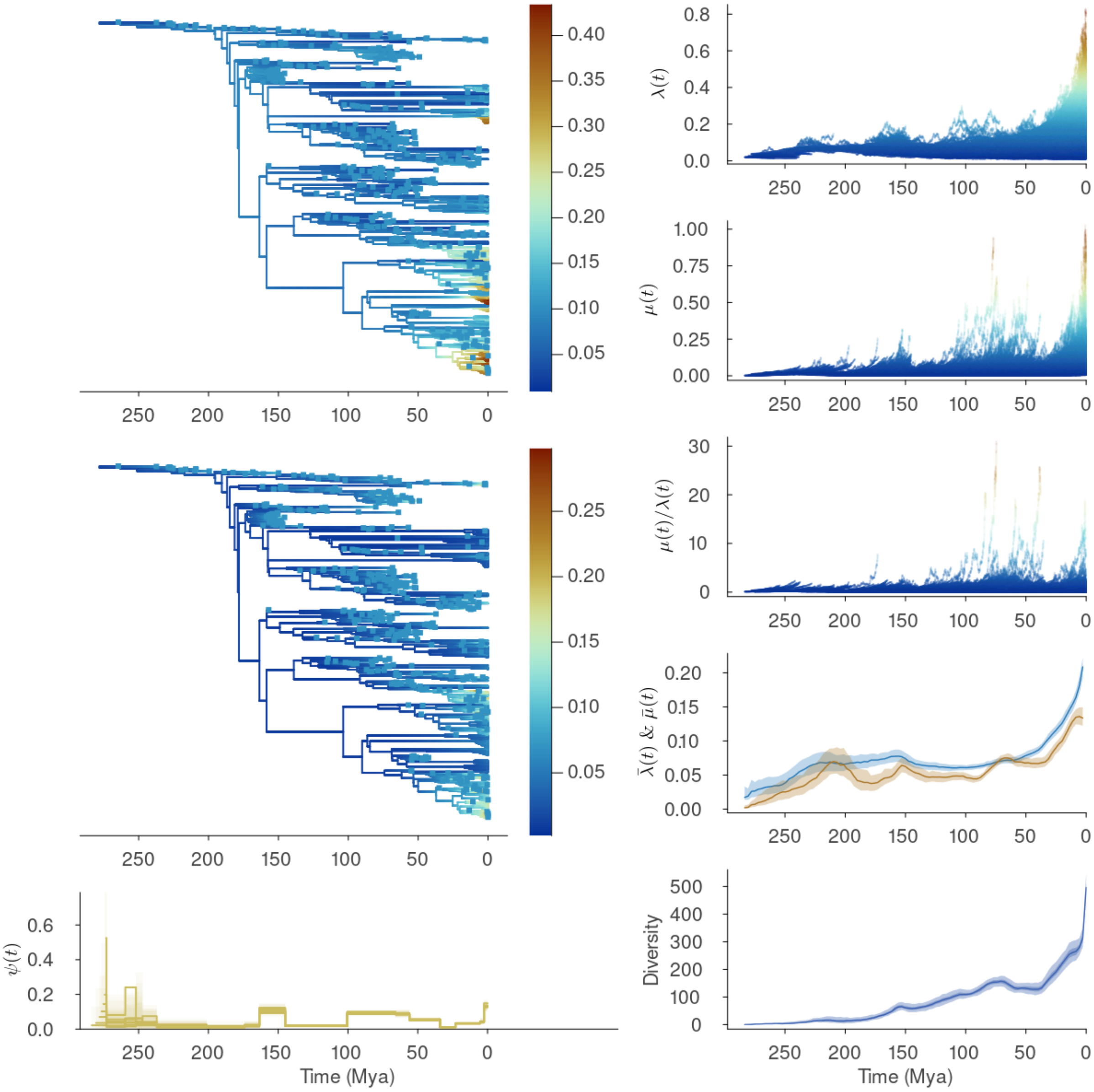


**Supplementary Figure 13: FBDD results for Testudinata.** Details as in Fig. S9.

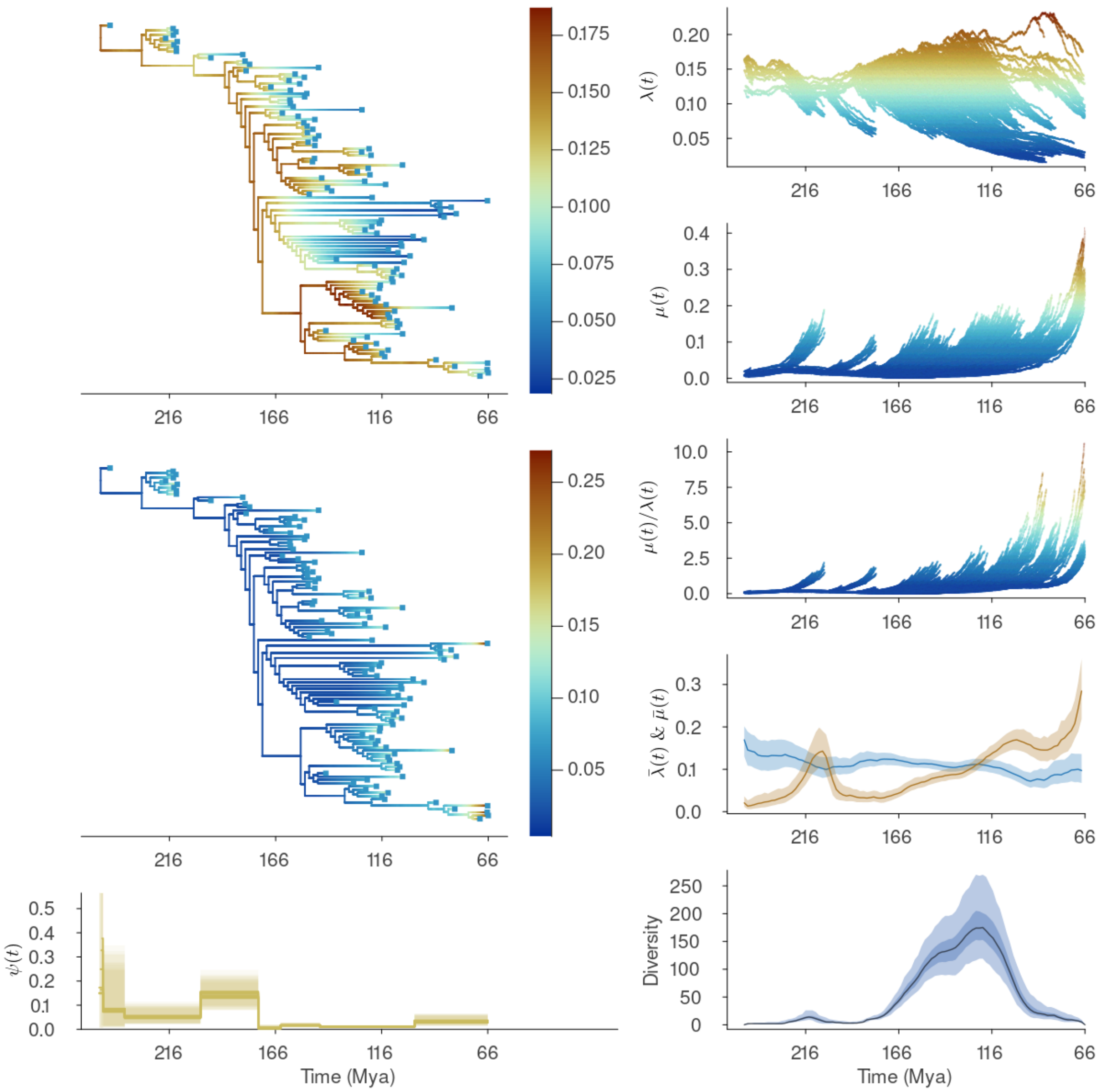


**Supplementary Figure 14: FBDD results for Pterosauria.** Details as in Fig. S9.

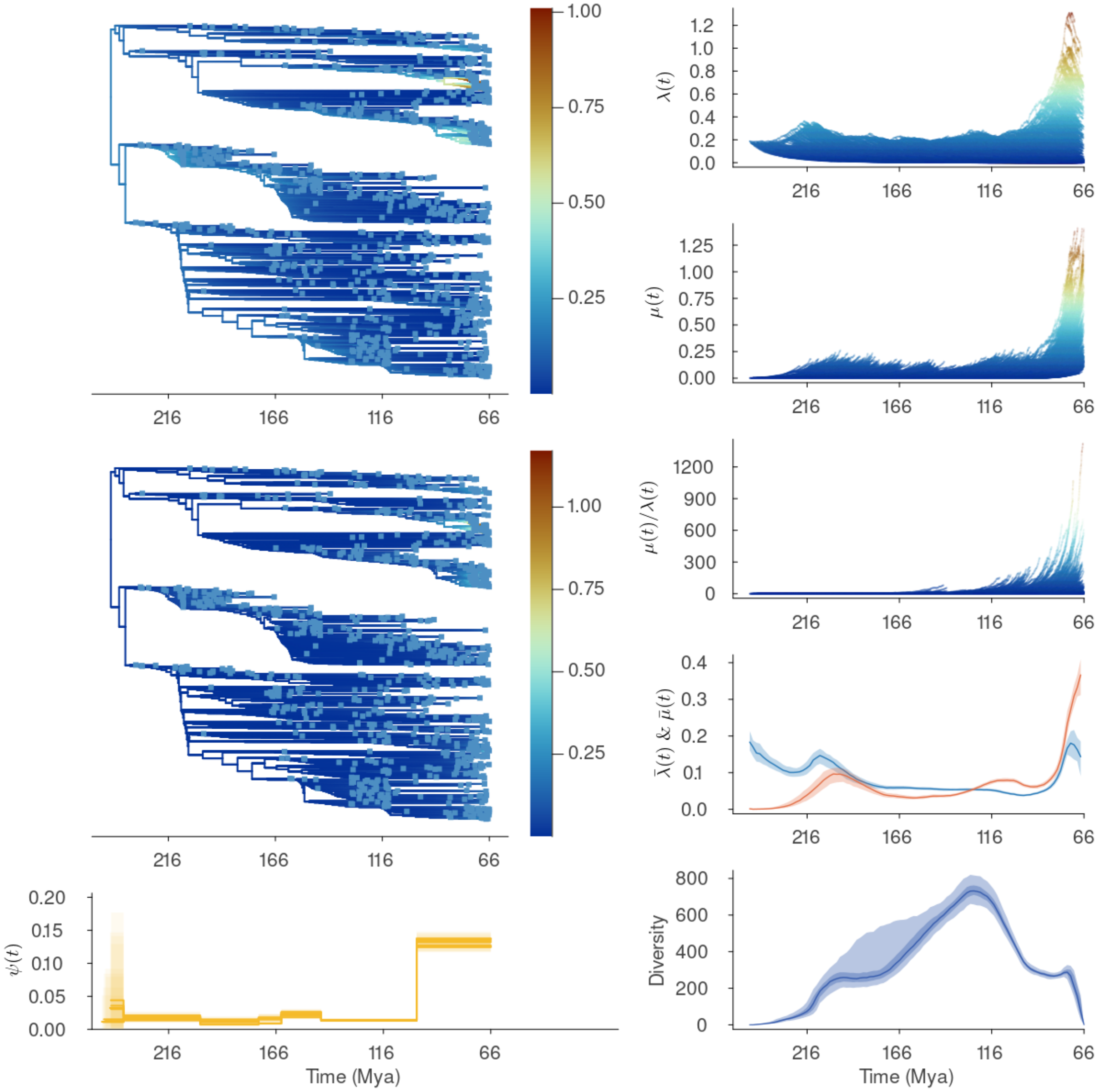


**Supplementary Figure 15: FBDD results for Dinosauria.** Details as in Fig. S9.

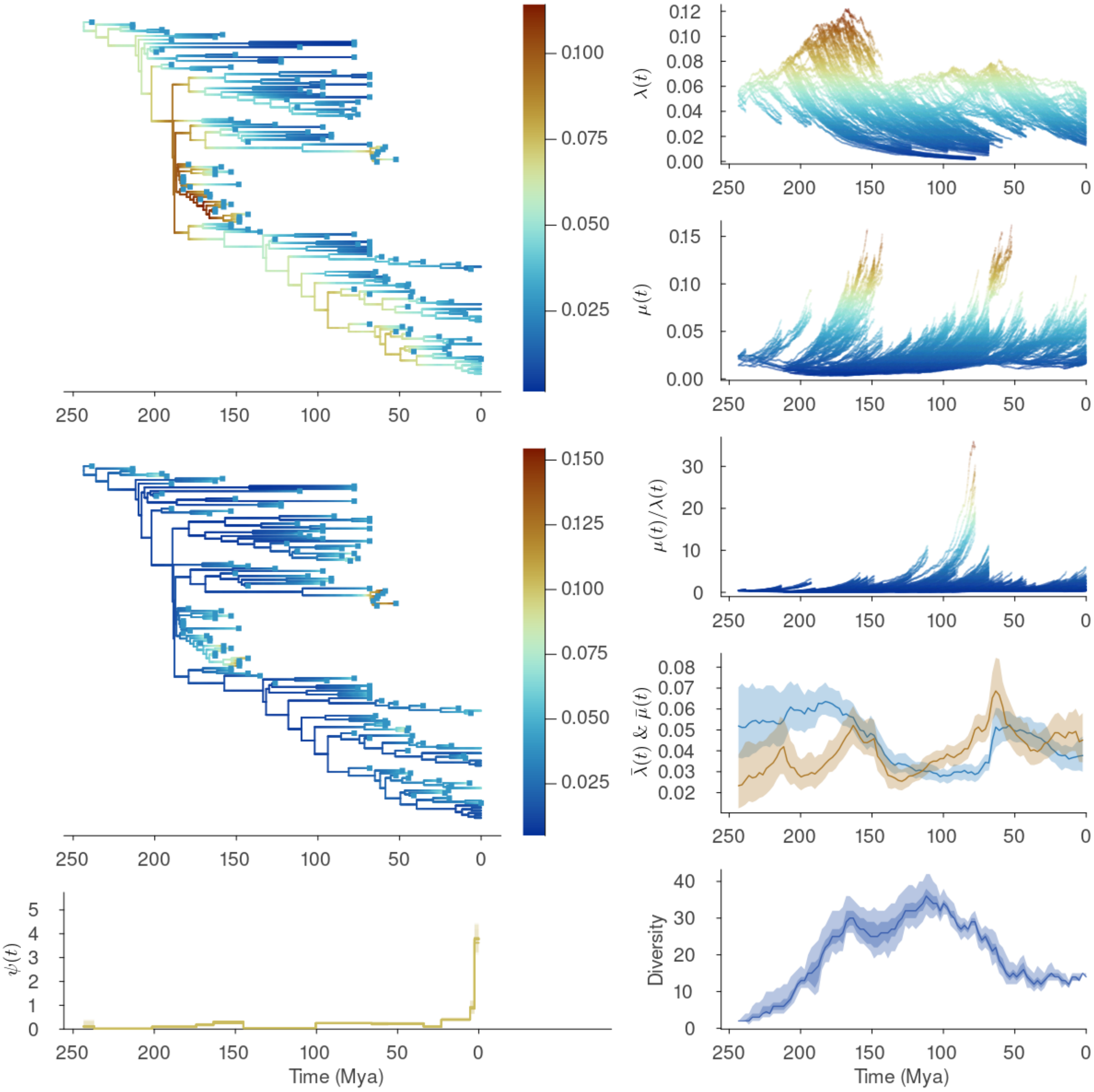


**Supplementary Figure 16: FBDD results for Crocodylomorpha.** Details as in Fig. S9.

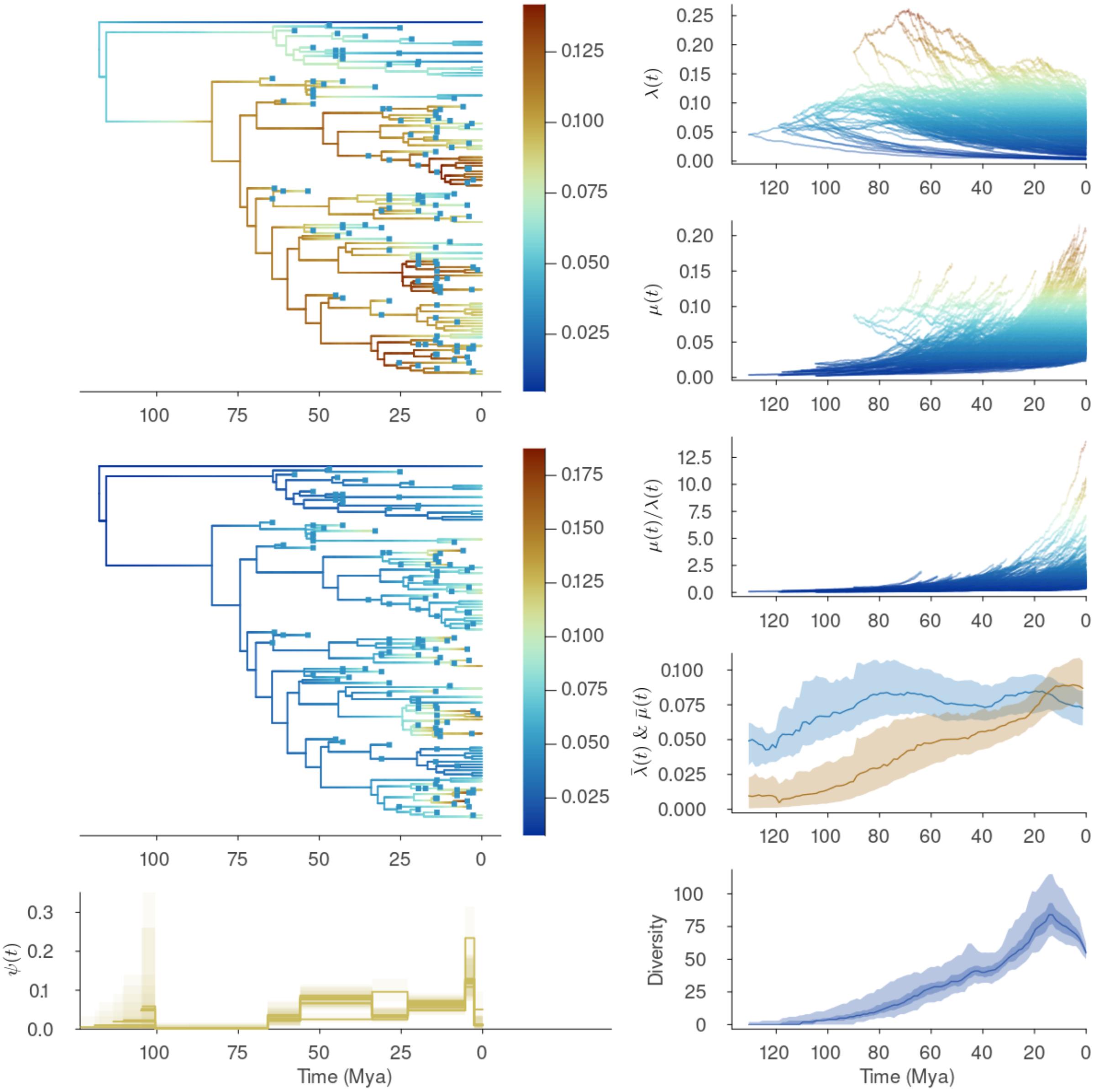


**Supplementary Figure 17: FBDD results for Juglandaceae.** Details as in Fig. S9.

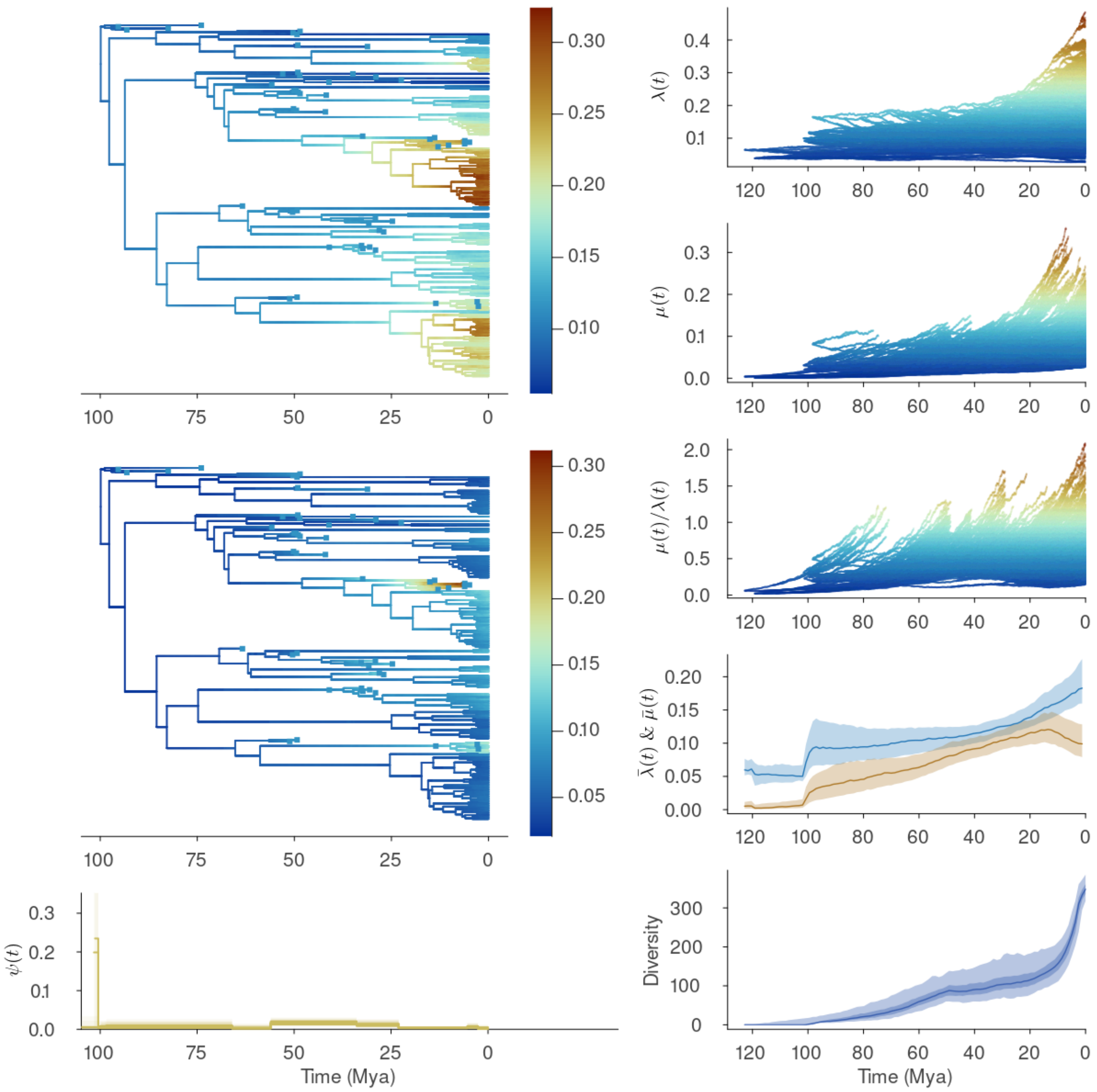


**Supplementary Figure 18: FBDD results for Tetraodontiformes.** Details as in Fig. S9.

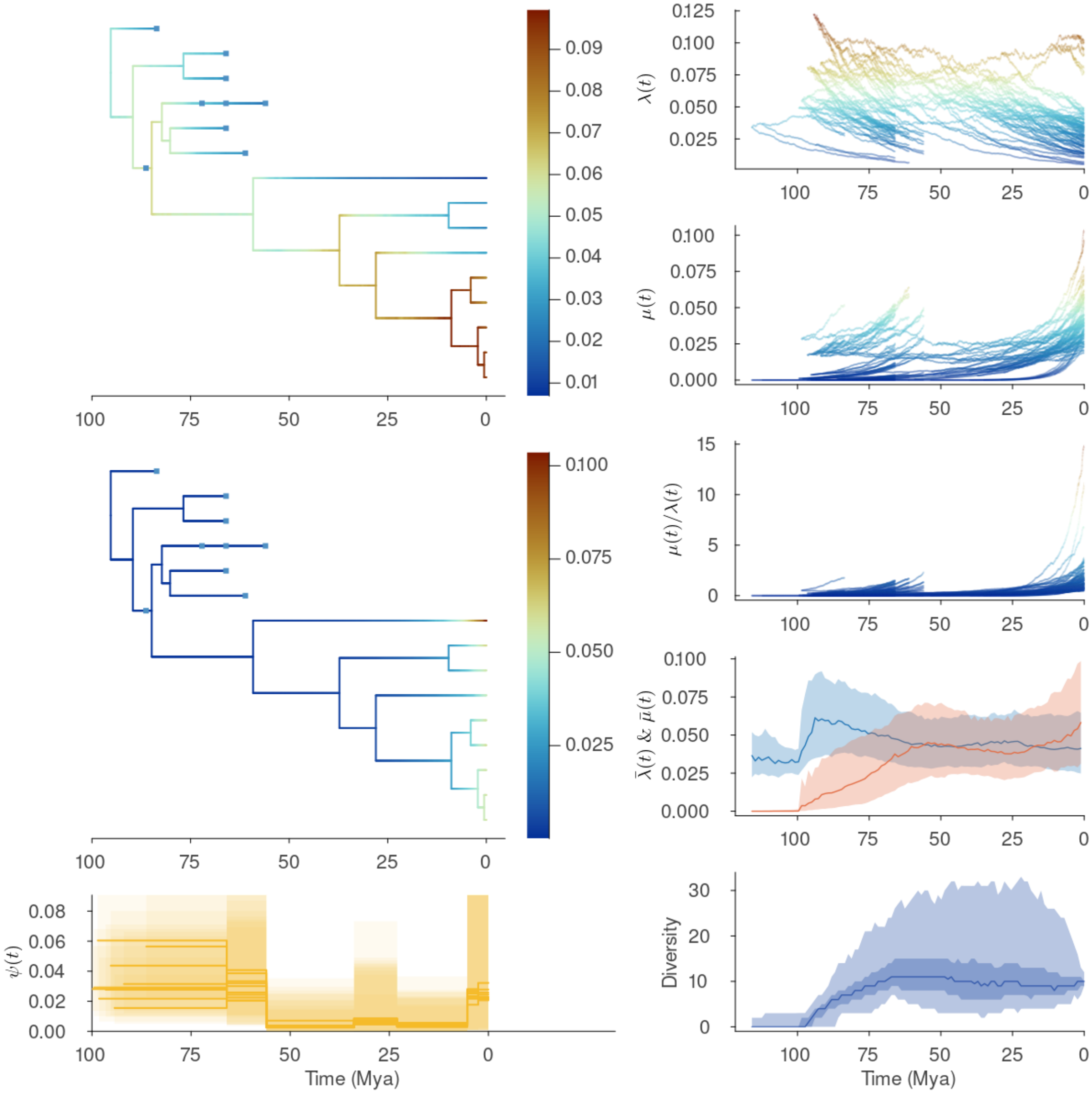


**Supplementary Figure 19: FBDD results for Mauritiinae.** Details as in Fig. S9.

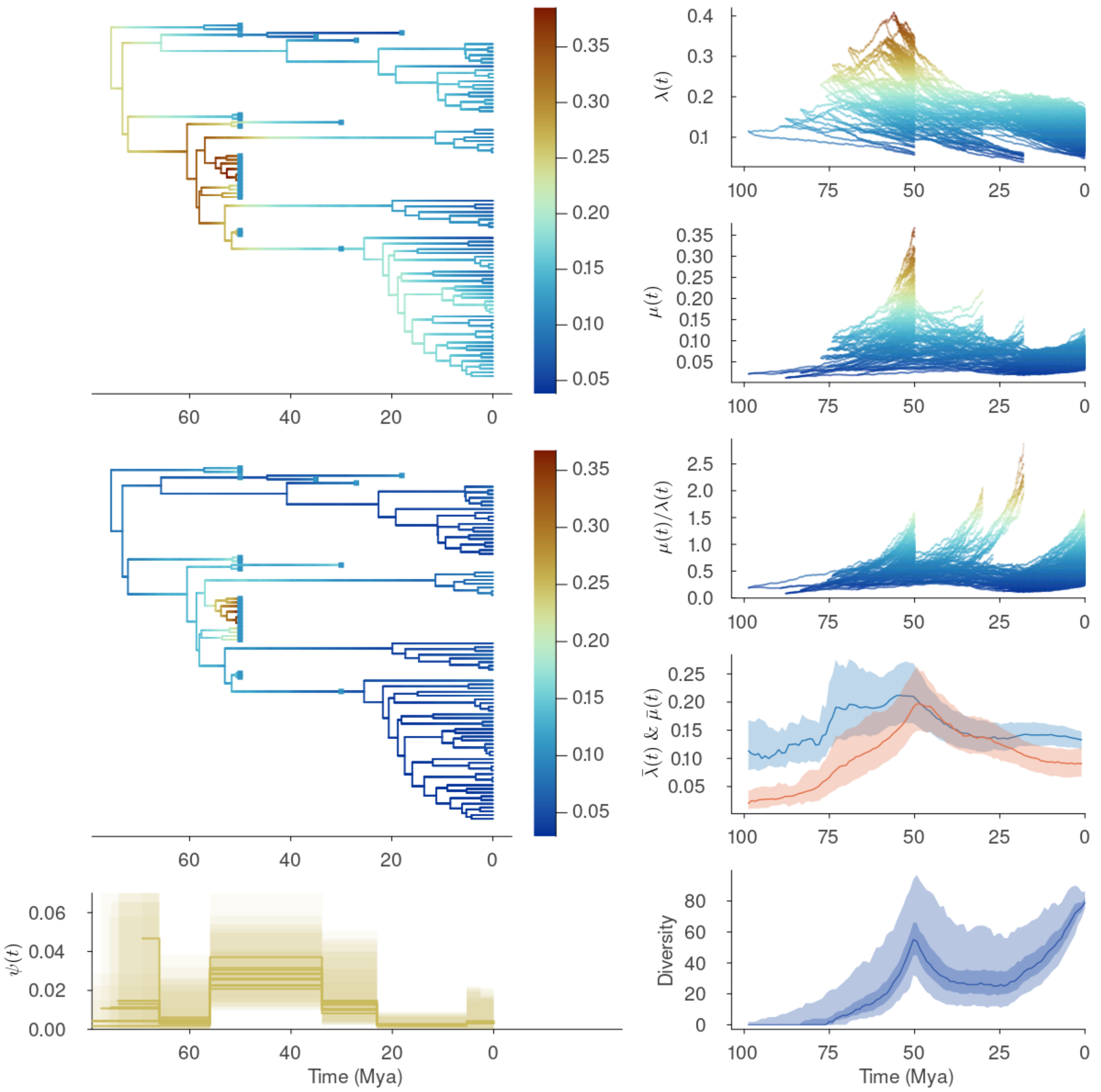


**Supplementary Figure 20: FBDD results for Acanthuridae.** Details as in Fig. S9.

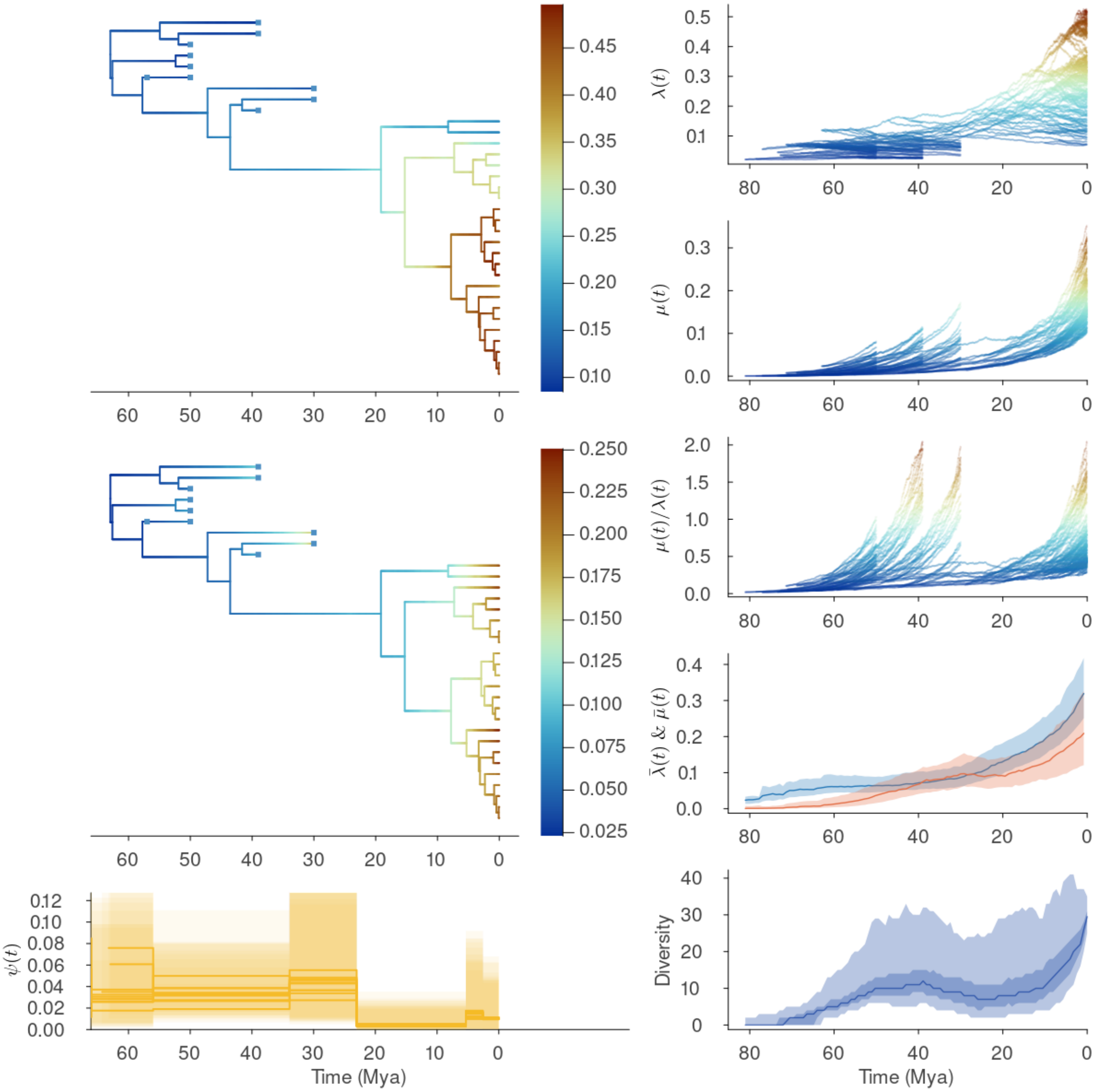


**Supplementary Figure 21: FBDD results for Siganidae.** Details as in Fig. S9.

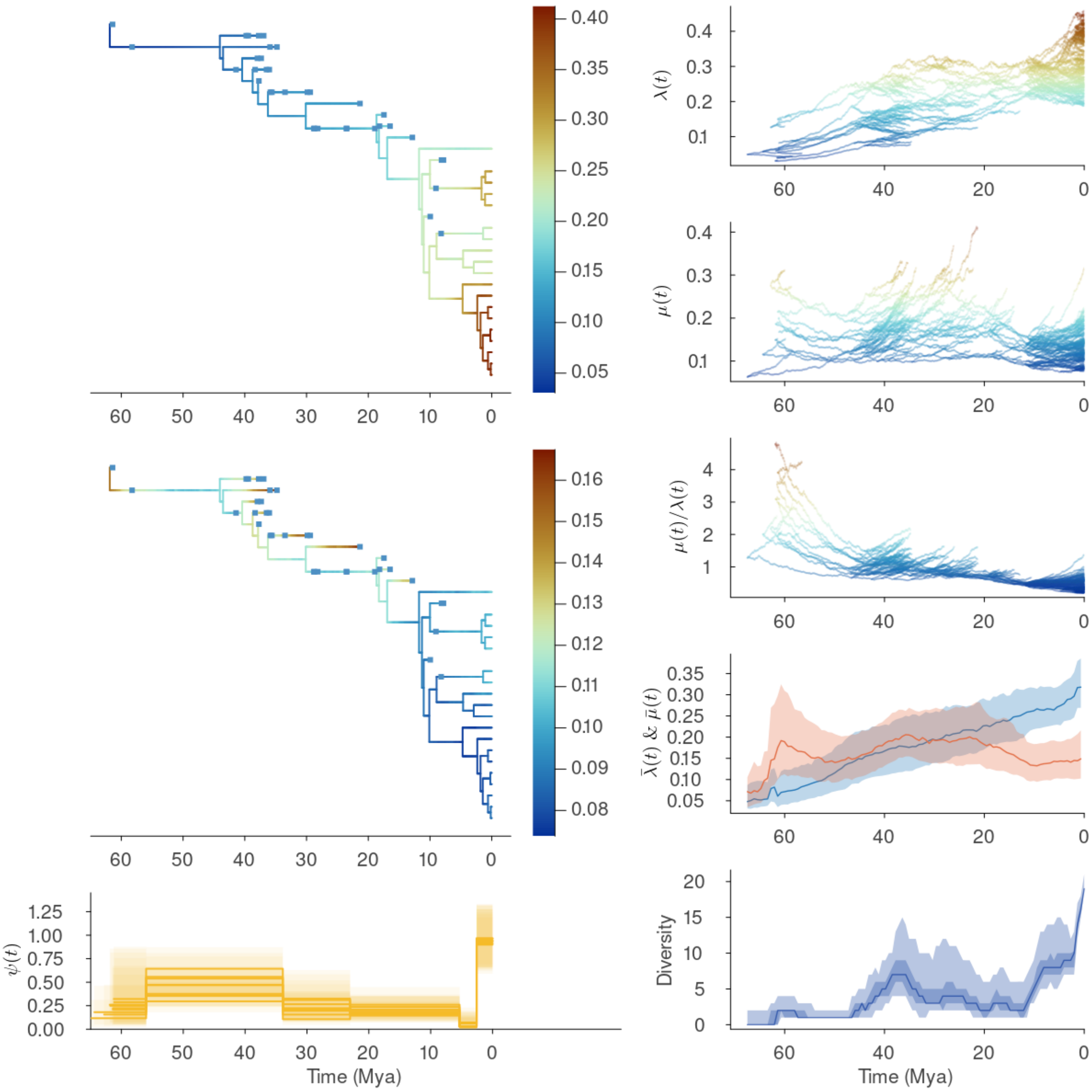


**Supplementary Figure 22: FBDD results for Spheniscidae.** Details as in Fig. S9.

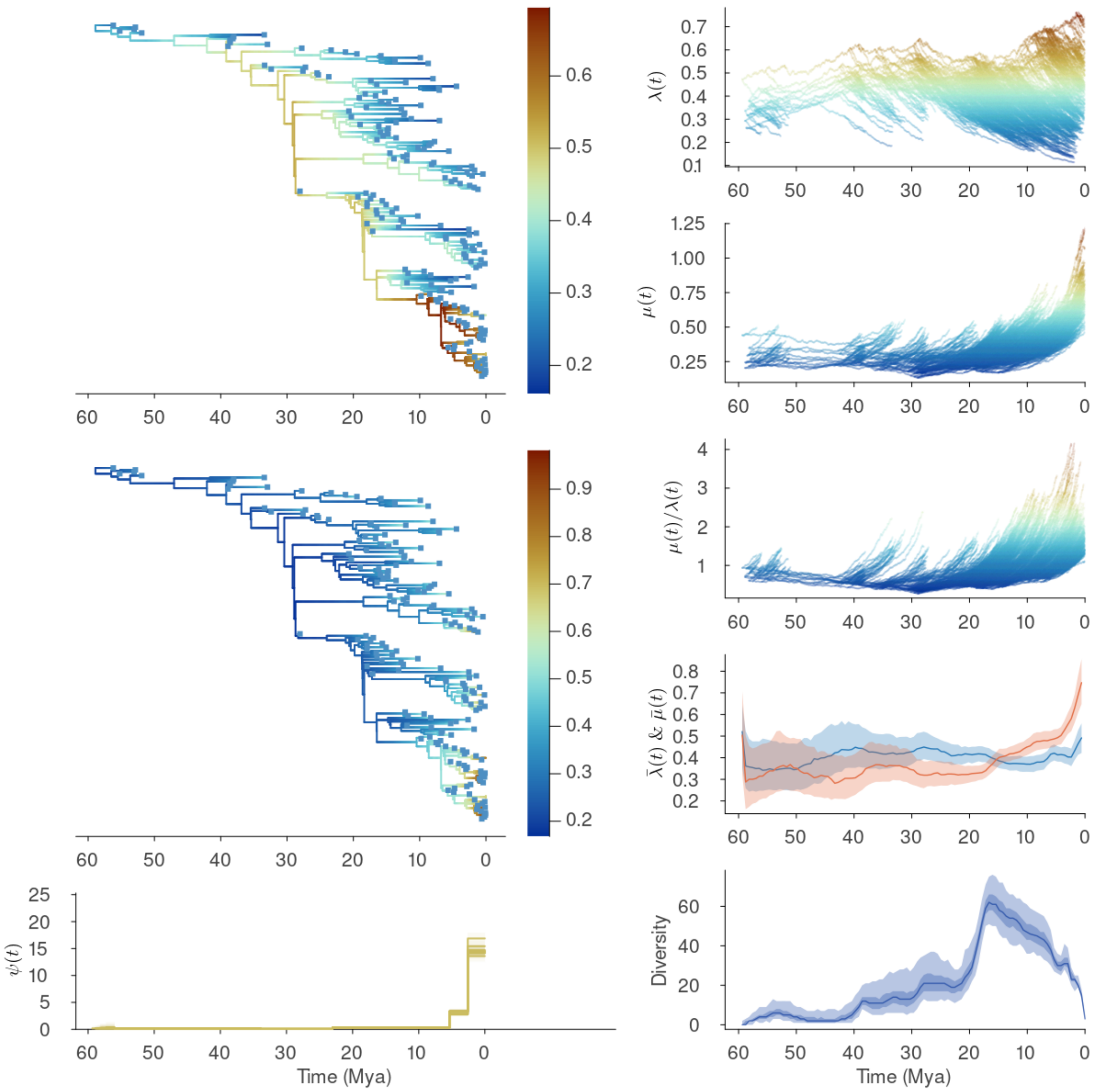


**Supplementary Figure 23: FBDD results for Proboscidea.** Details as in Fig. S9.

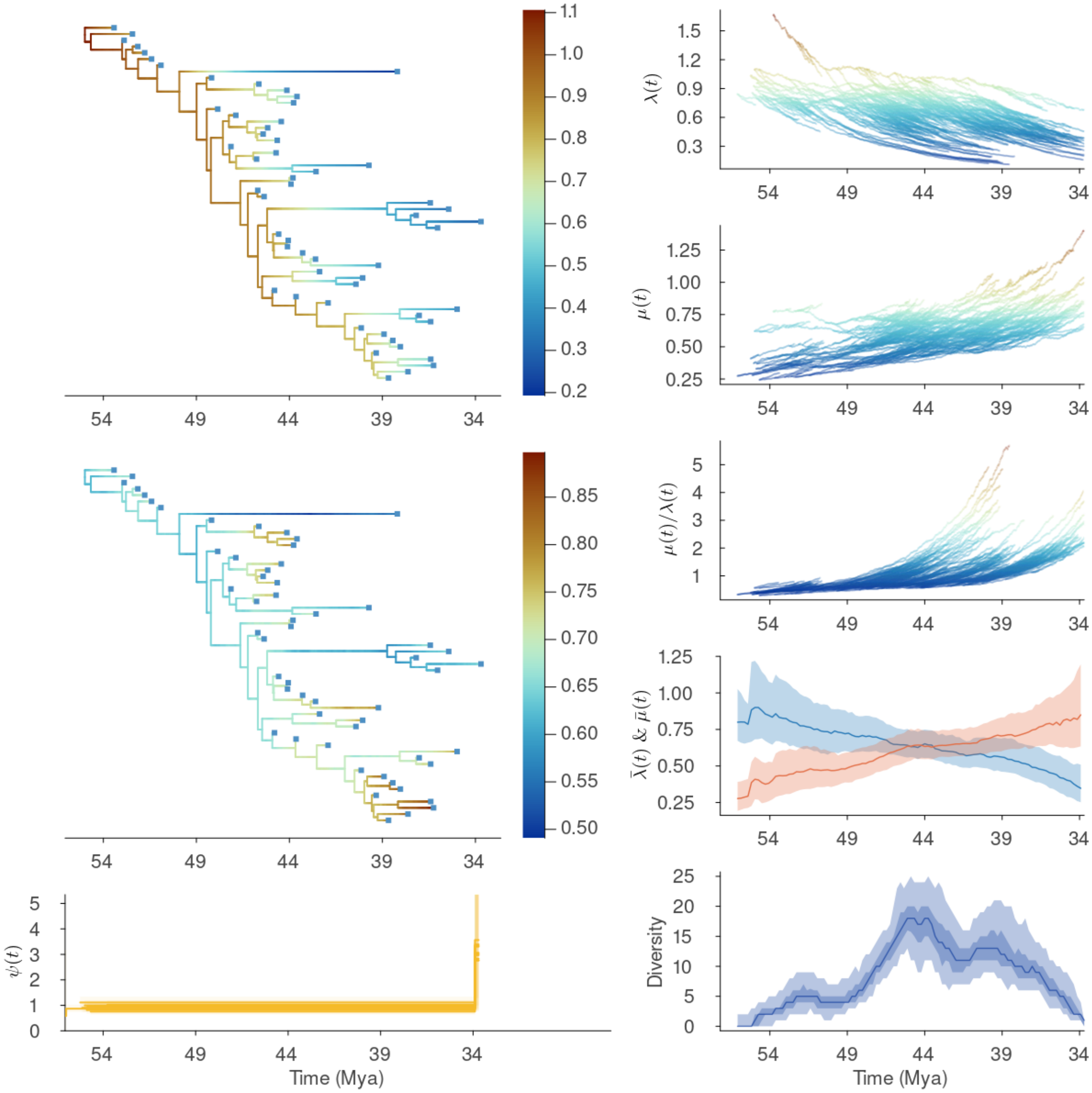


**Supplementary Figure 24: FBDD results for Brontotheriioidea.** Details as in Fig. S9.

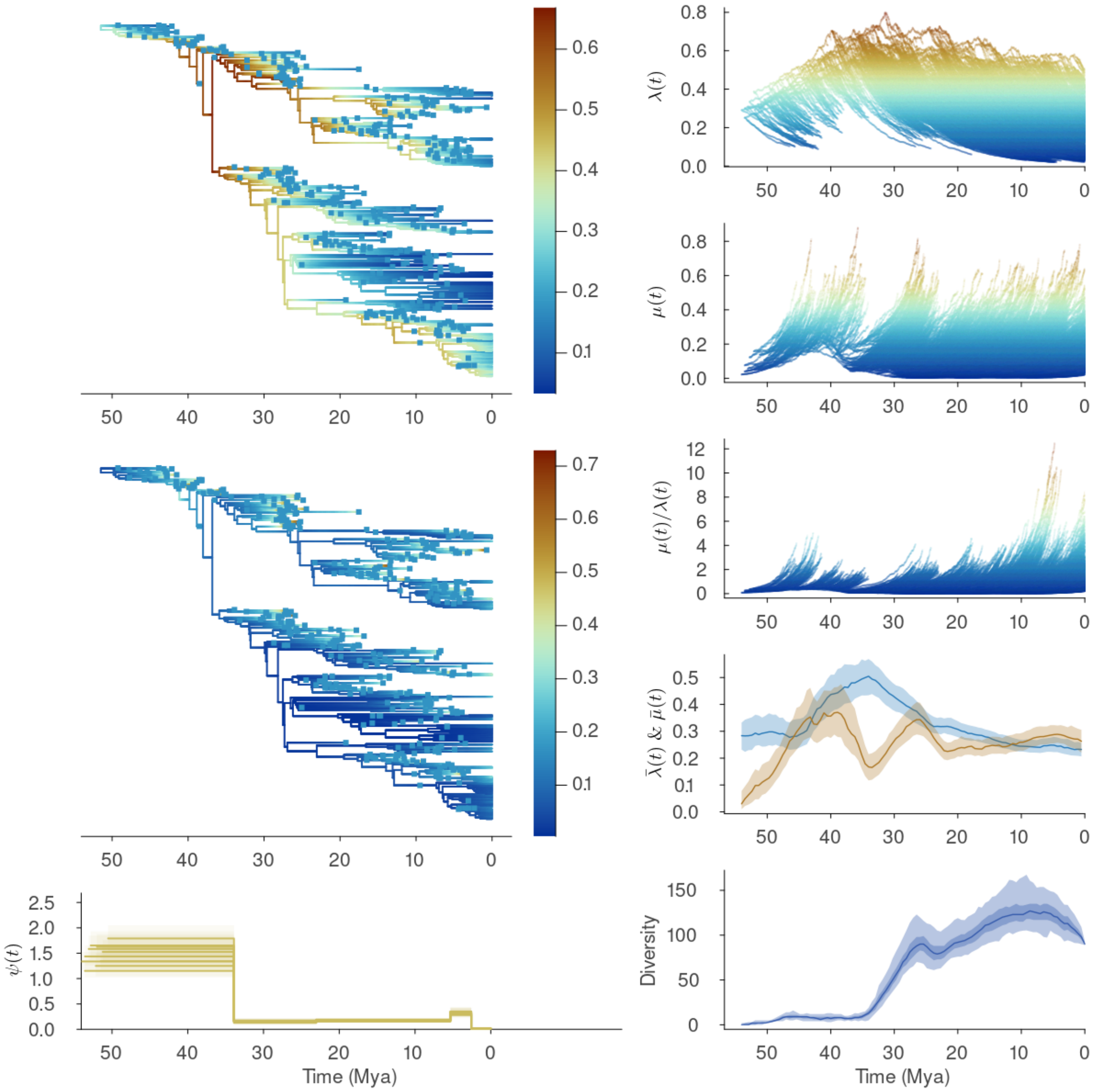


**Supplementary Figure 25: FBDD results for Cetacea.** Details as in Fig. S9.

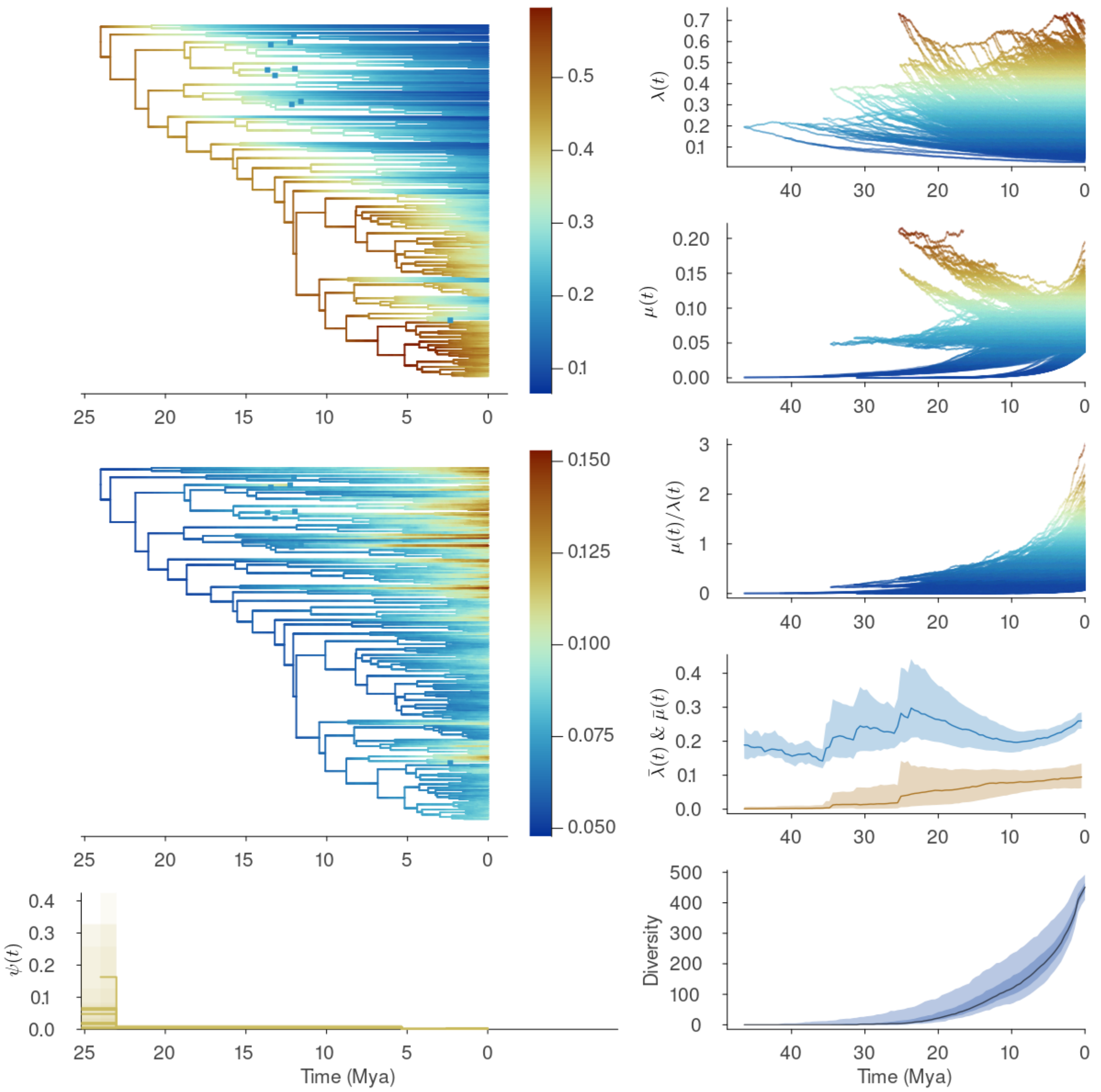


**Supplementary Figure 26: FBDD results for Zoarcoidei.** Details as in Fig. S9.

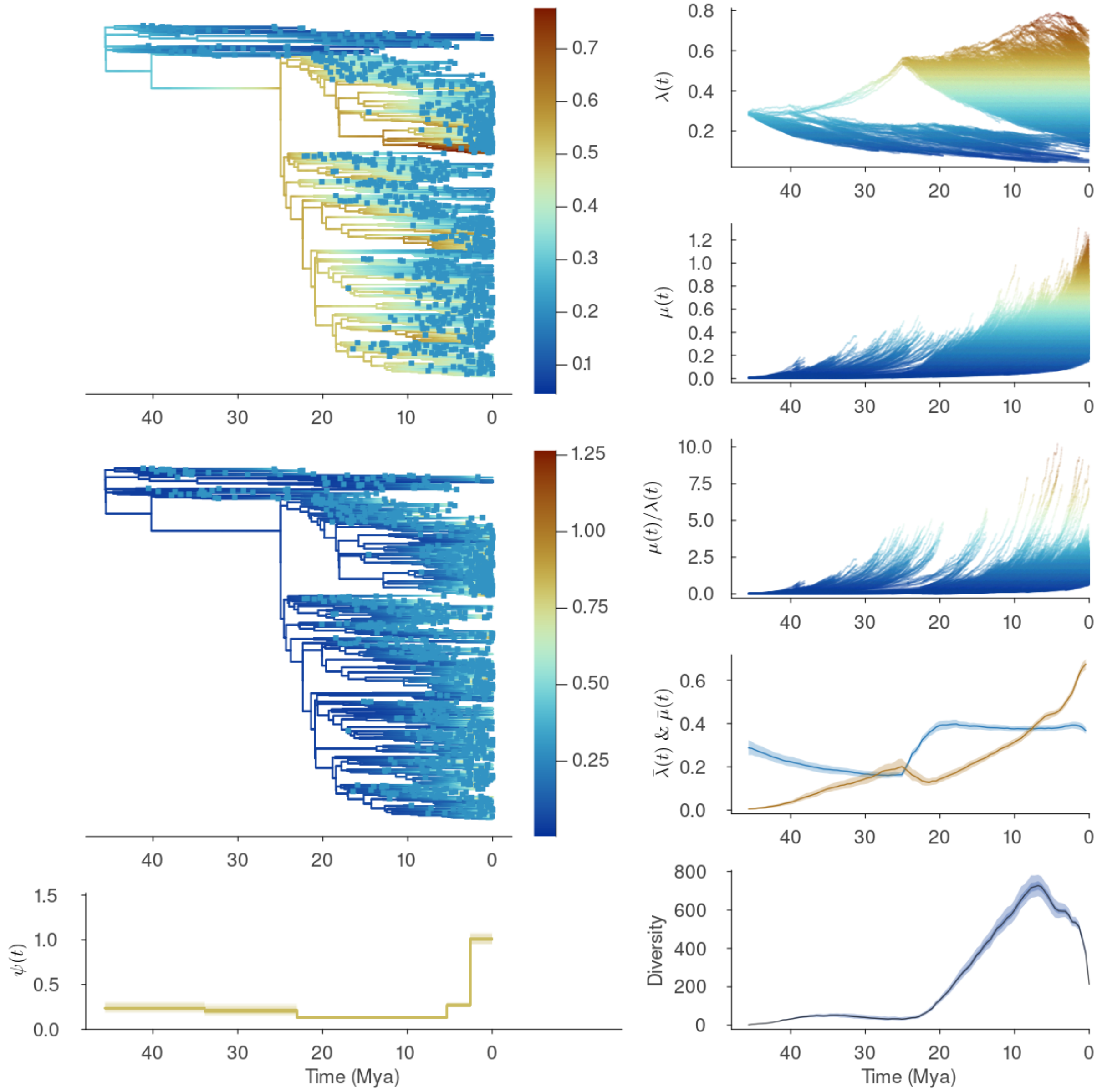


**Supplementary Figure 27: FBDD results for Ruminantia.** Details as in Fig. S9.

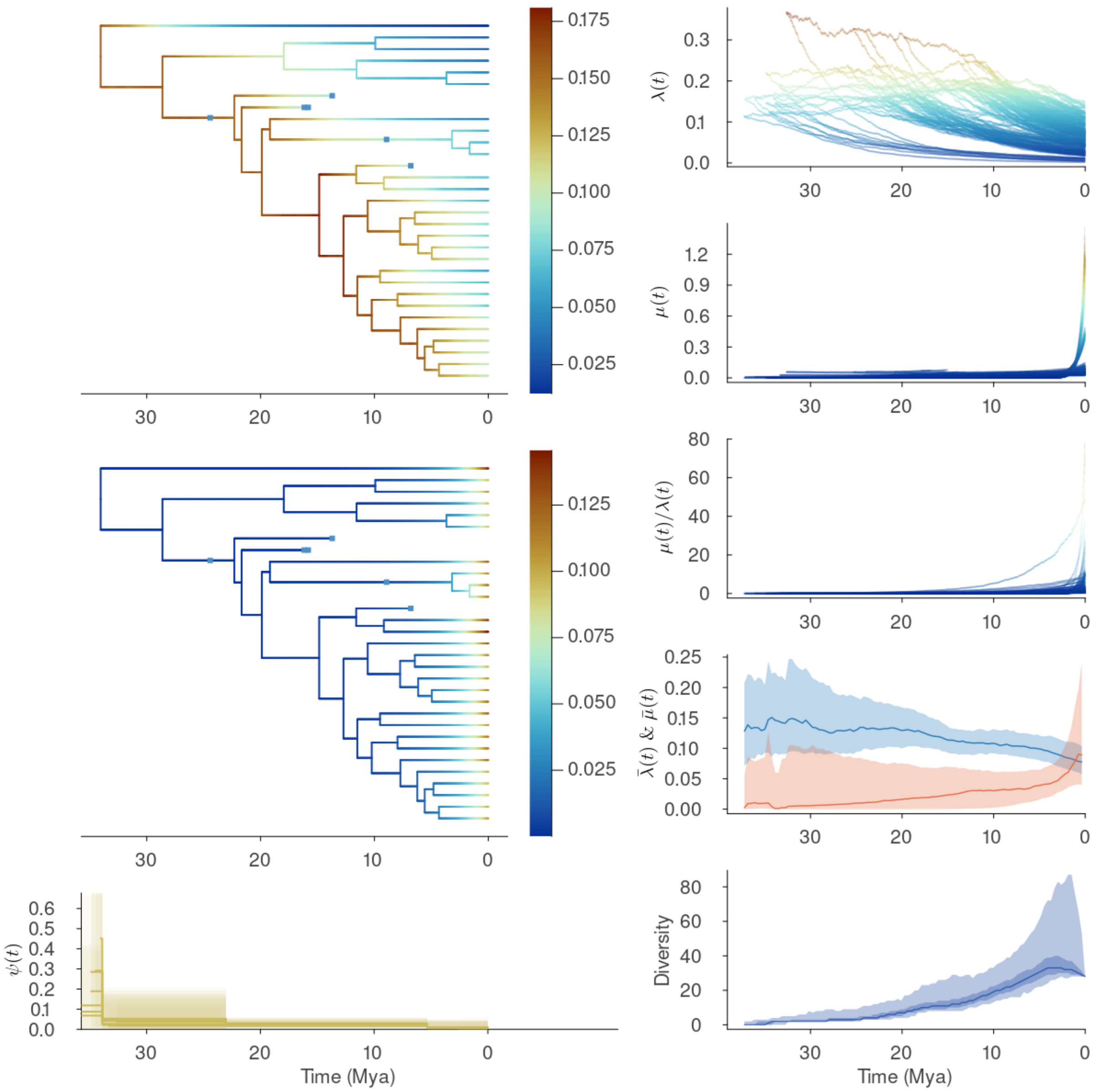


**Supplementary Figure 28: FBDD results for Sthenurinae.** Details as in Fig. S9.

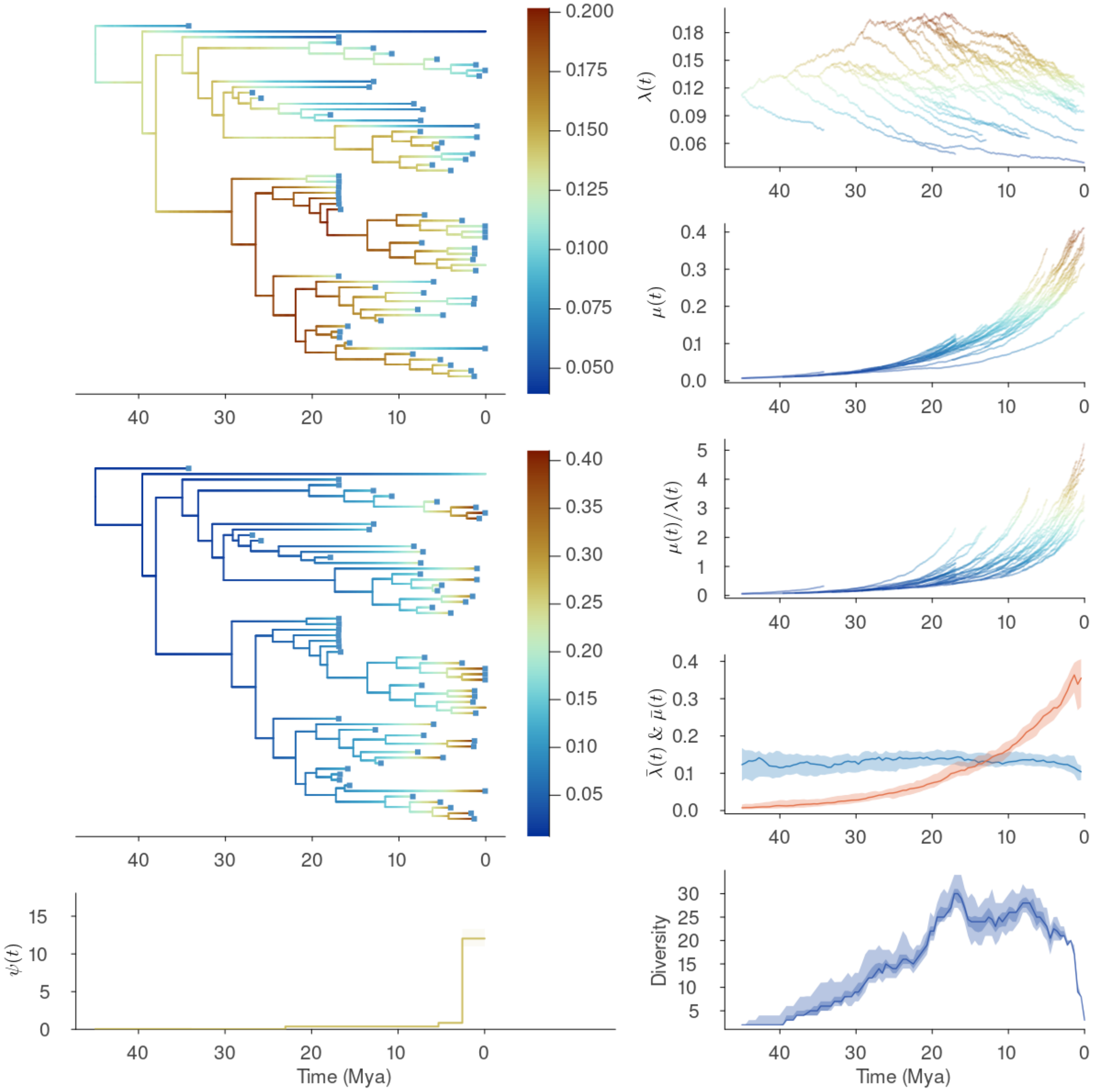


**Supplementary Figure 29: FBDD results for Folivora.** Details as in Fig. S9.

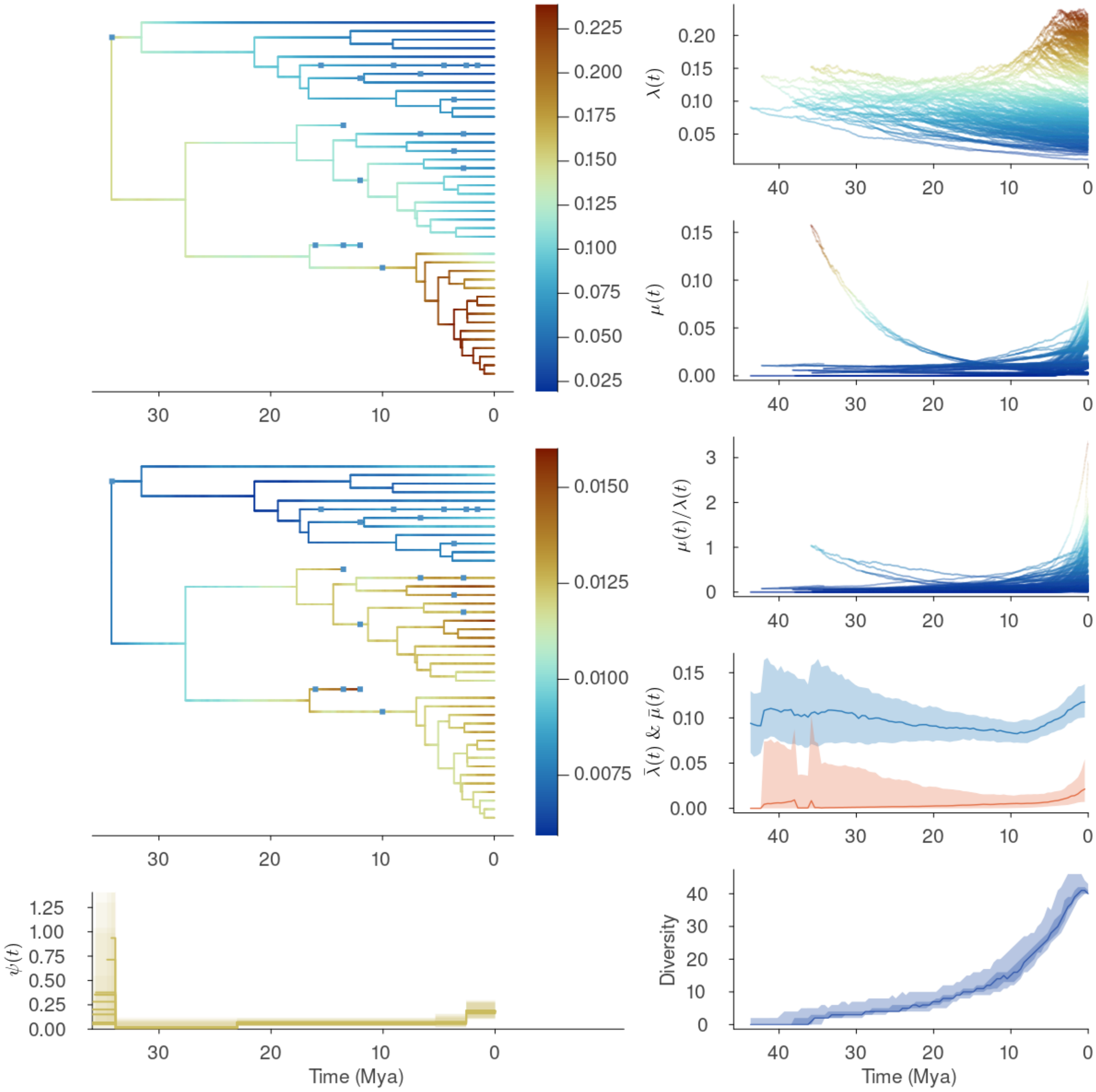


**Supplementary Figure 30: FBDD results for Centrarchidae.** Details as in Fig. S9.

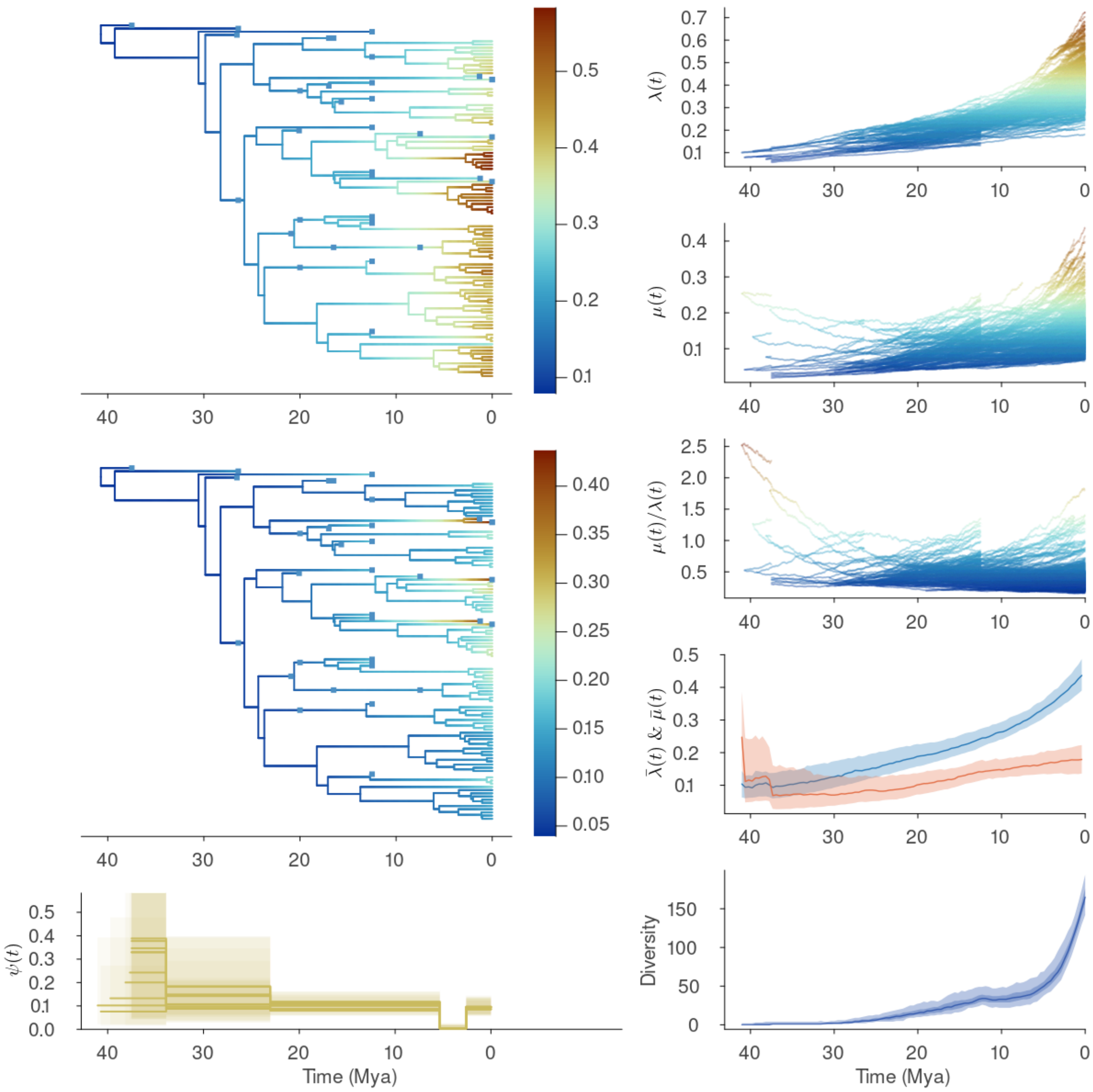


**Supplementary Figure 31: FBDD results for Platyrrhines.** Details as in Fig. S9.

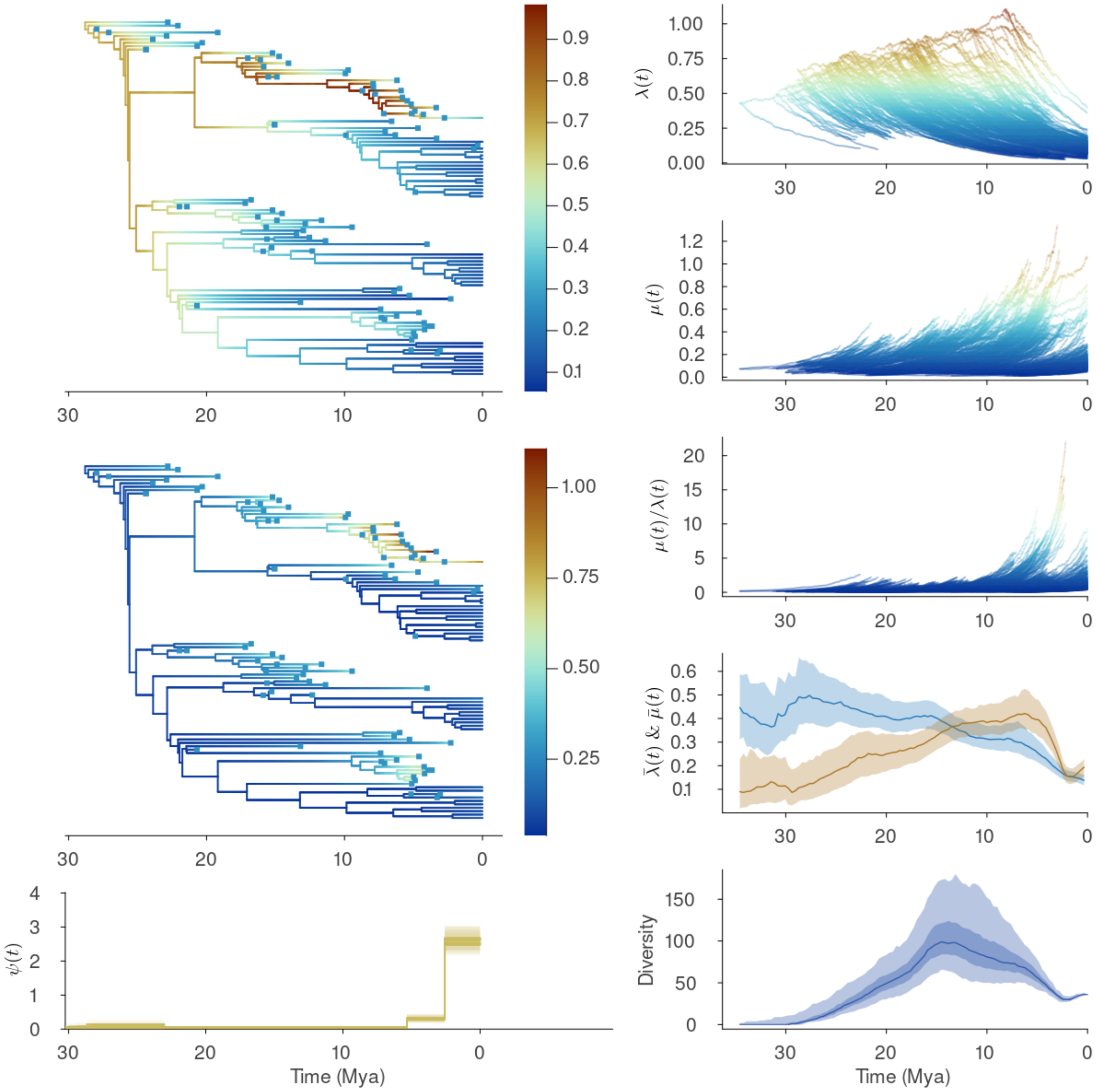


**Supplementary Figure 32: FBDD results for Pinnipedia.** Details as in Fig. S9.

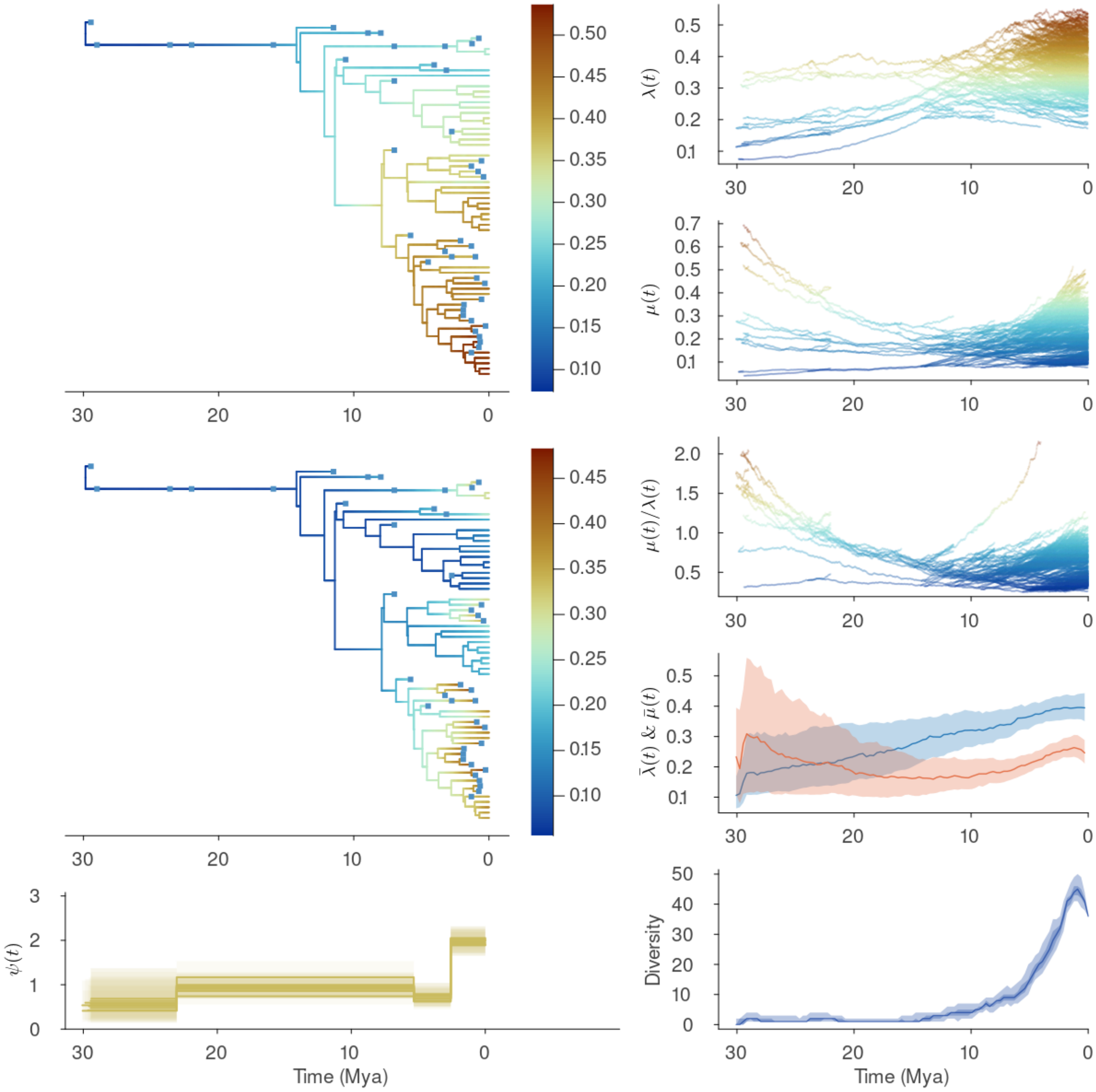


**Supplementary Figure 33: FBDD results for Caninae.** Details as in Fig. S9.

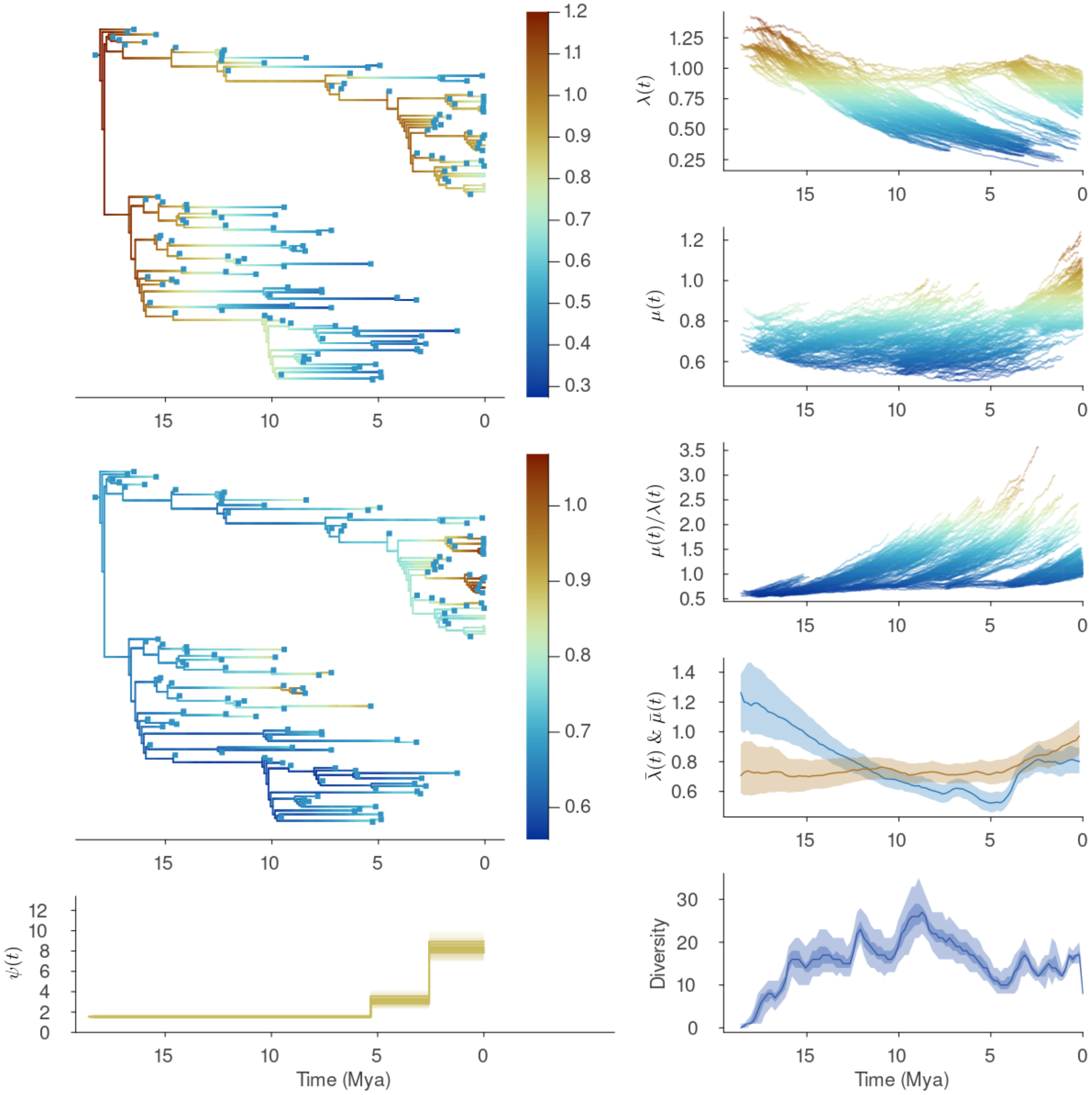


**Supplementary Figure 34: FBDD results for Equidae.** Details as in Fig. S9.

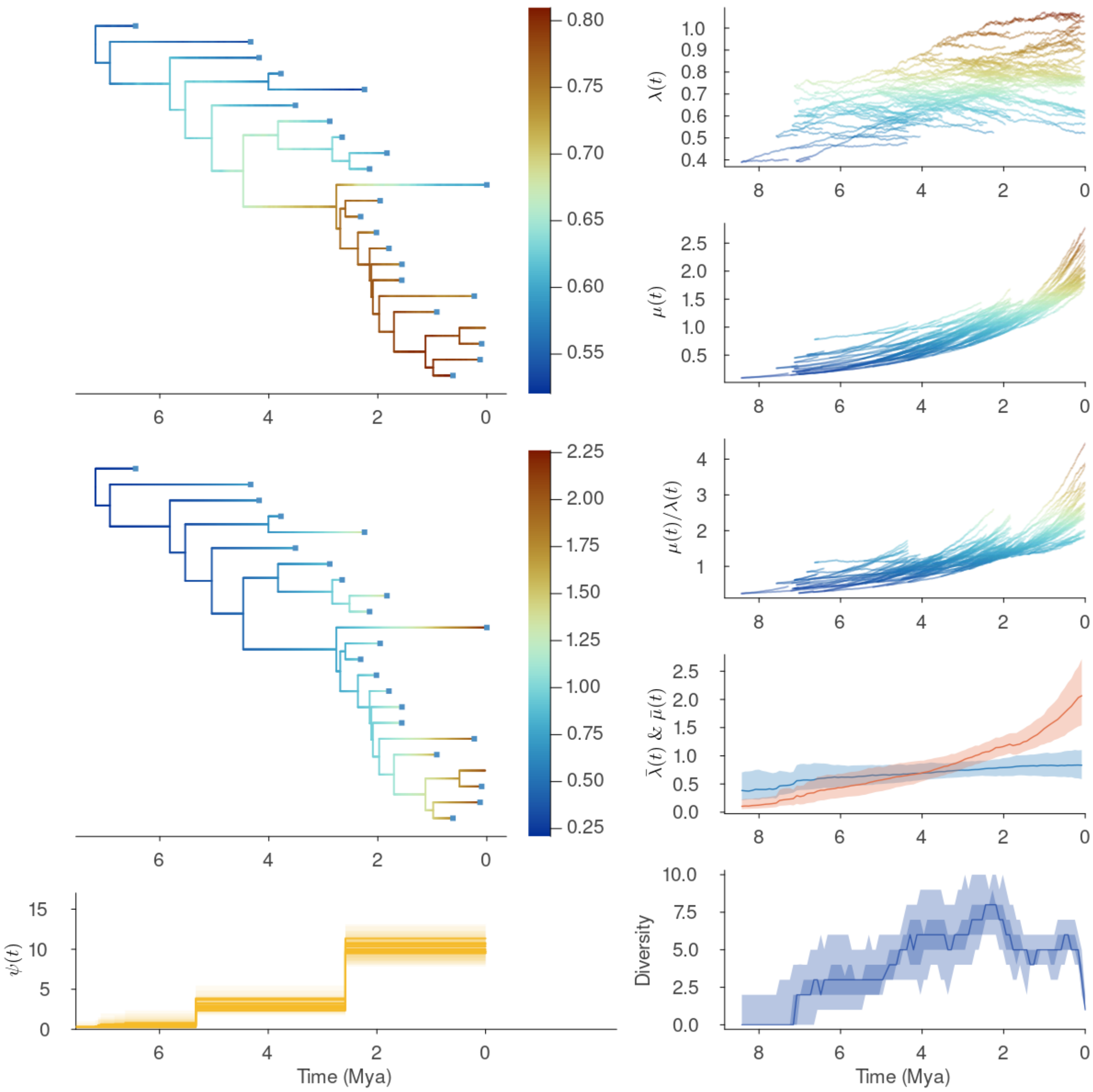


**Supplementary Figure 35: FBDD results for Hominini.** Details as in Fig. S9.

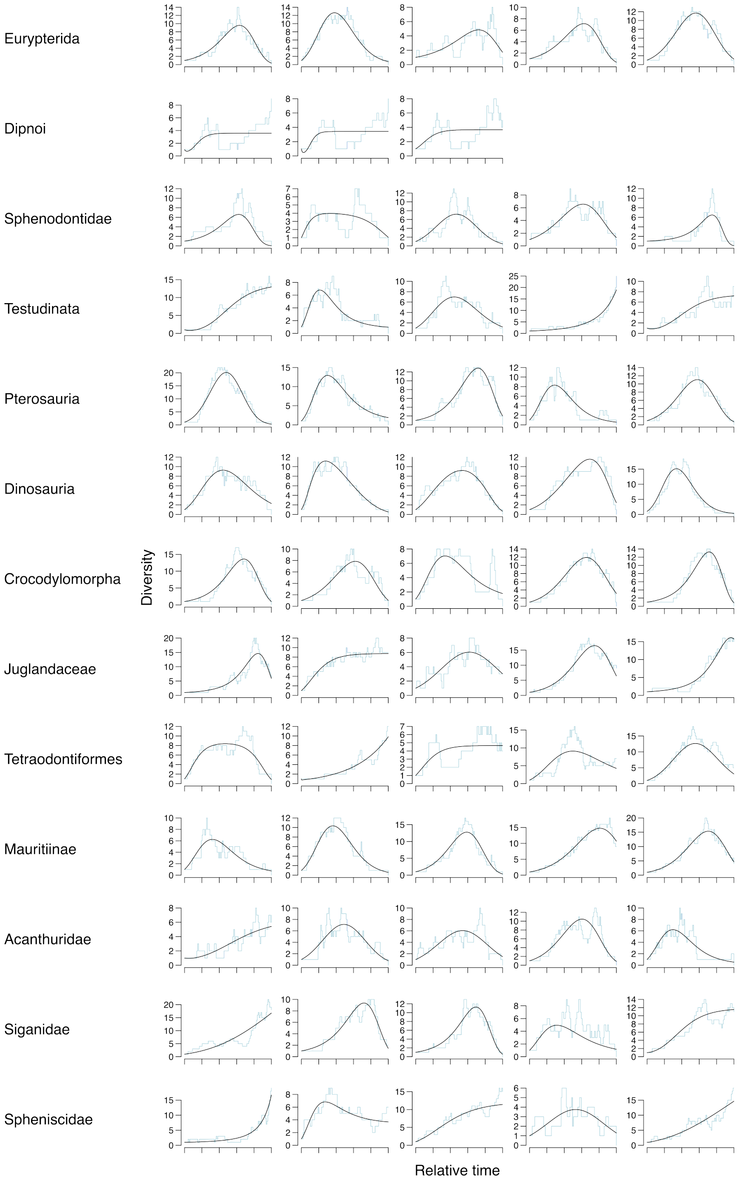


**Supplementary Figure 36: Sample of subclade diversity trajectories for the first 14 larger taxonomic groups.** Each row is a clade, and each column is a randomly sampled diversity trajectory from a subclade (holding 20 to 40 species). Times is standardized to 1 unit. In light blue we show the subclade diversity and in black we show the Expectation of the Negative Binomial regression (Materials and Methods) of the diversity trajectory. Note the great variation in diversity trajectories across subclades for a given taxonomic group.

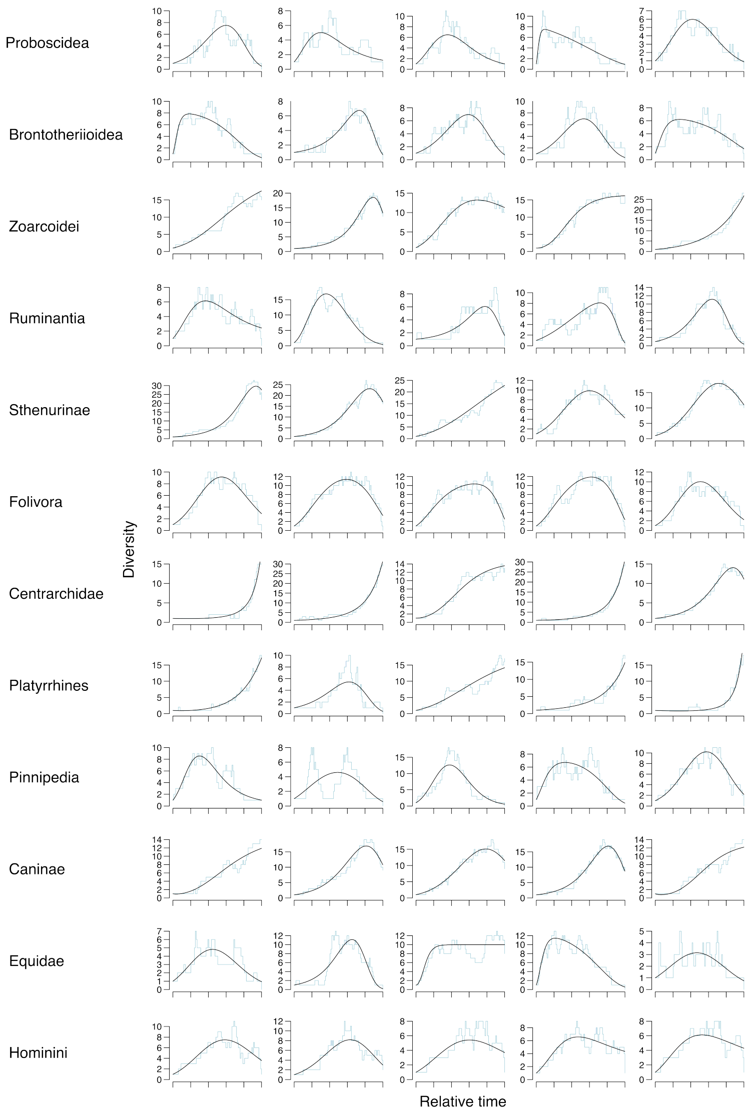


**Supplementary Figure 37: Sample of subclade diversity trajectories for the second 13 larger clades.** Details as in Fig. S36 but for the remaining 13 clades.

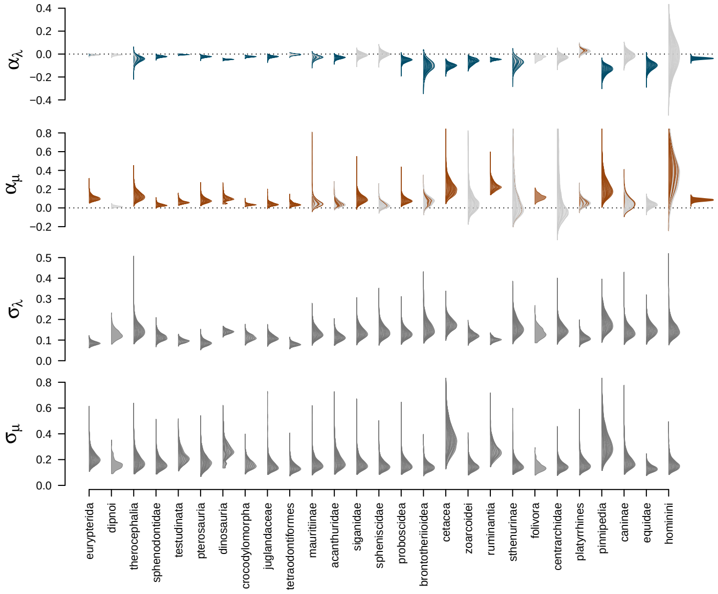


**Supplementary Figure 38: Posterior distributions for the governing parameters from the FBDD model across evolutionary radiations.** Posterior distributions for each sample, with their density scaled for visual purposes, stacked over the set of empirical trees for each taxonomic group for, from top to bottom, $\alpha_\lambda$, $\alpha_\mu$, $\sigma_\lambda$, $\sigma_\mu$. For $\alpha_\lambda$ and $\alpha_\mu$, the rightmost distribution displays the overall posterior distribution given by the hierarchical cross-taxa model used to plot Fig 4 (Materials and Methods).
